# Genomic Architecture of the *Resistance to Phytophthora Cactorum 2* (*RPc2*) Locus in Strawberry (*Fragaria* × *ananassa*)

**DOI:** 10.1101/2024.11.01.621609

**Authors:** Hyeondae Han, Youngjae Oh, Yoon Jeong Jang, Saket Chandra, Upinder Gill, Sujeet Verma, Young-Hee Noh, Sachiko Isobe, Dominique D. A. Pincot, Glenn S. Cole, Randi A. Famula, Mitchell J. Feldmann, Steven J. Knapp, Vance M. Whitaker, Seonghee Lee

**Author notes:** Correspondence: Seonghee Lee. Authors equally contributed to the manuscript (Hyeondae Han, Youngjae Oh).

## Abstract

Phytophthora crown rot (PhCR), caused by *Phytophthora cactorum*, is a soilborne disease with broad impacts to cultivated strawberries in the United States and worldwide. While a resistance locus, *RPc2*, has been identified in octoploid strawberries, the underlying genomic architecture and mechanism remain unclear. Here, we fine-mapped the *RPc2* region to 546 kb, containing 92 genes, and constructed a chromosome-scale haplotype-phased genome of resistant breeding selection FL 16.33-8. Comparative genome analyses with high-quality octoploid reference genomes, ‘Florida Brilliance’ (FaFB1) and ‘Royal Royce’ (FaRR1), identified two candidate genes, *Wall Associated Kinase 1* (*WAK1*) and *Cyclic Nucleotide Gated Channel 1* (*CNGC1*). Gene functions were validated using an Agrobacterium-mediated transient expression assay. Furthermore, by leveraging the population structure of the *RPc2* locus among genetically diverse breeding populations, we unveiled the complexity of genetic architecture and recent selective sweeps associated with *RPc2*. Notably, the predominant resistant haplotype (*RPc2*-H3) is prevalent in most commercial strawberry varieties in the US. The findings from this study are facilitating the advancement of genome-assisted breeding strategies for resistance to PhCR in strawberries.

**Summary:**

- Phytophthora crown rot (PhCR), caused by the soilborne pathogen *Phytophthora cactorum*, is a disease with broad impacts to cultivated strawberries worldwide. The *Resistance to Phytophthora Cactorum 2* (*RPc2*) locus is widespread in important cultivars and is a target of selection in breeding programs. Despite its importance, the genomic architecture and mechanisms of *RPc2* remain poorly characterized.
- We performed fine-mapping and delimited the genomic region to a 546 kb segment containing 92 genes. We developed a chromosome-scale haplotype-phased genome from an elite resistant line, FL16.33-8, and made comparisons to reference genomes of ‘Florida Brilliance’ (FaFB1) and ‘Royal Royce’ (FaRR1) to uncover gene sequence and structural variants in resistant accessions.
- Through comparative genomics and transcriptome profiling, we identified three candidate genes, *Wall Associated Kinase 1* (*WAK1*) and *Cyclic Nucleotide Gated Channel 1* (*CNGC1*) and *CNGC2*. The functions of these genes were validated using a simplified *Agrobacterium*-mediated transformation method for transient gene expression in strawberry crowns and roots.
- An examination of the population structure of the *RPc2* locus among genetically diverse accessions (N=1029) revealed that the predominant *H3* resistant haplotype is widespread in breeding accessions in the US, supporting the findings of a previous study on recent selective sweeps associated with *RPc2*. Results from this study are advancing genome-assisted breeding strategies for enhancing resistance to PhCR in strawberry.

## Introduction

Phytophthora crown rot disease (PhCR) of cultivated strawberry (*Fragaria* ×*ananassa*) is caused by *Phytophthora cactorum*, a hemi-biotrophic, oomycete plant pathogen. PhCR is considered one of the most significant soilborne diseases affecting strawberries in the temperate regions of the world and causes production losses of up to 40% (Stensvand *et al*., 1999). Soil fumigations using methyl bromide, dazomet, and chloropicrin 1,3-dichloropropene are effective ways to control *P. cactorum* (Cal *et al*., 2004). However, increasing limitations on the use of fumigants have led to deterioration in the control of *P*. *cactorum* in recent years (Cal *et al*., 2004; Holmes *et al*., 2020). The development of resistant cultivars has become important for strawberry breeding programs and a critical strategy for combating PhCR in an era where fumigation alternatives are decreasing while global consumption of strawberries continues to increase.

The resistance locus *Phytophthora cactorum 1* (*RPc-1*) was initially reported in *F. vesca* (Davik *et al*., 2015) but has not been found in cultivated strawberry. In a study conducted at UF, *FaRPc2* (hereafter referred to as *RPc2*) on chromosome 7B was identified using a pedigree-based analysis in complex and multiparent population sets (Mangandi *et al*., 2017). It was described that two primary haplotypes, *RPc2-H2* and *RPc2-H3* (hereafter referred to as *H2* and *H3*) were associated with resistance against *P. cactorum*. Each resistant haplotype explained about 40% of the phenotypic variance in the tested populations (Mangandi *et al*., 2017). These two resistant haplotypes seem to be originated from different sources and may represent distinct functional alleles. While all varieties with the H3 haplotype are resistant to PhCR, some susceptible varieties contain the H2 haplotype. This suggests that within the H2 haplotype, there may be very small allelic changes or sequence variants responsible for PhCR resistance. Although the exact cause is unknown, future studies should be conducted to elucidate the genomic region of *RPc2-H2*. he *H2* haplotype exhibits strong resistance to PhCR but is more poorly characterized.

Recently, Jiménez *et al*. (Jiménez *et al*., 2023) reported minimal genetic gains in breeding for Phytophthora crown rot (PhCR) resistance, with only 3% of cultivars exhibiting high resistance in a genetically diverse training population. However, through multivariate genome-wide association studies (GWAS), multiple significant loci contributing to PhCR resistance were uncovered, notably including the *RPc2* locus. Also, the observed differences in phenotypic and breeding values between UCD and non-UCD individuals in the training population align with insights from previous genome-wide studies, indicating a decrease in genetic variation in the UCD population due to strong directional selection, breeding bottlenecks, and selective sweeps (Jiménez *et al*., 2023). The result from this study also suggests a decline in resistance to PhCR in the UCD population over the last half-century, with non-UCD individuals harboring both favorable and unfavorable alleles not present in the UCD population or commonly found in gene banks (Jiménez *et al*., 2023). Following the searching of the genomic region near the SNP marker AX-184109190 associated with *RPc2*, (Jiménez *et al*., 2023) described several potential defense related candidate genes for *RPc2* In another study from a European breeding group, three resistance loci, *RPc6C, RPc6D,* and *RPc7D*, were identified using a bi-parental population (‘Emily’ × ‘Fenella’) (Nellist *et al*., 2019). The *RPc7D* locus was shown to locate in the same genomic region as *RPc2*, suggesting that these are same QTL.

The cultivated strawberry has a complex allo-octoploid genome (2n = 8x = 56), consisting of four subgenomes with total genome size of 850 Mb (Hirakawa *et al*., 2014; Hummer *et al*., 2011; Tennessen *et al*., 2014). The first chromosome-scale reference genome was assembled from ‘Camarosa’(Edger *et al*., 2019), and subsequent genomes were published from the Japanese cultivar ‘Reikou’ (Shirasawa *et al*., 2021) and a highly homozygous inbred line ‘Wongyo3115’ (Lee *et al*., 2021). Recently, two fully haplotype-phased genomes were published from a day-neutral cultivar, ‘Royal Royce’ (FaRR1) and short-day cultivar ‘Florida Brilliance’ (FaFB1). ‘Florida Brilliance’ is susceptible to PhCR, and the FaFB1 genome does not possess the resistant haplotype of *RPc2*. It has been reported that the cultivar ‘Royal Royce’ is moderately resistant to Verticillium wilt (*Verticillium dahlia*), moderate susceptibility to Phytophthora crown rot (*Phytophthora cactorum*) and Fusarium wilt (*Fusarium oxysporum*), and susceptible to Charcoal rot (*Macrophomina phaseolina*) (https://itc.ucdavis.edu/wp-content/uploads/UCD-Royal-Royce-Cultivar.pdf). To determine the genomic architecture of *RPc2*, it will be necessary to construct a chromosome-scale haplotype-phased genome for a resistant cultivar.

The molecular mechanisms underlying *RPc2*-mediated resistance against *P. cactorum* are largely unexplored. Although a transcriptome analysis has been conducted in diploid strawberry (*F. vesca*) challenged with *P. cactorum*, it is important to note that the regulation of defense responses during the pathogen infection in octoploid strawberry appear to differ from that in *F. vesca* due to diverse gene expression patterns across homoeologous copies (Toljamo *et al*., 2016). Studies have highlighted the importance of genes encoding cell wall strengthening proteins, particularly those involved in lignin biosynthesis enzymes associated with Damage-Associated Molecular Patterns (DAMPs)-mediated defense pathways, in responding to *Phytophthora* infection in plants. Additionally, there have been reports of the differential expression of plant surface receptors, including 25 receptor-like proteins and 224 receptor-like kinases, in tomato plants infected with *Phytophthora cactorum* (Zhou *et al*., 2023). Jiménez *et al*. (2023) described that the *RPc2* locus spanning 1.26 Mb contained 233 annotated genes with 18 of them associated with plant defense responses. Notably, these genes include intracellular leucine-rich repeat (LRR)-type immune receptors, membrane-localized immune receptors, Ca2+-dependent protein kinases, and cyclic-nucleotide-gated calcium channels (*CNGCs*), known to play important roles in plant disease resistance. Moreover, a gene homologous to *WAK1*, encoding a wall-associated receptor kinase galacturonan-binding protein, was highlighted as a promising candidate (Jiménez *et al*., 2023).

Despite the abundance of genetic information available for *RPc2*, there is currently no study specifically characterizing causal variants, sequence structural variants, or testing of gene function for this locus. Exploring these aspects could offer valuable insights into mechanisms of resistance and facilitate the improvement of resistance to PhCR through genome-assisted breeding approaches. In this study, we finely mapped the genomic region for *RPc2* using a diverse mapping panel. A haplotype-phased genome of an elite resistant accession (FL 16.33-8) was developed in order to characterize the resistant haplotypes and comparatively analyzed with other publicly available high-quality haplotype-phased genomes of octoploid strawberry. Furthermore, a genome-wide transcriptome profiling analysis highlighted two candidate genes, *Wall Associated Kinase 1* (*WAK1*) and *Cyclic Nucleotide Gated Channel 1* (*CNGC1*), that appear to play pivotal roles in regulating resistance to PhCR in strawberry. Using an efficient *Agrobacterium*-mediated transient gene expression method, we successfully validated the functions of these candidates in the crowns and roots of strawberry plants. To understand population genetic structure of the *RPc2* locus, we conducted a survey of globally diverse breeding accessions (N=1029), revealing recent positive selection of the *H3* haplotype.

## Materials and Methods

### Genome sequencing and assembly

An elite resistant breeding selection FL16.33-8, which contains the major resistant haplotype *RPc2*-*H3* was identified for genome assembly. FL16.33-8 originated from a 2016 cross between unreleased selection FL13.27-142 and FL12.93-4. FL13.27-142 was chosen as a resistant source of *H3*. Etiolated young white leaf tissues were used for extracting high molecular weight DNA. The single-molecule real-time sequencing (SMRT) bell library construction and Hi-Fi long read PacBio DNA sequencing were conducted at DNA Link (Seoul, South Korea). The genome of FL 16.33-8 was assembled using the Hifiasm (Cheng *et al*., 2021), which is a de novo assembler that utilizes long high-fidelity sequence reads to represent the haplotype information in a phased assembly graph. Two parents’ Illumina paired-end (PE) data was utilized to phase haplotype scaffolds. The genome continuity of the Illumina draft genome assemblies was assessed with QUAST (Gurevich *et al*., 2013). A BUSCO analysis was performed using default parameters to assess the core gene set (Simão *et al*., 2015). The de novo genome assembly, raw genomic reads were used as input data for Ragtag (Alonge *et al*., 2022), a tool designed for contig-ordering and scaffolding genome assemblies. Genome quality and completeness was assessed using Merqury ver 1.3 (Rhie *et al*., 2020), which evaluates assembly based on efficient *k*-mer set operations. The collinearity with the genetic linkage map for FL 16.33-8 and ‘Florida Brilliance’ was visually inspected using Circa (https://omgenomics.com/circa/). The LTR Assembly Index for each haplotype-phased genome was calculated based on the custom repeat library (Ou *et al*., 2018).

To identify and classify repetitive elements in the genome, LTR retrotransposon candidates were searched using EDTA (Su *et al*., 2021). For the gene annotation, two haplotype-phased assembly was annotated using Genome Sequence Annotation Server (Humann *et al*., 2019). Two methodologies in parallel were used to mask and annotate the repeat library. A library of predicted repeats detected with RepeatModeler (version 2.0.1; www.repeatmasker.org), was combined with a library of repeats identified using RepeatMasker (version 4.1.1; www.repeatmasker.org) and an *Arabidopsis thaliana* repeat library within GenSAS to create a consensus library of repeats and mask polished assembly. Nucleotide sequences were aligned to NCBI refseq plant using BlastN ver. 2.7.1 (Camacho *et al*., 2009) as well as Diamond (Buchfink *et al*., 2015). These alignments were combined with results of several gene prediction modeling including AGUSTUS ver 3.3.1 (Hoff and Stanke, 2019) to generate an official gene set and identify predicted transcripts within each masked assembly using EVidenceModeler (Haas *et al*., 2008).

### Scaffold validation using a genetic linkage map

A high-density genetic map was developed using a total of 169 F_1_ individuals from a cross between ‘Florida Brilliance’ and breeding selection FL 16.33-8. Axiom™ IStraw35 SNP arrays were used to genotype all 169 F_1_ individuals. Markers were filtered to have approach was applied to identify singletons to fix the marker order and regions with low marker density or gaps caused by segregation distortion. A final linkage map consisting of 10,269 SNP markers was used to validate the scaffolds from the whole genome assembly. SNP probes sequences used for the construction of linkage maps were mapped to the FL 16.33-8 assembly sequence using blastn procedure. Alignments were filtered to retain markers if they matched to unique sequence position in the phased genome assembly and with a maximum of 2 mismatches in the second-best hit. The alignments were queried to detect potentially problematic scaffolds mapped with SNP probes from different LGs. The number of scaffolds with SNP probes mapped from different LGs was used as a metric in the quality assessment.

### Fine-mapping

A total of 339 accessions with three seasons field phenotypic data (2013-2015) were used for fine-mapping analysis (Supplementary Table S1). DNA of additional strawberry accessions was extracted by the simplified CTAB (cetyltrimethylammonium bromide) method with minor modifications (Keb-Llanes *et al*., 2002; Noh *et al*., 2017). To further fine map the *RPc2* region, we developed sub-genome specific high-resolution melting (HRM) markers by selecting FanaSNP array probes every 10 kb in the *RPc2* region, and selected HRM markers discriminating H2 and H3 alleles through HRM analysis. The HRM markers representing 15 of Axiom® IStraw90 SNP probes and 10 FanaSNP array probes encompassing *RPc2* region (Hardigan *et al*., 2020; Mangandi *et al*., 2017; Noh *et al*., 2018) were used to genotype 339 breeding accessions (Supplementary Table S2). Amplification products were resolved by high-resolution melting analysis in LightCycler® 480 system II (Roche Life Science, Germany). Melting curve data was analyzed by Melt Curve Genotyping and Gene Scanning software for Roche LightCycler 480 II system.

### RNA sequencing analysis

The plants were grown for four weeks in growth chamber under ideal conditions (73 ± 2°F, Relative humidity 57 - 63%, 16 h light and 8 h darkness). Total of four breeding accessions were used for RNA sequencing study: FL11.28-34 (H3:H1; heterozygous resistant haplotype), FL12.75-77 (H3:H3; homozygous resistant haplotype), FL12.82-44 (H1:H1; homozygous susceptible haplotype) and FL13.22-336 (H1:H1; homozygous susceptible haplotype). All accessions were highly connected and comprises two haplotypes (H1 and H3), as each of them has arose from elite breeding population and had undergone repeated selection for >15 generations (Mangandi *et al*., 2017). The pathogen inoculation and spore concentration were prepared according to the method as exactly mentioned in previous study (Mangandi *et al*., 2017). Crown tissues were collected at 0, 24-, 48-, 72-, and 96-hour post inoculation (hpi) with at least three biological replications.

For RNA extraction, crown tissues from each five time points were collected, and samples were ground into fine powder using liquid nitrogen in mortar and pestle. Total RNA was extracted according to the protocol of the Spectrum^TM^ Plant Total RNA Kit (Sigma-Aldrich, St. Louis, MO, United States). To remove any traces of DNA, the isolated RNA was treated with DNAse I (Invitrogen) and resuspended in a total volume of 50 μl of RNase-free water. In total of 1 μg total RNA was used for first-strand cDNA synthesis using Transcriptor First Strand cDNA synthesis kit (Roche, Switzerland) following the manufacturer’s instructions. The RNA samples of 0 and 96 hpi were selected for RNA-seq study. The paired-end library construction and sequencing was conducted on the Nextseq 500 Illumina sequencing platform (Novogene, USA). Illumina adapters were removed from the raw reads using Trimmomatic ver. 0.33 (Bolger *et al*., 2014). After quality assessment with FastQC (https://www.bioinformatics.babraham. ac.uk/projects/fastqc/), the filtered reads were then mapped to each haplotype-phased genome assembly using HISAT2 (Kim *et al*., 2019). The sequence alignment map (SAM) format was converted to Binary Alignment Map (BAM) format. To import the BAM files into R, ‘Rsamtools’ package was used. Differentially expressed genes in resistant vs susceptible were identified using the DESeq2 pipeline. DESeq2 software was employed to calculate the expression between genotypes (Love *et al*., 2014). Pairwise comparison was performed between *H1*-water control (WC) vs. *H1*-pathogen inoculation (PI), *H1*-PI vs. *H3*-PI, *H3*-WC vs. *H3*-PI, and total resistance *vs.* total susceptibility. The genes with a | log2 fold change| ≥ 1 and FDR-adjusted *p*-value (q-value) < 0.05 were selected as DEGs. Heatmap was generated using R package ‘gplots’ using expression data for genes locating at interval of *RPc2*. The differentially expressed sequences were annotated with the help of sequence homology search using blastx program in BLAST+ package (Camacho *et al*., 2009). The sequences were analyzed at an e-value of 1e^-6^ to NCBI non-redundant protein database. The homologues were selected on the basis of the e-value of less than 1e^-6^ and highest high-scoring segment pairs (HSP).

### Comparative genomic data analysis

For the comparative genomic analysis of the two different haplotypes of *H1* and *H3*, we selected the 546 kb *H3* region identified from fine-mapping. This region was extracted from the genomes of ‘Royal Royce’ (homozygous *H3*), ‘Florida Brilliance’ (homozygous *H1*), and the phase 1 and 2 assemblies of FL16.33-8 (*H3*), respectively. Genomic sequences were compared with Mauve software in the Geneious Prime software 2020.2.4. The nucleotide and amino acid sequences of candidate genes, *WAK1*, *CNGC1*, and *CNGC2*, were compared using MAFFT alignment in Geneious Prime software version 2020.2.4.

### *Agrobacterium*-based transient assay

To validate candidate gene function in strawberries, crown and root transient overexpression and RNAi transient assays were performed using *A. tumefaciens*, with modifications to the previous method (Zhong *et al*., 2016). We used three-week-old strawberry daughter plants of ‘Florida Brilliance’ (susceptible) for overexpression transient assays and ‘Florida Beauty’ (moderately resistant) for RNAi transient assays. The pMDC32 and pK7GWIWG(II) gateway vector systems were used for overexpression and RNAi transient assays (Curtis and Grossniklaus, 2003; Karimi *et al*., 2002). The final cloning products of pMDC32::*WAK1*, pMDC32::*CNGC1*, pK7::*WAK1*, and pk7::*CNGC1/CNGC2* were introduced *A. tumefaciens* EHA105.

Transformed *A. tumefaciens* were grown at 28℃ overnight in LB medium containing antibiotics appropriate for each vector. When the culture density reached to OD_600_ of 1.0, *A. tumefaciens* were harvested by centrifuge at 4500rpm for 25min and resuspended in adjusted to an OD_600_ of 0.6 in MMA activation buffer (10mM 2-N-morpholino) ethanesulphonic acid (MES), 10mM MgCl_2,_ pH5.6 and 200 μM acetosyringone) and shaken 50-70rpm for 3hr at room temperature before infiltration of root and crown. To enhance the transformation efficiency, we applied the vacuum infiltration method during the overexpression and RNAi transient assays. Briefly, roots and crowns were gently scraped with a syringe needle before vacuum infiltration, then transferred to a suspension *A. tumefaciens* with 0.05% silwet L-77. Vacuum infiltration was conducted for five minutes. Subsequently, the inoculated plants were soaked in the same solution for three hours at room temperature. Afterward, the plants were transferred to 4-inch pots in a humid box in a growth room for four days (12/12hr light/night, 23-25°C). Four days later, the pathogen inoculum (zoospore suspension with a final concentration of 1 × 10^4^ zoospores/ml) was prepared according to the method in previous study (Mangandi *et al*., 2017). Inoculations were performed by immersing the plant roots for five seconds in 500 ml of the zoospore suspension up to the base of the crown immediately before transferring to pots. The RNAi and overexpression experiments were conducted twice over time. Each experiment was conducted with three biological replicates with empty vector controls. After 17 days post-infection, plant crowns were halved, and photos were taken from each plant. The percentage of symptomatic areas was assessed using ImageJ software to quantify the areas showing typical *P*. *cactorum* symptoms on the crown and roots, including the areas of halved crowns and the internal symptomatic areas. Subsequent statistical analysis was performed using R software, conducting One-way ANOVA, and Tukey’s honestly significant difference (HSD) test was used to compare the means of all treatments to each other and statistically validate the data (*p* ≤ 0.05). The target genes were *FaWAK* and *FaCNGC*, while *FaGAPDH2* was selected as housekeeping gene control. Primers were designed using Primer 3 (https://primer3.ut.ee/) tool, and all primer sequences were represented in Supplementary Table S3. The qRT-PCR experiment was performed using LightCycler® 480 system (Roche Diagnostics, Mannheim, Germany) using Forget-Me-Not™ EvaGreen® qPCR Master Mix (Biotium, CA, USA).

### Population structure analysis and phylogenetic analysis

The 1,029 octoploid accessions (Supplementary Dataset2 and 3). A total of 17 SNP markers in the 546 kb of *RPc2* region genotype matrices were used to evaluate population structure by principal component analysis and clustering with an admixture model using STRUCTURE (v.2.3.4) (Pritchard *et al*., 2000). The STRUCTURE analysis tested for K=2-10 subpopulations (10,000 burn-in steps and 10,000 Markov-Chain Monte Carlo steps with 10 replicates per K value). The optimal subpopulation (K) value was determined using STRUCTURE HARVESTER (v0.6.94) (Earl and VonHoldt, 2012). The results were visualized using pophelpher (Francis, 2017). Principal component analysis was conducted using TASSEL5 (Bradbury *et al*., 2007). Cladogram was generated using fastTree (Price *et al*., 2009) on Geneious Prime. After estimating genetic distance matrices were generated, a maximum-likelihood phylogenetic tree was constructed.

## Results

### Fine-Mapping of *RPc2* Conferring Resistance to Phytophthora Crown Rot in Octoploid Strawberry

In our previous investigation, we mapped *RPc2* to a 1.12 Mb segment using 15 SNP markers derived from Axiom® IStraw90 SNP probes (Noh *et al*., 2018) (Fig. 1A, Supplementary Tables S1 and S2). To refine this interval for effective marker-assisted selection in resistance breeding, we assembled an association mapping panel consisting of 339 accessions. This panel comprised 206 accessions used in the previous study for QTL analysis (Noh *et al*., 2018) and an additional 133 accessions selected based on consistent field data for mortality due to PhCR collected over three seasons (Supplementary Table S1). Employing these association mapping panels, family-based linkage studies were performed to identify within-family recombination events. To further enhance the resolution of the identified interval, 15 sub-genome-specific SNP or InDel-based high-resolution melt (HRM) markers were employed, including four newly developed markers using the 50K FanaSNP Array (Hardigan *et al*., 2021) (Fig. 1A, Supplementary Table S1). The genotyping of all 339 accessions with these 15 HRM markers revealed recombination events at markers AX-89845076 and AX-184173618, resulting in a refined *RPc2* region reduced from 1.12 Mb to 546 kb (Fig. 1A, Supplementary Table S1). The markers between AX-18423389 and AX-184173618 did not accurately predict the phenotype of some individuals containing the H2 haplotype.

**Fig. 1.**
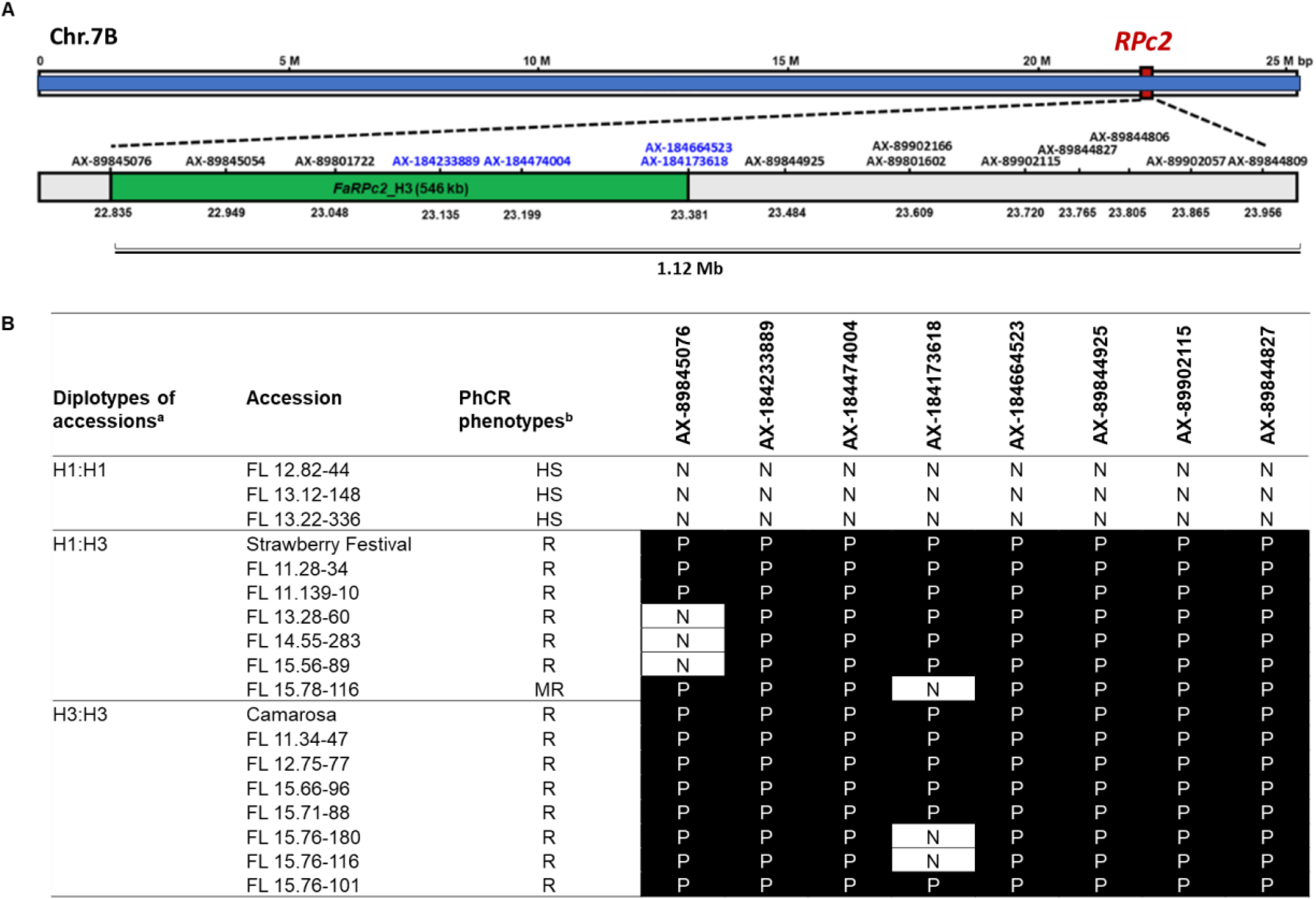
Fine-mapping of the *RPc2* genomic region conferring resistance to Phytophthora crown rot resistance. (A) Preliminary mapping of *RPc2* region spans approximately 1.12 Mb and is based on 11 markers (black) derived from Axiom® IStraw90 SNP probes (Mangandi et al., 2017). Using the additional subgenome-specific HRM markers (blue) derived from FanaSNP 50k array, *RPc2* was delimited to 546 Kb. (B) HRM marker genotype results (N: negative, P: positive) associated with the resistance haplotype, *RPc2*-*H3*; (a) showcases the diplotypes of accessions in the *RPc2* region from Mangandi et al. (2017). *H1* and *H2*: susceptible haplotype, *H3*: resistance haplotype (b) is based on the area under the disease progress curve (AUDPC) values reported in Mangandi et al. (2017), with R: resistance, MR: moderate resistance, HS: high susceptibility, S: susceptibility.

### Haplotype-Phased Genome Assembly and Annotation of Resistant Breeding Selection ‘FL 16.33-8’

The trio-binning method was applied to develop a high-quality haplotype-phased genome of the resistant accession FL 16.33-8 using PacBio HiFi long reads and parental short reads. The contig N50 values for FL 16.33-8-Phase1 and FL 16.33-8-Phase2 were 16.4 Mb and 14.3 Mb (Table 1). The assembly spans a total length of 782.4 Mb for FL 16.33-8-Phase1 and 778.1 Mb in FL 16.33-8-Phase2, distributed across 56 sub-chromosomal pseudomolecules (Supplementary Table S4). These pseudomolecules were numbered following the convention of reference genomes from ‘Camarosa’, ‘Royal Royce’-FaRR1, and ‘Florida Brilliance’ - FaFB1 (Fig. 2A, Supplementary Table S4). In FL 16.33-8-Phase1, the shortest and longest chromosomes were chromosome 7D (19.35 Mb) and chromosome 6B (43.16 Mb), while in FL 16.33-8-Phase2, they were chromosome 7C (21.75Mb) and chromosome 6C (34.25 Mb) (Supplementary Table S4). Genome assembly and annotation completeness showed 98.2% of complete BUSCO in both FL 16.33-8-Phase1 and FL 16.33-8-Phase2, indicating a high-quality chromosome-scale assembly (Supplementary Table S5).

**Fig. 2.**
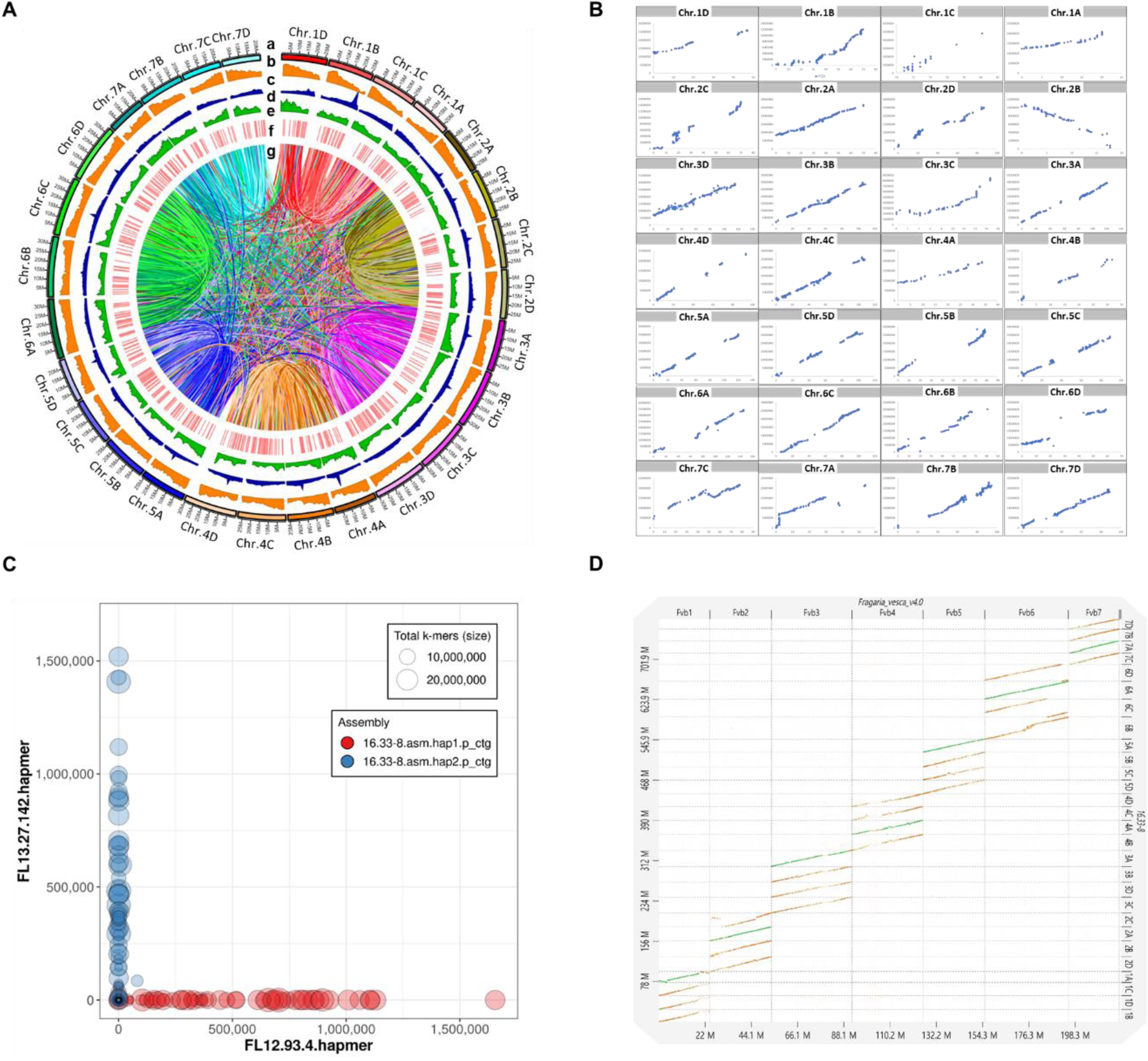
The haplotype-phased genome assembly of an elite resistant line, FL 16.33-8 containing the resistance haplotype *H3*. (A) Genome visualization of haplotype-phased genome A of FL 16.33-8. The tracks from outside to inside are: a = haplotype-phased subgenome name, b = length (Mb) of sub-genome, c = number of genes per 1 Mb window, d = density of long term repeated (LTR) per 1 Mb window, e = LTR assembly index (LAI) in sub-genomes from 1A to 7D plotted in 1 Mb window, f = The indel position differing by more than 30,000 bp among ‘Royal Royce’, ‘Wonkyo 3115’, and FL 16.33-8, g = synteny comparison within the haplotype-phased genome 1. (B) Comparison of the collinearity between a genetic linkage map and haplotype-phased genome 1 of FL 16.33-8. Genetic positions from a linkage map for ‘Florida Brilliance’ × FL 16.33-8 were plotted against the corresponding physical positions in the ‘Florida Brilliance’ genome assembly, FaFB1 (blue points). The X-axis indicates the genetic linkage map in centimorgans (cM), while the Y-axis represents genome coordinates in the haplotype-phased genome A of FL 16.33-8. (C) The k-mer blob Plot illustrates the distribution of maternal (red) and paternal (blue) k-mer within each scaffold of the respective assemblies. The size of the blob corresponds to the total number of k-mers in that scaffold. (D) Dotplot of FL 16.33-8-phase1 assembly against diploid *Fragaria vesca* ver. 4.0. Dot plots are produced using the DGENIE software and alignments with minimap2.

**Table 1.** Statistics on assembled haplotype-phased genome of a resistant accession, FL 16.33-8.

| Metrics | Value |  |
| --- | --- | --- |
|  | Phase 1 | Phase 2 |
| Number of contigs ( $\geq 10,000$ bp ) | 1,468 | 592 |
| Number of contigs ( $\geq 25,000$ bp ) | 1,447 | 587 |
| Number of contigs ( $\geq 50,000$ bp ) | 637 | 456 |
| Total length ( $\geq 10,000$ bp) | 869,351,047 | 826,921,887 |
| Total length ( $\geq 25,000$ bp) | 868,872,493 | 826,804,304 |
| Total length ( $\geq 50,000$ bp) | 837,337,314 | 821,510,393 |
| Number of contigs | 1,468 | 592 |
| Largest contig | 27,317,239 | 27,253,556 |
| Total length | 869,351,047 | 826,921,887 |
| GC (%) | 40.1 | 39.99 |
| N50 | 16,418,230 | 14,284,693 |
| N75 | 10,170,993 | 9,465,336 |
| L50 | 23 | 22 |
| L75 | 39 | 40 |
| Number of N's per 100 kbp | 0 | 0 |

To validate the accuracy of haplotype phasing in the genome assembly, we employed a high-density genetic linkage map developed from a cross of ‘Florida Brilliance’ by ‘FL 16.33-8’. The genetic map consisted of 13,910 SNP markers at 10,269 distinct positions across 31 linkage groups (LGs) with a total genetic length of 2,382.03 cM (Supplementary Table S6; Supplementary Dataset 1). The 31 LGs from LG1A to LG7D showed synteny with one of the 28 sub-chromosomal pseudomolecules from the assembly with less than 0.5% of markers exhibiting conflicts with their positions in the assembled chromosomes (Fig. 2B, Supplementary Table S6). Sub-genome specific FanaSNP markers used in constructing the linkage map were anchored to sub-chromosomal pseudomolecules of the reference genome and assigned to each subgenome from chromosome 1-1 or 1A to chromosome 7-1 or 7D based on nomenclature of ‘Camarosa’ or ‘FaRR1’ reference genomes (Fig. 2B). Trio binning with hifiasm (v0.12) successfully classified both haplotypes of FL 16.33-8, resulting in *k*-mer distributions for the read bins and an assembly that fully resolved both parental haplotypes (Fig. 2C). This observation was further confirmed by the spectra plot for the combined assembly, where homozygous regions predominantly contained 2-copy *k*-mer, and heterozygous regions mostly contain 1-copy *k*-mer. This distribution aligns with expectations based on the presence of both complete parental haplotypes and minimal artefactual duplication (Supplementary Fig. 1). Additionally, conserved synteny blocks between *F. vesca* and FL 16.33-8 genomes were confirmed (Fig. 2D). When comparing three putative ancestral diploid strawberry genomes, *F*. *vesca*, *F*. *iinumae*, and *F. viridis*, the haplotype-phased genome of FL 16.33-8 showed a high level of collinearity with all diploid strawberry genomes (Supplementary Figs. S2 and S3).

The total TE content of haplotype-phased genomes of FL 16.33-8 were 345 Mb in 16.33-8-Phase1 and 343 Mb in 16.33-8-Phase2, including long terminal repeat (LTR) retrotransposons (Class I: 55.95% and 58.53%) and terminal inverted repeat (TIR) elements (Class II: 31.59% and 27.02%), respectively. TEs were evenly distributed across the 28 chromosomes of FL 16.33-8 with a small increase in content near the centromeres (Fig. 2A and Supplementary Table S7). The most abundant Class I TEs were LTR retrotransposons, particularly the *Gypsy* superfamily followed by *Copia*. Mutator was the most abundant among Class II TEs. Comparing TEs in both 16.33-8-Phase1 and 16.33-8-Phase2 revealed differences between each super-families, indicating a high level of divergence in TE families between the haplotype genomes. This divergence resulted in structural and sequential variations in protein coding genes and their regulatory sequences. The genome quality was further assessed by examining assembly continuity in repeat space using the LTR Assembly Index (LAI) (Ou *et al*., 2018). The adjusted LAI score of each 16.33-8-Phase1 and 16.33-8-Phase2 are 15 and 14, which fall within the “reference” standard range according to the LAI classification. Estimation of regional LAI in 1 Mb sliding windows demonstrated uniform and high-quality assembly continuity across the entire genome (Fig. 2A).

### Comparative Transcriptome and Genome Analysis Revealed Candidate Genes for the *RPc2*-Associated Resistance Against *P. cactorum*

Genome-wide transcriptome profiling analysis was conducted on two highly resistant (FL 12.75-77 and FL 11.28-34) and two highly susceptible accessions (FL 12.82-44 and FL 13.22-336) 96 hours after *P. cactorum* infection (Fig. 3A and Supplementary Table S8). The whole genome heatmap of differentially expressed gene (DEGs) revealed significant patterns of gene expression between the resistant (*H3*) and susceptible (*h3*) accessions post-pathogen infection (PI) (Fig. 3A, Supplementary Table S9). Infected roots were observed in both resistant and susceptible accessions after 96 hours of inoculation, but interestingly, newly developed roots were more frequently observed in resistant accessions compared to susceptible ones. Additionally, more severe disease symptoms were observed in the roots of susceptible accessions (Fig. 3A). A total of 409 DEGs (gray) were upregulated in resistant accessions after the pathogen infection (Fig. 3B and Supplementary Table S10). Notably, among the 34 DEGs identified from the comparison of resistant and susceptible accessions (*H3*-P.c Inoculation vs. *h3*-P.c Inoculation; gray ∩ orange), three DEGs (Fxa7Bg964820, Fxa7Bg964830, and Fxa2Bg234610) and four DEGs (Fxa3Ag419260, Fxa2Cg141260, Fxa2Dg203300, Fxa2Bg262750) were found to be closely associated with plant defense signaling and lignin biosynthesis pathways (Supplementary Fig. S4 and S5). In the volcano plot analysis of whole-genome DEGs, we identified distinct gene expression patterns. The two *cyclic nucleotide-gated ion channel 1-like* (*CNGCs*) genes: Fxa7Bg964820 and Fxa7Bg964830 were the most significantly upregulated in response to the pathogen inoculation. (Fig. 3C). There were number of upregulated DEGs with up to 10-fold changes including *WAK1*. In this study, we focused on *CNGC1*, *CNGC2*, and *WAK1*, as these DEGs are located in the *RPc2* region. Global analyses of DEGs in each pairwise comparison demonstrated significant alterations in gene expressions associated with ‘Binding’ and ‘Catalytic Activity’ across all comparisons following the inoculation of *P. cactorum* (Supplementary Fig. S6).

**Fig. 3.**
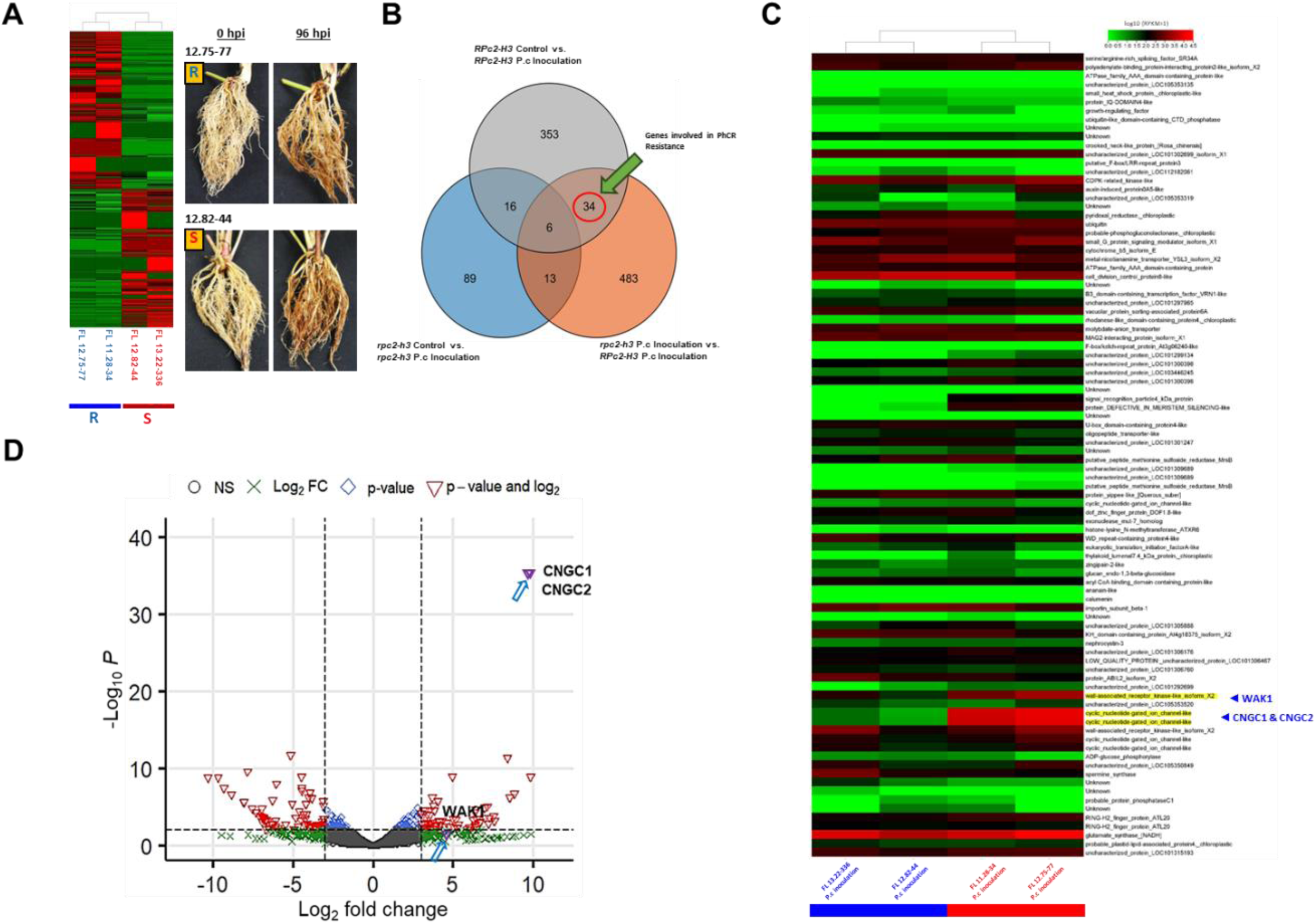
Differentially expressed genes associated with the resistance against *P*. *cactorum*. (A) Heatmap showing differentially expressed genes at the whole-genome level in response to inoculation with an oospore suspension. Disease symptoms were observed in the roots of resistant and susceptible genotypes 96 hours post-inoculation. (B) Venn diagram of upregulated DEGs in the water and oospore treatments between susceptible and resistance accessions. (C) Heatmap visualization of the genes present in the fine-mapped region (546 kb) of *RPc2.* Red box: susceptible genotypes, Blue box: resistant genotypes. The genes highly upregulated in the resistance genotype compared to the susceptible genotype, were highlighted in yellow. (D) Volcano plot comparing gene expression levels between susceptible and resistance accessions after infection. *CNGC1*, *CNGC2*, and *WAK1* are marked with purple triangle.

The transcriptome expression profiling was conducted on 92 genes located within the genomic region of the resistant haplotype, *H3*, in response to *P. cactorum* infection. Two *cyclic nucleotide-gated ion channel 1-like* (*CNGC1* and *CNGC2*; Fxa7Bg964820, Fxa7Bg964830) genes and *One Wall-Associated Kinase 1-like* (*WAK1*; Fxa7Bg964800) were constitutively upregulated in the resistant accessions compared to susceptible accessions (Fig. 3C, Supplementary Table S11). To identify genomic structure and causal variants associated with *RPc2*, the resistant haplotype genome, 16.33-8-Phase1 containing *H3*, was comparatively analyzed with ‘Florida Brilliance’ (FaFB1) and ‘Royal Royce’ (FaRR1). The genomic region of the resistant haplotype *H3* from FL 16.33-8 closely resembled that of ‘Royal Royce’, suggesting that ‘Royal Royce’ possesses the resistant haplotype *H3*. On the other hand, there were significant structural and sequence variations between the genomes of resistant (16.33-8-Phase1, FaRR1-Phase1 and FaRR1-Phase2) and susceptible (FaFB1 both haplotypes) accessions (Fig. 4A). The sequence variations of candidate genes *WAK1*, *CNGC1*, and *CNGC2* were specifically examined. The amino acid sequence of *WAK1* from 16.33-8-Phase1 was identical with both FaRR1 haplotypes and was 91.25% consistent with the *WAK1* in FaFB1 (Supplementary Fig. S7). Two copies of the *CNGC* gene (*CNGC1* and *CNGC2*) were exclusively identified in the genomes of 16.33-8-Phase1 and FaRR1, both carrying the resistance haplotype *H3*. The amino acid sequence similarity between CNGC1 and CNGC2 was 98.67% (Supplementary Fig. S8). Two homoeologous copies of WAK1 were found in chromosomes 7C and 7D, sharing 86% similarity. However, the amino acid sequences of WAK1 present in chromosome 7B were 30% consistent with a copy on chromosome 7C and 45% consistent with another copy on chromosome 7D (Supplementary Fig. S9). Two homoeologous copies of *CNGC1* and *CNGC2* were present in chromosomes 7A and 7D. The amino acid sequences of CNGC1 were 84.58% consistent with homoeologous copy on chromosome 7A, and 62.93% consistent with another homoeologous copy on chromosome 7D (Supplementary Fig. S10).

**Fig. 4.**
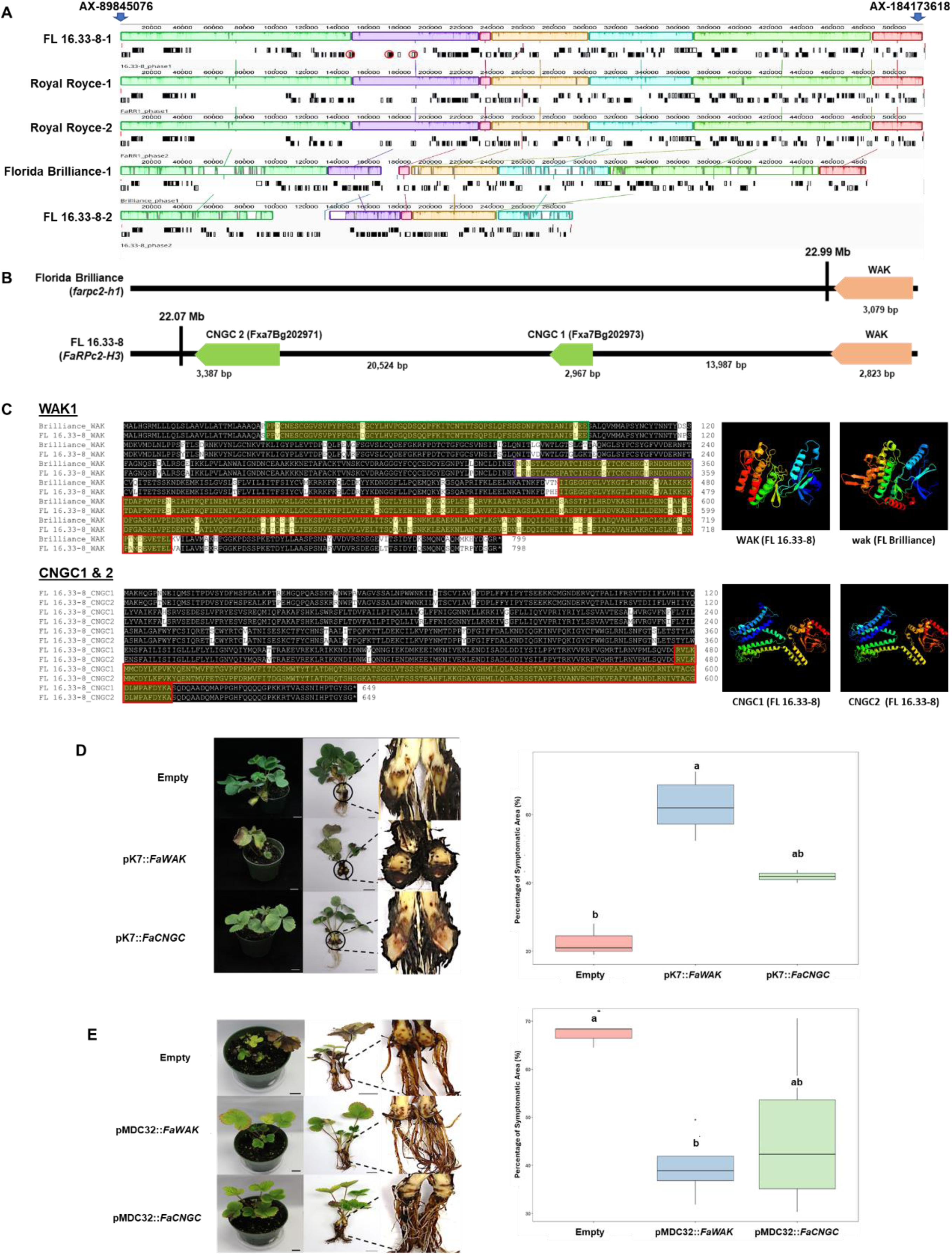
Identification of candidate genes for *RPc2* and functional validations in strawberry crowns and roots. **(A)** FL 16.33-8 phase1, ‘Royal Royce’ FaRR1 phase1 and phase2 represent the structure of resistance haplotype *RPc2*-H3, and ‘Florida Brilliance’ FaFB1 represents the structure of susceptible haplotype *RPc2*-H1 (referred to as *rpc-h3*). Mauve alignment of genomic region of *RPc2* between ‘Florida Brilliance’ (FaFB1), FL 16.33-8 phase1 and 2, ‘Royal Royce’ (FaRR1) phase1 and phase2. Each color indicates the locally collinear blocks. Same color of blocks represents local collinearity. Molecular marker linked to the candidate genes is indicated with a blue arrow. (B) Physical locations of WAK1, CNGC1, and CNGC2 in genomes of ‘Florida Brilliance’ and FL 16.33-8. (C) Sequence comparisons were conducted for WAK1 between resistant and susceptible varieties. Additionally, CNGC1 and CNGC2 sequences present in FL 16.33-8 Phase 1 genome were compared each other. Different functional domains and motifs were marked and described separately using different colors. 3D protein structures of WAK1, CNGC1, CNGC2 from FL 16.33-8, and WAK1 from ‘Florida Brilliance’. “(D) *Agrobacterium*-mediated transient gene expression of *WAK1* and *CNGC1/CNGC2* for RNAi-mediated knockdown in the moderately resistant ‘Florida Beauty’ (RPc2-H3). (E) overexpression of *WAK1* and *CNGC1* in susceptible ‘Florida Brilliance’ (*rpc2*-*h3*).

To examine the functions of *WAK1*,*CNGC1*, and *CNGC2*, we conducted *Agrobacterium*-mediated transient gene expression assays for RNAi-mediated knockdown and overexpression (Figs. 4D, and E). Due to the high sequence identity between *CNGC1* and *CNGC2*, it was not possible to target only one gene individually. Therefore, we designed the RNAi construct to knockdown the expression of both genes. For the overexpression study, however, we specifically cloned *CNGC1* into the overexpression vector to assess its effects for PhCR phenotype in a susceptible variety. The qRT-PCR results showed a significant downregulation of *WAK1* and *CNGC1/CNGC2* gene expression levels in RNAi plants and an upregulation in overexpression plants compared to the empty vector controls (Supplementary Fig. S11). The differences in disease symptoms between the RNAi and overexpression treatment groups were assessed using one-way ANOVA. The RNAi-mediated knockdown of *WAK1* and *CNGC1/CNGC2* in the moderately resistant ‘Florida Beauty’ (*H3*) displayed significantly increased symptomatic area compared to the control plants (empty vector) against *P. cactorum* (Figs. 4D; Supplementary Fig. 11B). The overexpression of *WAK1* and *CNGC1* in susceptible ‘Florida Brilliance’ (*h3*) resulted in 27.3% and 20.7% reductions in symptomatic area of crown respectively compared to control plants with empty vector treatments following *P. cactorum* infection (Figs. 4E; Supplementary Fig. 11A; Supplementary Table S12). However, the overexpression of *CNGC1* appears to be statistically less significant compared to the control plants (empty vector), likely due to variations between replicated experiments. The *WAK1*-RNAi plants shown in Fig. 4D are from the moderately resistant ‘Florida Beauty’ PhCR, while Fig. 4E features the susceptible ‘Florida Brilliance’ (*h3*) group for *WAK1* and *CNGC1*. These plants represent different genotypes, leading to distinct phenotypes. In Fig. 4D, *WAK1*-RNAi knockdown plants display intermediate browning in the crown and a lack of new bare roots, indicating a partial compromise in resistance. Conversely, Fig. 4E shows ‘Florida Brilliance’ with browning starting at the crown and also a failure to produce new bare roots, highlighting its susceptibility. Taken together, the results from comparative genomic and transcriptomic analyses indicated *WAK1*, *CNGC1*, and *CNGC2* as candidate genes involved in *RPc2*-mediated resistance to PhCR in strawberry.

### Population structure and selection signatures of the *RPc2* locus among genetically diverse strawberry accessions

We assessed the population structure of *RPc2* across diverse breeding populations (UCD: n=501, UF: n=61, other USA: n=267, Canada: n=42, Europe: n=91, Japan: n=46, Taiwan: n=3, Chile: n=6, Ecuador: n=1). Comparative genetic structure and principal component analysis (PCA) were performed using 76 SNP markers spanning the genomic region of *RPc2*, approximately 1.2 Mb. The results of the PCA showed that the accessions were segregated into four distinct groups, based on the presence of either the resistant or susceptible haplotype of *RPc2* (Fig. 5A, Supplementary Dataset2, Supplementary Table S13). Additionally, the dosage of resistant haplotype (*H3*) or susceptible haplotype groups (*h3*) was clearly discriminated by the PCA analysis. The group of susceptible haplotypes (*h3*) clustered in the intermediate region between homozygous- and heterozygous-*RPc2-H3*. As shown in Figure 5b, the phylogenetic analysis of 1,029 accessions largely categorized them into the groups with and without the resistant haplotype *H3* (Supplementary Dataset 3; Fig. 5B).

**Fig. 5.**
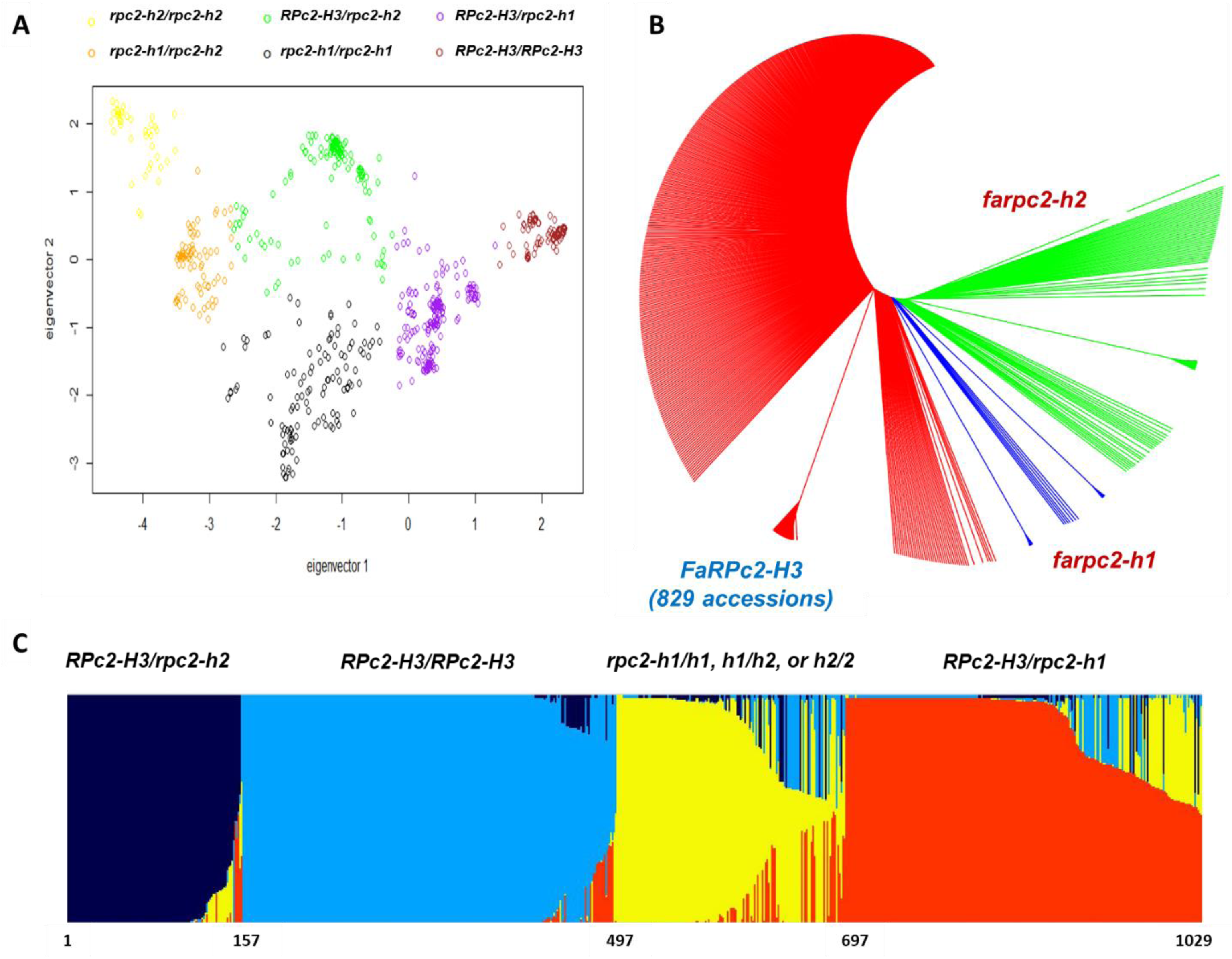
Genetic structure analysis of octoploid strawberry populations using 76 SNP markers in the 1.2 Mb of *RPc2* region. (A) Principal component analysis that shows different genetic clusters with and without the resistance haplotype, *RPc2-H3*. (B) Phylogenetic analysis revealed three groups characterized by *RPc2-h1*, *RPc2-h2*, and *RPc2-H3*. (C) Admixture analysis using Structure ver. 2.3.4 created four clusters with and without the resistance haplotype *RPc2-H3*.

Bayesian inference of population structure was also performed. The optimal number of clusters (K) was determined to be k=4, revealing 340 accessions as homozygous *RPc2-H3* (cluster 2) and 376 as heterozygous *RPc2-H3* (cluster 1 and cluster 4) (Fig. 5C, Supplementary Fig. S12, Supplementary Table S14). The cluster 1 (*RPc2-H3/rpc2-h2*) and cluster 4 (*RPc2-H3/rpc2-h1*) mainly comprise accessions with the heterozygous resistant *RPc2-H3* haplotype with the susceptible haplotypes. Investigation of the geographical origin revealed that relatively high frequency of resistant haplotype *RPc2-H3* was present in the US strawberry breeding accessions, respectively (Supplementary Table S15). This finding indicates that resistant *RPc2* alleles have undergone positive selection during strawberry breeding in the US. Additionally, the resistant haplotype *RPc2-H3* is well preserved in 829 (80.5%) accessions, with 60 out of 91 cultivars bred in Europe possessing *RPc2*-H3. This distribution supports the notion of a selective sweep driven by continuous selection of *P. cactorum*-resistant individuals in breeding programs worldwide.

## Discussion

In previous studies, a large-effect locus, *RPc2*, for resistance to crown rot caused by *P. cactorum* was initially identified in the UF breeding program through pedigree-based QTL detection (Mangandi et al., 2017). Subsequent multivariate genome-wide association studies (GWAS) confirmed the significance of the locus on chromosome 7B, describing a substantial portion of genetic variance and suggesting its necessity, though not sufficiency for resistance to PhCR (Jiménez *et al*., 2023). Despite its importance, this genomic region has not been fully characterized; thus this study highlights the need for further research to uncover potential resistance mechanisms and facilitate more effective breeding strategies to control PhCR.

This study developed a high-quality phased genome for the resistant accession FL16.33-8 and compared the *RPc2* locus across multiple genomes, revealing shared origins of the resistant haplotype between California and Florida breeding programs. Building upon previous studies (Lin *et al*., 2023; Lozada *et al*., 2021; Wang *et al*., 2021), the high-quality genome information would provide a foundation for identifying candidate genes associated with resistance traits and for elucidating the molecular mechanisms underlying the resistance to the soilborne pathogen. In this study, we identified two *Cyclic Nucleotide-Gated Channels* (*CNGC*) and one *Wall-Associated Kinase 1* (*WAK1*) gene for the resistance of PhCR.. Previous studies have highlighted the significance of *CNGC* genes in plant defense against pathogens (Yoshioka *et al*., 2006). It was also mentioned that calcium channel and other candidate genes with known plant defense functions could be associated with the *RPc2* resistance to PhCR (Jiménez *et al*., 2023). Similar findings have been reported in other plant species, where overexpression of *WAK* and *CNGC* genes conferred enhanced resistance to various pathogens (He *et al*., 1998; Navarro *et al*., 2004). These results underscore the importance of *WAK1* and *CNGC1* as key genetic factors in *RPc2*-mediated resistance in strawberries and provide valuable insights into the molecular mechanisms underlying resistance to *P. cactorum*.

The *WAK* play vital role in expansion of cell, protection against heavy metal and imparts resistance against plant-pathogens (He *et al*., 1998; Hou *et al*., 2005; Lally *et al*., 2001; Tang *et al*., 2017; Verica and He, 2002b). Protein kinase has also been reported to have function related to signaling during osmotic stress, keeping plant-specific cell wall and cytoskeleton continuum and also helps in closure of stomata by combining the environmental stimulus with cellular action with the help of H_2_O_2_ (Baluška *et al*., 2003; Desikan *et al*., 2004; Urao *et al*., 1999). These reports indicate that *WAK* has an important role against fungal pathogen inside the root cortex by initiating the defense response by modulating cytoskeleton-cell wall border. Also, *Phytophthora* species exudes hydrolase that damages the plant cell wall and leads to the degradation of homogalacturonan that ultimately releases Oligogalacturonides and these further binds with *WAK* to initiate defense response (Ferrari *et al*., 2013). The upregulation of *WAK* in response to *Phytophthora* has also been found in previous studies (Burra *et al*., 2014; Khatib *et al*., 2004; Toljamo *et al*., 2016; Xiao *et al*., 2019). *WAK* genes are receptor-like kinases localized in the cell wall, involved in recognizing pathogen-associated molecular patterns (PAMPs) and activating downstream defense responses (Supplementary Fig. S13) (Anderson *et al*., 2001; Verica and He, 2002a). Upon recognition of fungal PAMPs, *WAK* genes initiate signaling cascades that lead to the activation of various defense mechanisms, including the production of reactive oxygen species (ROS), the induction of pathogenesis-related (PR) genes, and the reinforcement of cell walls through callose deposition (Brutus *et al*., 2010; Kohorn and Kohorn, 2012).

During the sample collection of RNA-seq experiment, we often observed new root initiation near the crown in resistant plants after pathogen inoculation, while susceptible plants showed more severe root damage and lacked new root development near the crown as shown in Figure 3A. This suggests a potential role for *CNGC* and *WAK* in promoting root growth and development, particularly in response to soilborne pathogen infection, in resistant plant genotypes. MapMan analysis for the DEGs displayed upregulated genes involved in cell wall, phytohormone jasmonic acid, and wax in the resistant individuals in response to the pathogen (Fig. 6). Interestingly, the key genes in lignin biosynthesis are upregulated after the infection of *P. cactorum*, suggesting the important role of lignin in resistance of PhCR in strawberry. This observation has been also reported in Arabidopsis exposed to *Xanthomonas campestris* and flax (*Linum usitatissimum*) cell cultures treated with *Botrytis cinerea* where the expression of genes involved in lignin biosynthesis were observed (Hano *et al*., 2006; Lauvergeat *et al*., 2001; Zimmermann *et al*., 1999). The phytohormone jasmonic acid (JA), which plays an important role as a signaling molecule in plant defense mechanisms, was also upregulated in resistant plants (Campos *et al*., 2014).

**Fig. 6.**
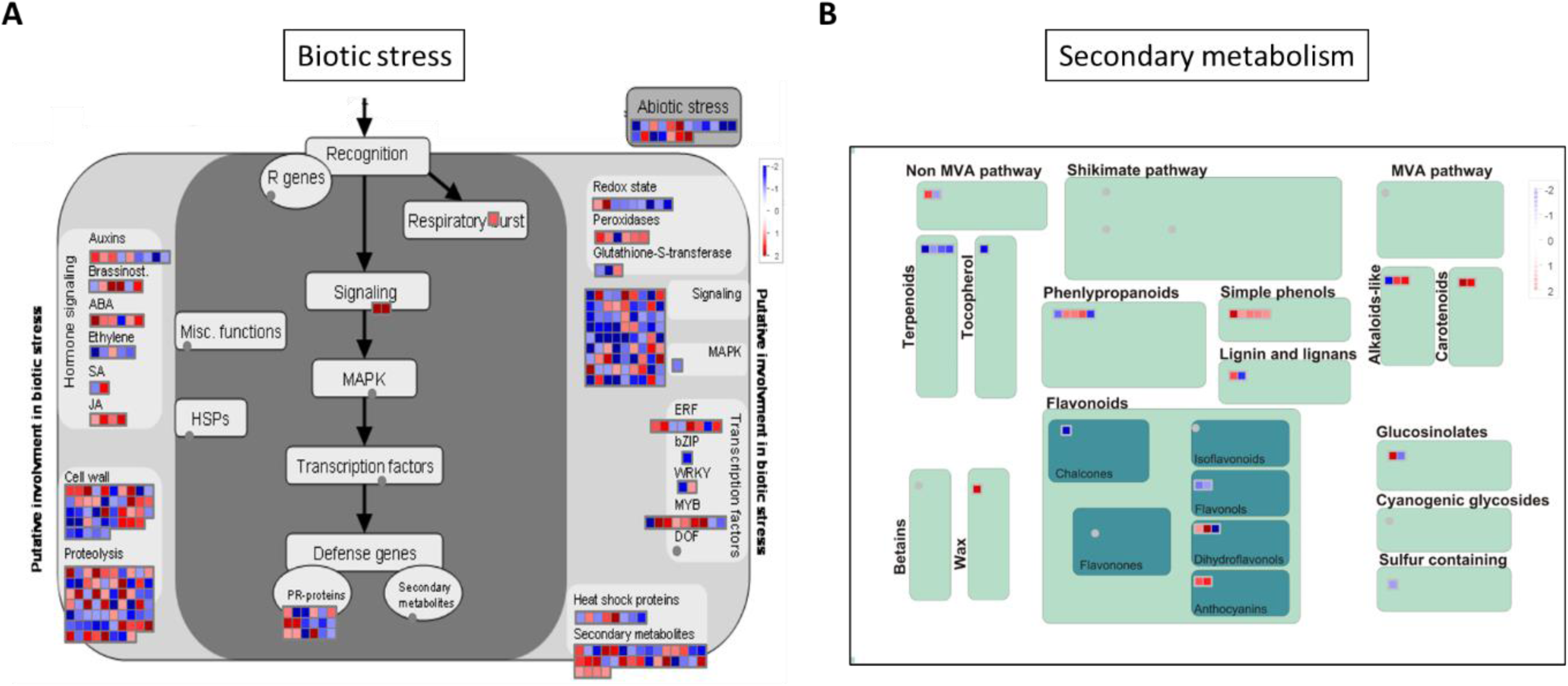
The MapMan-based classification of differentially expressed genes (DEGs) in response to *P. cactorum*. (A) Expression profiles of DEGs involved in biotic and abiotic stresses. (B) Expression profiles of DEGs involved in secondary metabolism. The scale bar represents positive (red) and negative (blue) regulation of gene expression based on log 2 fc scores.

*CNGCs* are known to have role in transport of ions, like Ca^2+^, through the plasma membrane and helps in regulating the defense signal (Deising, 2009). *CNGCs*, on the other hand, are plasma membrane-localized ion channels that regulate calcium influx in response to pathogen perception (Ma *et al*., 2009). Calcium signaling is a central component of plant defense signaling, triggering downstream immune responses such as the activation of mitogen-activated protein kinase (MAPK) cascades and the expression of defense-related genes (Ali *et al*., 2017; Ma *et al*., 2009). Additionally, *CNGCs* have been recognized for their involvement in mediating ion fluxes across the plasma membrane, which contribute to the generation of ROS and the establishment of an alkaline cytoplasmic pH, creating an unfavorable environment for fungal growth (Ali *et al*., 2017; Moeder *et al*., 2011). This gene has also been found to be an important defense-related gene in plants against *Phytophthora* infection (Ali *et al*., 2017; Toljamo *et al*., 2016; Wang *et al*., 2019). As shown in Figure 5, the plants with transient overexpression of *CNGC1* exhibited decreased levels of crown infection symptoms with newly developed roots but not in the empty vector control plants. CNGCs are not directly involved in lignin production in the plant cell wall, but they can play a role in signaling pathways that respond to biotic stresses, which may influence lignin biosynthesis (Hedrich and Neher et al., 2018). Genes directly involved in lignin biosynthesis and cell wall signaling pathways were expressed in resistant varieties against the PhCR pathogen (Figure 6). This finding demonstrates the involvement of the *WAK*-*CNGC* defense pathway would act as a frontline defense mechanism against soilborne pathogens, initiating rapid and effective immune responses to restrict the pathogen invasion and colonization in plant roots.

By examining the genetic structure of the *RPc2* locus in a large set of genetically diverse breeding populations, we were able to understand the population structures controlling the genetic variance for resistance to PhCR. Jiménez *et al*. (2023) evaluated genetically diverse strawberry populations for resistance to California pathogen isolates, identifying five highly resistant individuals (1.3%) and 27 moderately resistant individuals (5.5%). Our analysis of 30 individuals revealed that 27 belonged to the resistant *H3* haplotype. Importantly, many historically resistant cultivars, including ‘Cyclone’, ‘Massey’, ‘Addie’, ‘MD683’, and ‘Camarosa’ were developed over 30 years ago, suggesting a longstanding presence of the PhCR resistance. Positive selection for disease resistance genes during the breeding process has been well-documented in various crop species (Gross and Olsen, 2010; Mondragón-Palomino *et al*., 2002; Turner-Hissong *et al*., 2020). The selective sweep observed in our study, where the *H3* haplotype was found to be well-preserved in breeding populations globally, further supports the notion of positive selection for disease resistance alleles during breeding. This is consistent with the previous finding of the extremely high frequency of the *RPc2*-associated favorable allele in the UCD training population. Our genetic structure analysis of the *RPc2* locus across diverse breeding populations also revealed a high frequency of the resistant haplotype *H3*, particularly in strawberries bred in the US and Europe. This skewed distribution of *H3* suggests ongoing selection for resistance to PhCR in breeding programs worldwide. The identification and preservation of resistance alleles, whether known or unknown, through selective breeding strategies underscore the importance of continued efforts to enhance disease resistance in strawberry. These efforts could be important for sustainable agriculture, particularly in the face of evolving pathogen populations and changing environmental conditions.

In conclusion, our study has effectively elucidated the genetic architecture of *RPc2* underlying resistance against *P. cactorum*. The development of a high-quality haplotype-phased genome for the elite resistant accession containing the resistant haplotype *H3* facilitated the discovery of candidate genes, *WAK1*, *CNGC1*, and *CNGC2*, for resistance to PhCR. These findings facilitate understanding of the molecular mechanisms underlying the resistance against the soilborne pathogen, *P. cactorum*, in strawberries and provide valuable insights for developing strategies to enhance resistance in breeding programs. Furthermore, the integration of pan-genomic analyses confirmed the historical sharing of the *RPc2* genomic region during strawberry breeding in the US. Our study supports the evidence of positive selection for a favorable resistance allele, demonstrated by the widespread distribution of the *H3* haplotype across breeding populations in the US and potentially worldwide. These efforts highlight the importance of continued research and genomic-driven breeding efforts to effectively manage the disease and ensure the long-term sustainability of strawberry.

## Acknowledgements

This research is supported by grants from the United States Department of Agriculture National Institute of Food and Agriculture (NIFA) Specialty Crops Research Initiative (SCRI) “Delivering Breeding and Management Solutions to Prevent Losses to Emerging and Expanding Disease Threats in Strawberry” under award number (#2022-51181-38328). We also acknowledge Florida Strawberry Growers Association and the strawberry breeding lab members at UF/IFAS Gulf Coast Research and Education Center for their support of this study.

## Author contributions

H.H., S.C., Y.J., S.L conceived the hypothesis and original idea and designed the experiments. H.H., S.C., and U.G. performed the genome assembly and transcriptome analysis. Y.J., Y-H.N, and S.I. designed molecular markers to conduct the fine mapping analysis. S.V. constructed genetic linkage map to validate the collinearity. D.P., G.C., R.F., M.F., and S.K. provided field phenotype data of Phytophthora crow rot and SNP genotyping data. S.L., S.K., V.M. reviewed and edited the manuscript. All authors reviewed the manuscript.

## Data availability

RNA sequencing data for strawberry is available in NCBI SRA database with accession numbers of BioProject number PRJNA890188. DNA sequencing data for strawberry is available in NCBI SRA. The octoploid genome assembly and annotation files are available at Genome Database for Rosaceae (GDR; https://www.rosaceae.org/)

## Supplementary Data

Supplementary Fig. S1. Merqury’s k-mer-based assembly validation (A) The *k*-mer blob Plot illustrates the distribution of maternal (red) and paternal (blue) k-mer within each scaffold of the respective assemblies. The size of the blob corresponds to the total number of *k*-mers in that scaffold. (B) The combined spectra plot of inherited *k*-mers displays three peaks. The first peak (grey) signifies *k*-mers present in the raw reads but absent from the assembly due to sequencing errors. The second peak corresponds to *k*-mers from heterozygous regions, while the third peak corresponds to *k*-mers from homozygous regions. These plots indicate a complete and haplotype-resolved assembly.

## Supplementary Figure Legend

**Supplementary Fig. S1.** Merqury’s k-mer-based assembly validation. The combined spectra plot of inherited *k*-mers displays three peaks. The first peak (grey) signifies *k*-mers present in the raw reads but absent from the assembly due to sequencing errors. The second peak corresponds to *k*-mers from heterozygous regions, while the third peak corresponds to *k*-mers from homozygous regions. These plots indicate a complete and haplotype-resolved assembly.

**Supplementary Fig. S2.** Dotplot visualization of whole-genome alignment of the FL 16.33-8-phase1 assembly to the diploid *F. vesca* genome assembly ver. 4.0 using MCScanX.

**Supplementary Fig. S3.** Dotplot of FL 16.33-8-phase1 assembly to diploid *Fragaria vesca* ver. 4.0. (A), *F*. *iinumae* (B), and *F*. *viridis* (C). Dot plots are produced using the DGENIE software and alignments with minimap2.

**Supplementary Fig. S4.** KEGG pathway maps for plant-pathogen interaction in three pairwise comparison.

**Supplementary Fig. S5.** MapMan visualization of DEGs related to lignin biosynthesis pathways. The log2 fold changes of significant DEGs were imported and visualized in MapMan software (3.5.1 R2).

**Supplementary Fig. S6.** Gene ontology (GO) of enriched genes across all comparisons following the inoculation of *P*. *cactorum*.

**Supplementary Fig. S7.** Genomic, CDS, and peptide sequence alignment of *WAK1* gene derived from ‘Florida Brilliance’, FL 16.33-8-phase1, and ‘Royal Royce’ phase1 and 2 assemblies.

**Supplementary Fig. S8.** Genomic, CDS, and peptide sequence alignment of *CNGC* genes derived from FL 16.33-8-phase1, and ‘Royal Royce’ phase1 and 2 assemblies.

**Supplementary Fig. S9.** Genomic, CDS, and peptide sequence alignment of *WAK1* and homoeologous genes from FL 16.33-8-phase1 assembly.

**Supplementary Fig. S10.** Genomic, CDS, and peptide sequence alignment of *CNGC* and homoeologous genes from FL 16.33-8-phase1 assembly.

**Supplementary Fig. S11.** Gene expression analysis of *WAK1* and *CNGC1* by qRT-PCR in strawberry resistance to *P. cactorum* pathogens. (A) The transient expression assay was conducted employing the pMDC32 overexpression vector to drive the expression of *WAK1* and *CNGC1* genes. (B) The transient expression assay was conducted employing the pK7GWIWG(II) RNAi vector to drive the knockdown expression of *WAK1* and *CNGC1* genes. The x-axis represents the vector composition used. EV denotes ‘Empty Vector, ‘Error bars represent standard deviation. Asterisks indicate significant outcomes with P values of Student’s t-test (*P < 0.05, **P < 0.01, ***P < 0.001, Student’s t-test), respectively.

**Supplementary Fig. S12.** Delta K values for STRUCTURE analysis of octoploid strawberries.

**Supplementary Fig. S13.** A hypothetical model for the enhanced resistance against *Phytophthora cactorum* in the *WAK1*, *CNGC1* knockdown strawberry root. Differentially expressed genes identified in *WAK1* and *CNGC1* knockdown crown in response to *P*. *cactorum* were represented. DAMPs such as Oligogalacturonides, and pectin fractions are sensed by *WAK1*, leading to the activation of *FaRLCK*. Subsequently, the activated *FaRLCK* phosphorylates *CNGC1* induces extracellular Ca^2+^ influx and then triggers PTI-related gene expressions, which result in disease resistance in strawberry.

**Supplementary Dataset1.** Detailed information on Genetic linkage map for ‘Florida Brilliance’ and FL 16.33-8

**Supplementary Dataset2.** Dataset used for phylogenetic analysis and principal component analysis.

**Supplementary Dataset3.** Dataset used for structure analysis.

