## Supplementary Figures for "Genomic Architecture of the *Resistance to Phytophthora Cactorum 2* (*RPc2*) Locus in Strawberry (*Fragaria* × *ananassa*)"

Unraveling Genomic Structures and Candidate Genes for Resistance to Phytophthora Crown Rot in Strawberry (*F. ×ananassa*)

### Contents

**Supplementary Fig. S1. Merqury's k-mer-based assembly validation.**

**Supplementary Fig. S2. Dotplot visualization of whole-genome alignment of the FL 16.33-8-phase1 assembly to the diploid *F. vesca* genome assembly ver. 4.0 using MCScanX.**

**Supplementary Fig. S3. Dotplot of FL 16.33-8-phase1 assembly to diploid *Fragaria vesca* ver. 4.0.**

**Supplementary Fig. S4. KEGG pathway maps for plant-pathogen interaction in three pairwise comparison.**

**Supplementary Fig. S5. MapMan visualization of DEGs related to lignin biosynthesis pathways.**

**Supplementary Fig. S6. Gene ontology (GO) of enriched genes across all comparisons following the inoculation of *P. cactorum.***

**Supplementary Fig. S7. Genomic, CDS, and peptide sequence alignment of *WAK* gene derived from 'Florida Brilliance', FL 16.33-8-phase1, and 'Royal Royce' phase1 and 2 assemblies.**

**Supplementary Fig. S8. Genomic, CDS, and peptide sequence alignment of *CNGC* genes derived from FL 16.33-8-phase1, and 'Royal Royce' phase1 and 2.**

**Supplementary Fig. S9. Genomic, CDS, and peptide sequence alignment of *WAK* and homoeologous genes from FL 16.33-8-phase1 assembly.**

**Supplementary Fig. S10. Genomic, CDS, and peptide sequence alignment of *CNGC* and homoeologous genes from FL 16.33-8-phase1 assembly.**

**Supplementary Fig. S11. Gene expression fold change of *FaWAK* (A) and *FaCNGC* (B) by qRT-PCR in strawberry resistance to *P. cactorum* pathogens.**

**Supplementary Fig. S12. Delta K values for STRUCTURE analysis of octoploid strawberries.**

**Supplementary Fig. S13. A hypothetical model for the enhanced resistance against *Phytophthora cactorum* in the *FaWAK*, *FaCNGC1* knockdown strawberry root.**

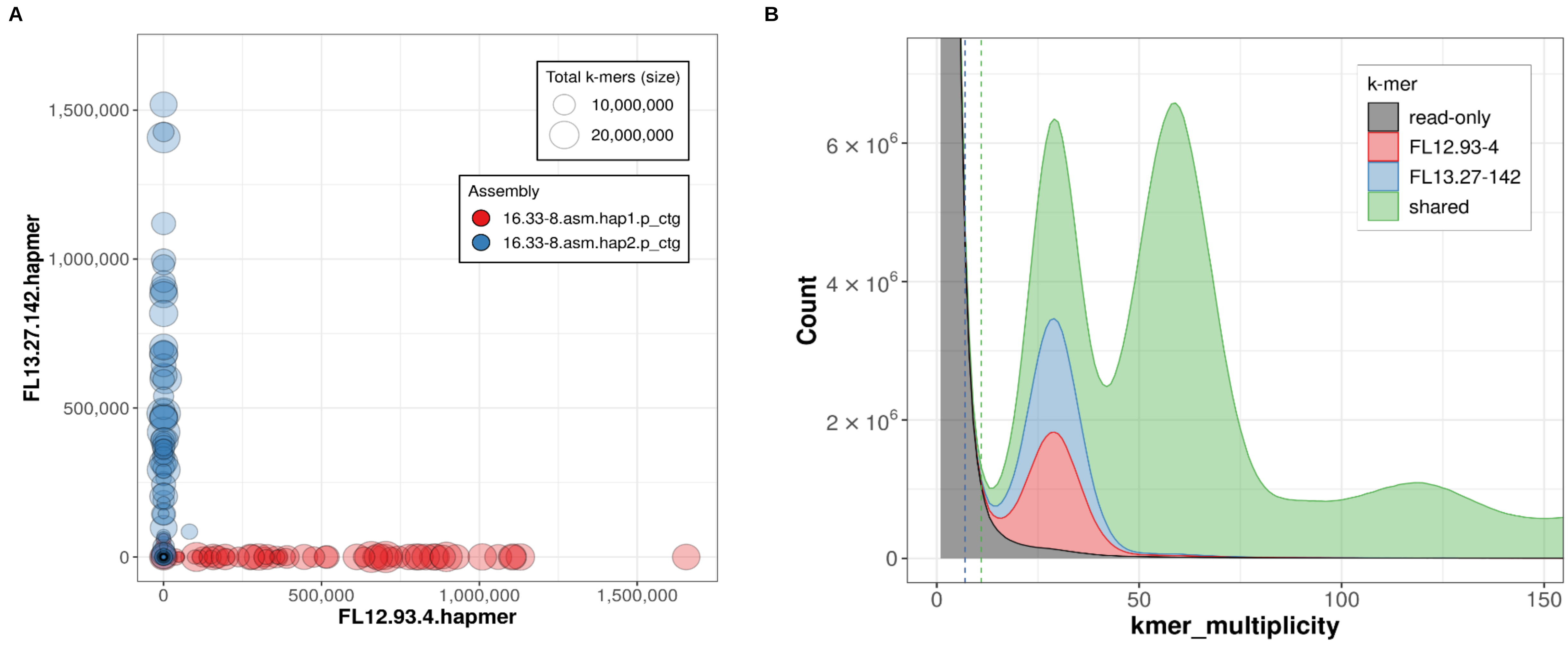


**Supplementary Fig. S1. Merqury's k-mer-based assembly validation. (A)** The *k*-mer blob Plot illustrates the distribution of maternal (red) and paternal (blue) k-mer within each scaffold of the respective assemblies. The size of the blob corresponds to the total number of *k*-mers in that scaffold. **(B)** The combined spectra plot of inherited *k*-mers displays three peaks. The first peak (grey) signifies *k*-mers present in the raw reads but absent from the assembly due to sequencing errors. The second peak corresponds to *k*-mers from heterozygous regions, while the third peak corresponds to *k*-mers from homozygous regions. These plots indicate a complete and haplotype-resolved assembly.

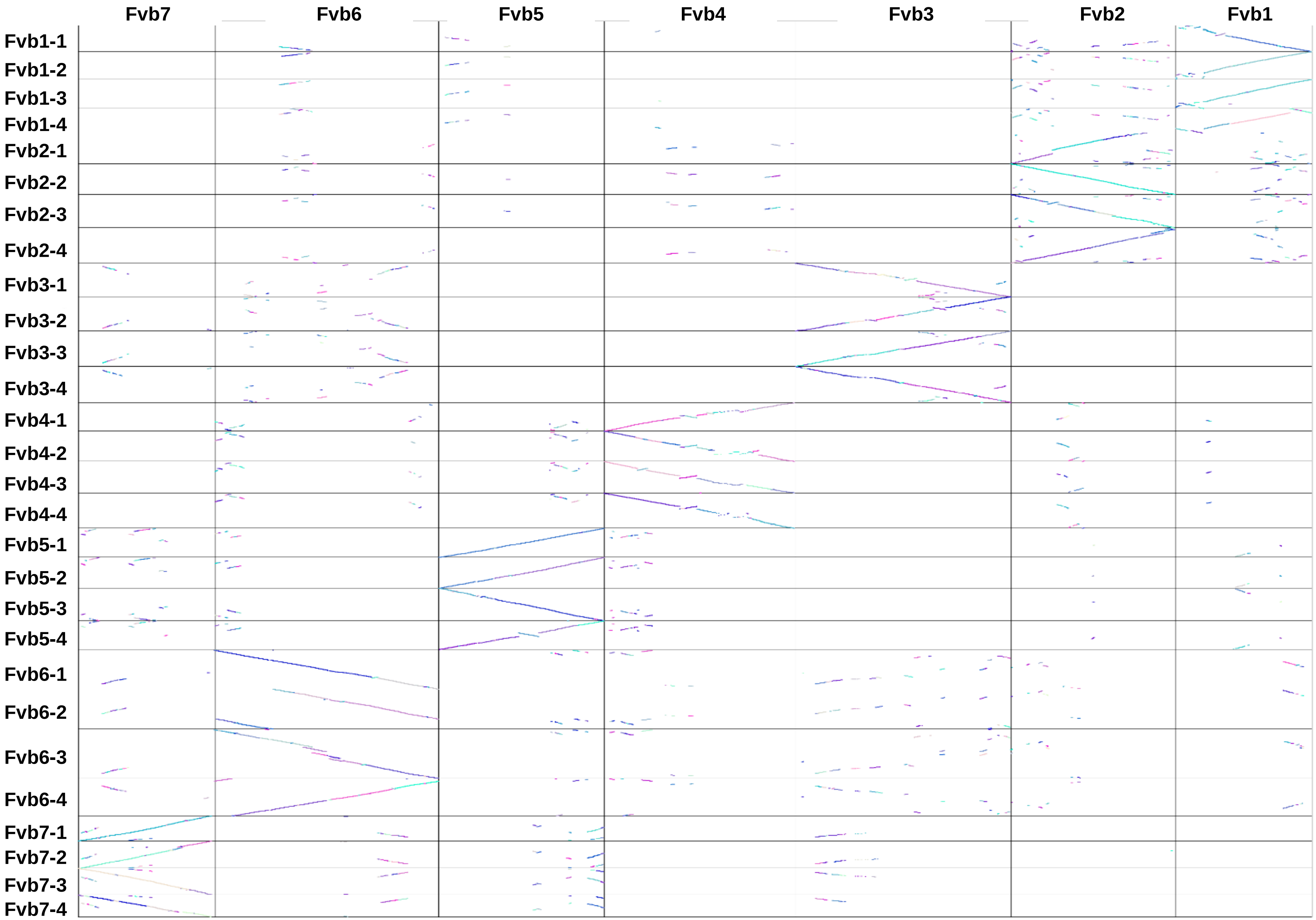


**Supplementary Fig. S2. Dotplot visualization of whole-genome alignment of the FL 16.33-8-phase1 assembly to the diploid *F. vesca* genome assembly ver. 4.0 using MCScanX.**

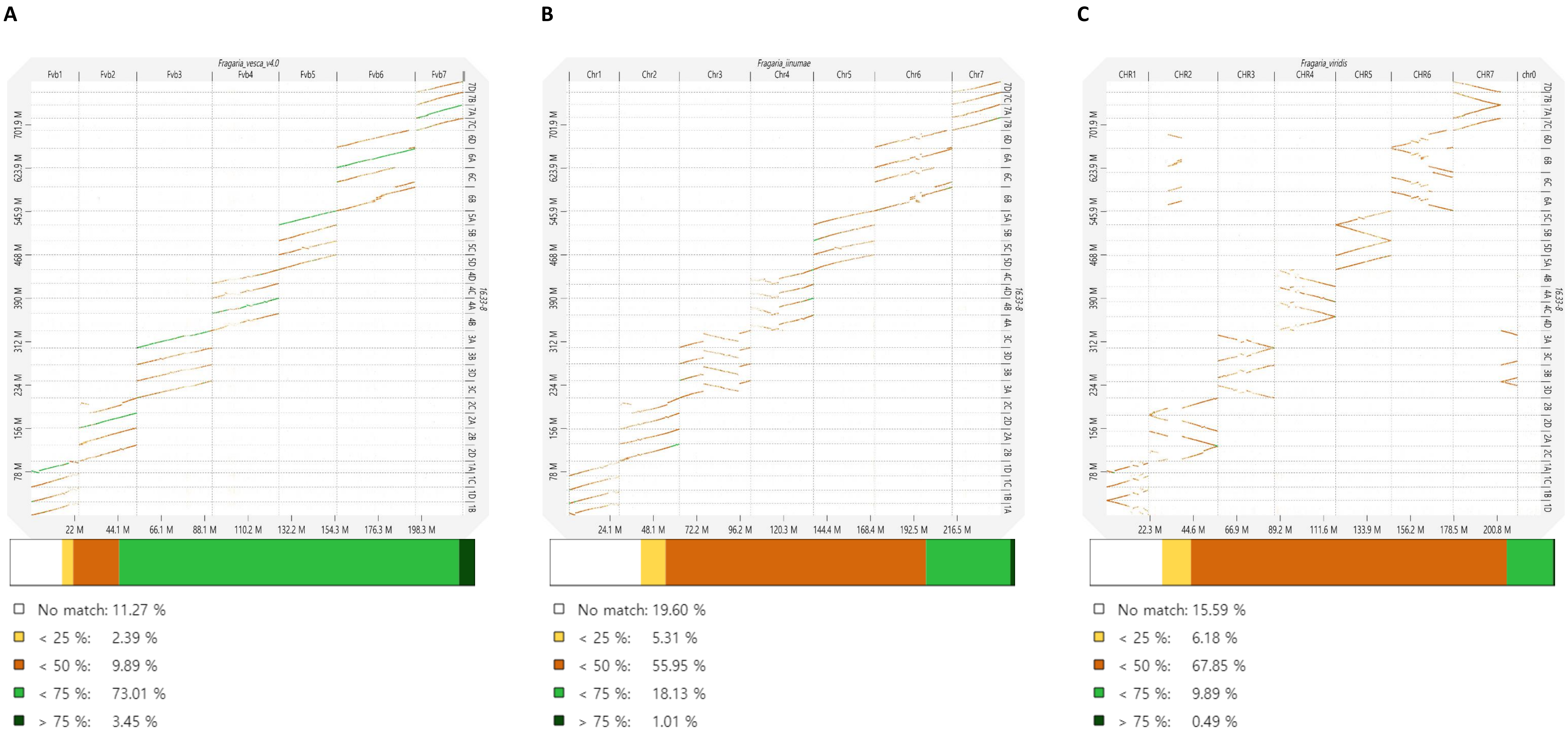


**Supplementary Fig. S3. Dotplot of FL 16.33-8-phase1 assembly to diploid *Fragaria vesca* ver. 4.0. (A)**, *F. iinumae* **(B)**, and *F. viridis* **(C)**. Dot plots are produced using the DGENIE software and alignments with minimap2.

A B

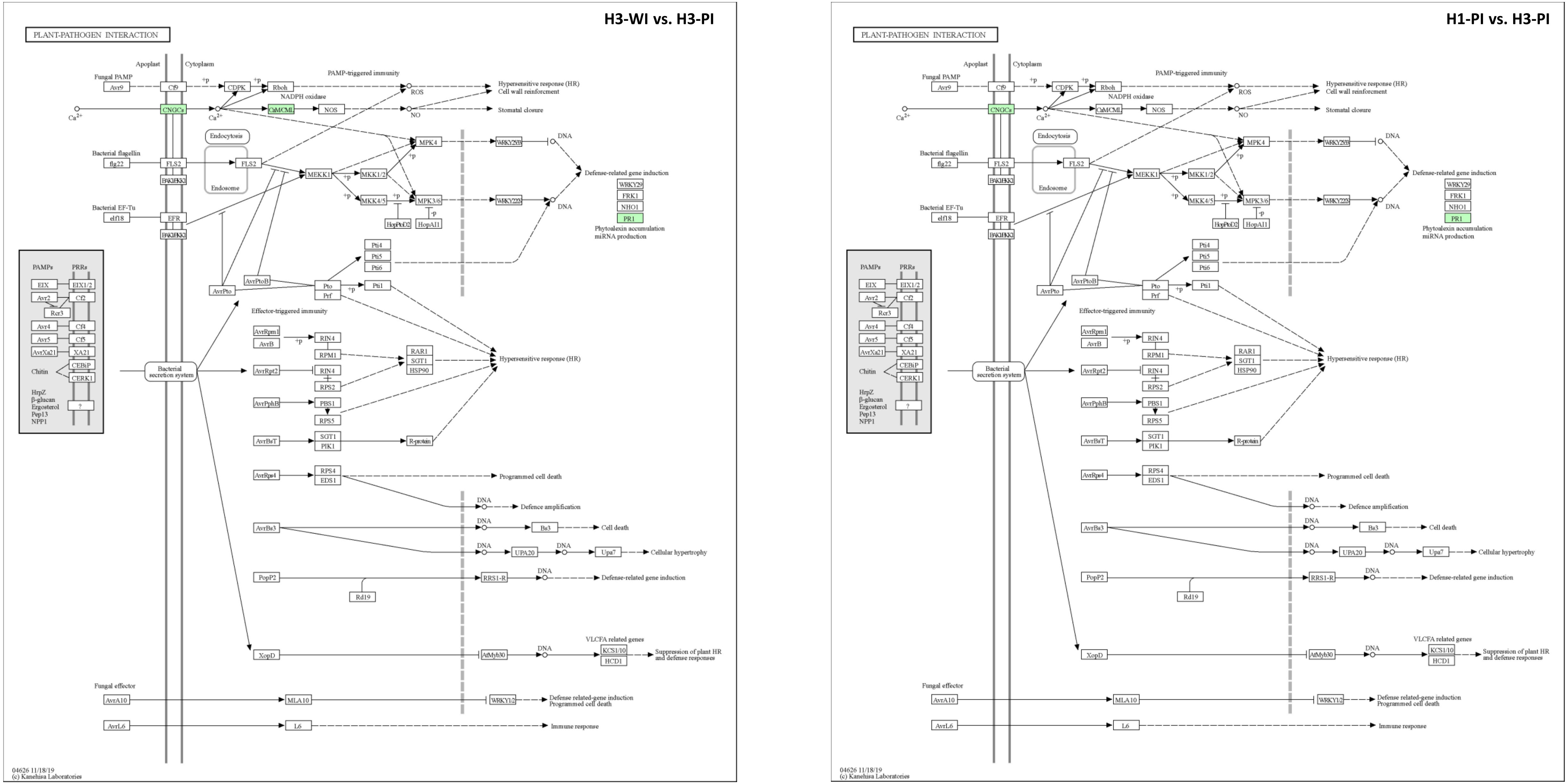


**Supplementary Fig. S4. KEGG pathway maps for plant-pathogen interaction in three pairwise comparison.** H3-WI vs. H3-PI (A) and H1-PI vs. H3-PI (B). Green indicates that map objects exist and are linked to corresponding entries. Abbreviations; cyclic nucleotide gated channel (CNCG), PR1 (pathogenesis-related protein 1), and CALM (calmodulin).

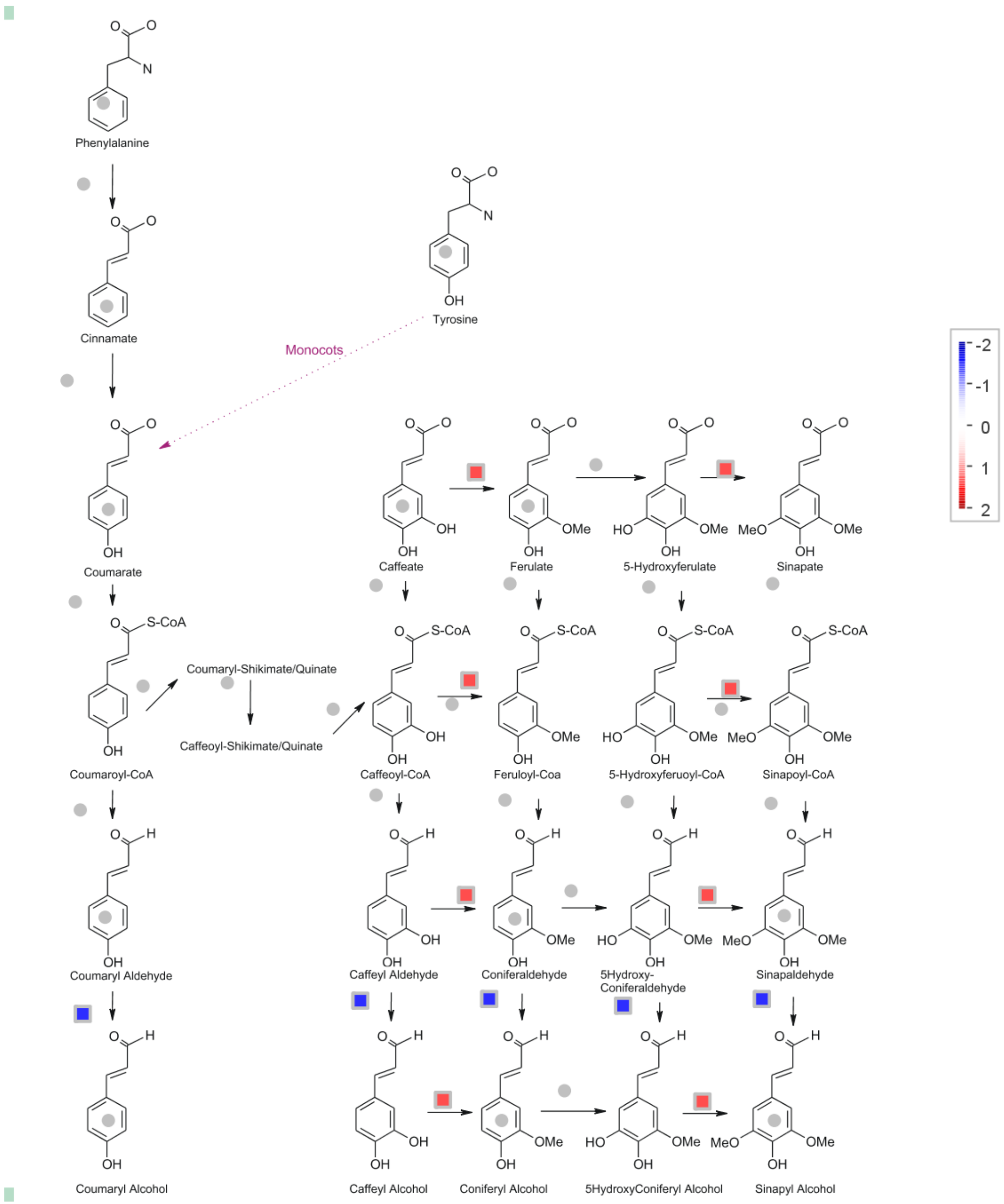


**Supplementary Fig. S5. MapMan visualization of DEGs related to lignin biosynthesis pathways. The log2 fold changes of significant DEGs were imported and visualized in MapMan software (3.5.1 R2).**

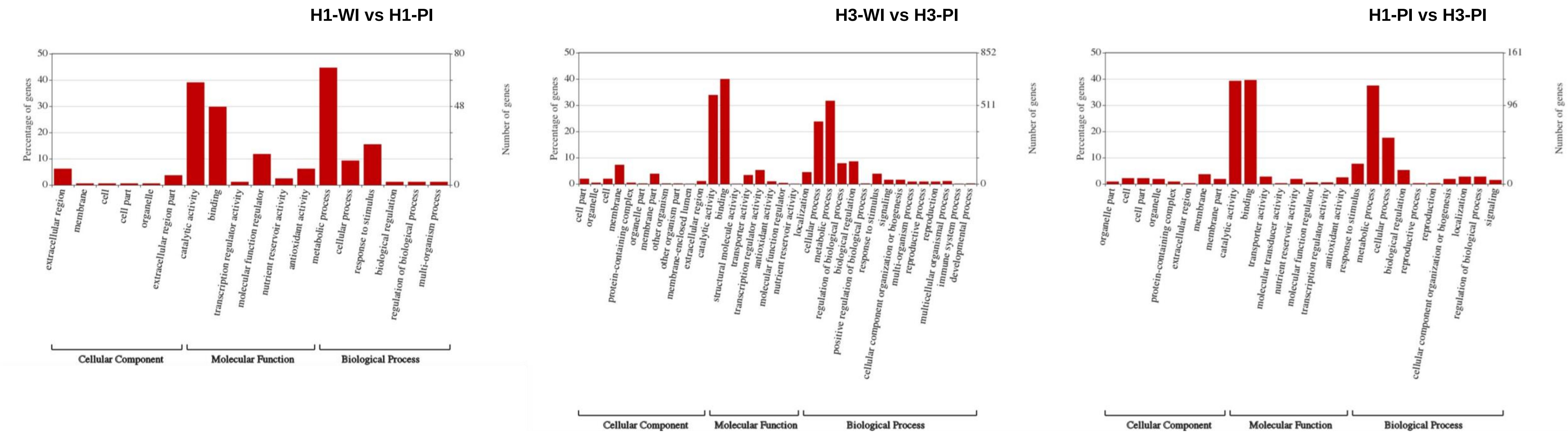


**Supplementary Fig. S6. Gene ontology (GO) of enriched genes across all comparisons following the inoculation of *P. cactorum***

**Supplementary Fig. S7. Genomic, CDS, and peptide sequence alignment of *WAK* gene derived from 'Florida Brilliance', FL 16.33-8-phase1, and 'Royal Royce' phase1 and 2 assemblies.**

**Genomic sequence of *WAK***

```
Brilliance        ATGGCCTTACATGGGAGAATGCTTCTTTTGCAACTCTCTTTGGCCGCAGTGCTATTAGCA    60
FL 16.33-8-Hap1   ATGGCCTTACATGGGAGAATGCTTCTTTTGCAACTCTCTTTGGCCGCAGTGCTATTAGCA    60
RR_7Ba            ATGGCCTTACATGGGAGAATGCTTCTTTTGCAACTCTCTTTGGCCGCAGTGCTATTAGCA    60
RR_7Bb            ATGGCCTTACATGGGAGAATGCTTCTTTTGCAACTCTCTTTGGCCGCAGTGCTATTAGCA    60

Brilliance        ACGACAATGCTAGCAGCTGCTCAAGCCCCGCCGCCTGACTGCAATGAGAGCTGCGGTGGT    120
FL 16.33-8-Hap1   ACGACAATGCTAGCAGCTGCTCAAGCCTCACCGCCTGTCTGCAATGAGAGCTGCGGTGGT    120
RR_7Ba            ACGACAATGCTAGCAGCTGCTCAAGCCTCACCGCCTGTCTGCAATGAGAGCTGCGGTGGT    120
RR_7Bb            ACGACAATGCTAGCAGCTGCTCAAGCCTCACCGCCTGTCTGCAATGAGAGCTGCGGTGGT    120

Brilliance        GTCTCAGTTCCATATCCATTTGGTCTCACTGAGGGTTGTTACCTACATGTGCCGGGACAA    180
FL 16.33-8-Hap1   GTCTCAGTTCCATATCCATTTGGTCTCACTGATGGTTGTTACCTACATGTGCCGGGACAA    180
RR_7Ba            GTCTCAGTTCCATATCCATTTGGTCTCACTGATGGTTGTTACCTACATGTGCCGGGACAA    180
RR_7Bb            GTCTCAGTTCCATATCCATTTGGTCTCACTGATGGTTGTTACCTACATGTGCCGGGACAA    180

Brilliance        GATTCTCAGCAGCCATTCAAGATCACTTGCAACACCACCACCTCGCAACCATCCCTACAG    240
FL 16.33-8-Hap1   GATTCTCAGCAGCCATTCAAGATCACTTGCAACACCACCACCTCGCAACCATCCCTACAG    240
RR_7Ba            GATTCTCAGCAGCCATTCAAGATCACTTGCAACACCACCACCTCGCAACCATCCCTACAG    240
RR_7Bb            GATTCTCAGCAGCCATTCAAGATCACTTGCAACACCACCACCTCGCAACCATCCCTACAG    240

Brilliance        TTCTCGGACAGCGATAACTTCCCCACAAATATTGCCAACATTTTTGTGGAAGAGAGTGCG    300
FL 16.33-8-Hap1   TTCTCGGACAGCGATAACTTCCCCACAAATATTGCCAACATTTTTTTGGAAGAGAGTGCG    300
RR_7Ba            TTCTCGGACAGCGATAACTTCCCCACAAATATTGCCAACATTTTTTTGGAAGAGAGTGCG    300
RR_7Bb            TTCTCGGACAGCGATAACTTCCCCACAAATATTGCCAACATTTTTTTGGAAGAGAGTGCG    300

Brilliance        CTGCAAGTAATGATGGCTCCGTCCTATAACTGCTACACCAACAATACATAGGACTCCTCC    360
FL 16.33-8-Hap1   CTGCAAGTAATGATGGCCCCGTCCTATAACTGCTACACCAACAATACATACAACCCCTCC    360
RR_7Ba            CTGCAAGTAATGATGGCCCCGTCCTATAACTGCTACACCAACAATACATACAACCCCTCC    360
RR_7Bb            CTGCAAGTAATGATGGCCCCGTCCTATAACTGCTACACCAACAATACATACAACCCCTCC    360

Brilliance        ATGGACAAAGTCATGGACCTCAATCTTCCGCCTTCTTTCACCCTCTCCGACAGAAACAAG    420
FL 16.33-8-Hap1   ATGGACAAAGTCATGGACCTCAATCTTCCGCCTTCTTACACCCTCTCCAACAGAAACAAG    420
RR_7Ba            ATGGACAAAGTCATGGACCTCAATCTTCCGCCTTCTTACACCCTCTCCAACAGAAACAAG    420
RR_7Bb            ATGGACAAAGTCATGGACCTCAATCTTCCGCCTTCTTACACCCTCTCCAACAGAAACAAG    420

Brilliance        GTTTATAACCTTGGGTGCAACAAAGTAACGAAGCTCATAGGTTATCCTCTTGAAGTTACA    480
FL 16.33-8-Hap1   GTTTATAACCTTGGGTGCAACAAAGTAACGAAACTCATAGGTTATCCTCTTGAAGTTACA    480
RR_7Ba            GTTTATAACCTTGGGTGCAACAAAGTAACGAAACTCATAGGTTATCCTCTTGAAGTTACA    480
RR_7Bb            GTTTATAACCTTGGGTGCAACAAAGTAACGAAACTCATAGGTTATCCTCTTGAAGTTACA    480

Brilliance        GACCCTGACCAGCTCTTTGAAGTAGGGTTCAGCGGTGTAAGCTTGTGCCAAGATGAATTT    540
FL 16.33-8-Hap1   GACCCTGAGCAGCTCTTTCAAGTACGGTTCAGCGGTGTAAGCTTGTGCCAAGATGAATTC    540
RR_7Ba            GACCCTGAGCAGCTCTTTCAAGTACGGTTCAGCGGTGTAAGCTTGTGCCAAGATGAATTC    540
RR_7Bb            GACCCTGAGCAGCTCTTTCAAGTACGGTTCAGCGGTGTAAGCTTGTGCCAAGATGAATTC    540

Brilliance        GGCAAAAGATTCCCCGACACTTGCACCGGATTTGGGTGCTCCGTAAATTCTATCCCTATC    600
FL 16.33-8-Hap1   GGCAAGAGATTCCCCGACACTTGCACCGGATTCGGGTGCTCCGTAAATTCTATCCCTAGC    600
RR_7Ba            GGCAAGAGATTCCCCGACACTTGCACCGGATTCGGGTGCTCCGTAAATTCTATCCCTAGC    600
RR_7Bb            GGCAAGAGATTCCCCGACACTTGCACCGGATTCGGGTGCTCCGTAAATTCTATCCCTAGC    600

Brilliance        GGACTGCAGAATATTACGTTGGGTGTGTGGACACTCGGTACCAGCCTCGGAACAACGGCG    660
FL 16.33-8-Hap1   GGACTGCAGAATATTACGGTGGATGTGTGGACACTCGGTACCGGCCTCGGAAAAACGGAG    660
RR_7Ba            GGACTGCAGAATATTACGGTGGATGTGTGGACACTCGGTACCGGCCTCGGAAAAACGGAG    660
RR_7Bb            GGACTGCAGAATATTACGGTGGATGTGTGGACACTCGGTACCGGCCTCGGAAAAACGGAG    660

Brilliance        CAGTGGGGCCTGAGCTACCCTTGCAGTTACGGTTTCGTTGTGGATGAACGCAACTTCACA    720
FL 16.33-8-Hap1   CAGTGGGGACTGAGCTACCCTTGCAGTTACGGCTTCGTTGTGGATGAACGCAACTTCACA    720
RR_7Ba            CAGTGGGGACTGAGCTACCCTTGCAGTTACGGCTTCGTTGTGGATGAACGCAACTTCACA    720
RR_7Bb            CAGTGGGGACTGAGCTACCCTTGCAGTTACGGCTTCGTTGTGGATGAACGCAACTTCACA    720

Brilliance        TTCGCCGGGAATCAAAGTTTTGATGCACTGAGGTCCGGCGAAATAAAGAAGCTTCCGGTG    780
FL 16.33-8-Hap1   TTCGCCGGGAATCAAAGTTTTGTTGCACTGAGGTCCGGCGCAATAAAGAAGCTTCCGGTG    780
RR_7Ba            TTCGCCGGGAATCAAAGTTTTGTTGCACTGAGGTCCGGCGCAATAAAGAAGCTTCCGGTG    780
RR_7Bb            TTCGCCGGGAATCAAAGTTTTGTTGCACTGAGGTCCGGCGCAATAAAGAAGCTTCCGGTG    780

Brilliance        CTTGCTAATTGGGCAATTGGGAATGACAACTGCGAAGCAGCAAAGAAGAAGAACGAGACT    840
FL 16.33-8-Hap1   CTTGCTAATTGGGCGATTGGGAATGACAACTGCGAAGCAGCAAAGAAGAAGAACGAGACT    840
RR_7Ba            CTTGCTAATTGGGCGATTGGGAATGACAACTGCGAAGCAGCAAAGAAGAAGAACGAGACT    840
RR_7Bb            CTTGCTAATTGGGCGATTGGGAATGACAACTGCGAAGCAGCAAAGAAGAAGAACGAGACT    840

Brilliance        GCTTTTGCATGCAAGACTGTGAATTCCAAGTGTGTCGACCGTGCTGGTGGTGGTTACTTT    900
FL 16.33-8-Hap1   GCTTTTGCATGCAAGACTGTGAATTCCAAGTGTGTCGACCGTGCTGGTGGTGGTTACTTT    900
RR_7Ba            GCTTTTGCATGCAAGACTGTGAATTCCAAGTGTGTCGACCGTGCTGGTGGTGGTTACTTT    900
RR_7Bb            GCTTTTGCATGCAAGACTGTGAATTCCAAGTGTGTCGACCGTGCTGGTGGTGGTTACTTT    900

Brilliance        TGCCAGTGTGAGGATGGTTACGAAGGGAACCCATACCTACTCGATAATTGCCTAGGTAA-    959
```

```
FL 16.33-8-Hap1 TGCCAGTGTGAGGATGGTTACGAAGGGAACCCATACCTCCCCGATAATTGCCTAGGTAAT   960
RR_7Ba          TGCCAGTGTGAGGATGGTTACGAAGGGAACCCATACCTCCCCGATAATTGCCTAGGTAAT   960
RR_7Bb          TGCCAGTGTGAGGATGGTTACGAAGGGAACCCATACCTCCCCGATAATTGCCTAGGTAAT   960

Brilliance      TACACACACAAATCATCTTAATTTCGTGTTAATTTTGTGATATATTTATATATATATACA  1019
FL 16.33-8-Hap1 TAATTACAAAAATCATCTTAATTTCGCGTTAA-TTTGTGATCTATATATAGGAAAGTG--  1017
RR_7Ba          TAATTACAAAAATCATCTTAATTTCGCGTTAA-TTTGTGATCTATATATAGGAAAGTG--  1017
RR_7Bb          TAATTACAAAAATCATCTTAATTTCGCGTTAA-TTTGTGATCTATATATAGGAAAGTG--  1017

Brilliance      CACACAGAGCTTCTTAGTTGCGAACGTCCGCATTTATTATTACGGTGCGGATTTCTATTT  1079
FL 16.33-8-Hap1 ------------------------------------------------------------  1017
RR_7Ba          ------------------------------------------------------------  1017
RR_7Bb          ------------------------------------------------------------  1017

Brilliance      TACACTCACTTTTCAATTGAATTTTCACATATCCACCGTCTAGTATTTAGATACTAATGT  1139
FL 16.33-8-Hap1 ------------------------------------------------------------  1017
RR_7Ba          ------------------------------------------------------------  1017
RR_7Bb          ------------------------------------------------------------  1017

Brilliance      ATAGATCATCTCTGCAAAATTTCACCCAATTTGGTTATCATTAAAGCACTCAAAACTGTA  1199
FL 16.33-8-Hap1 ------------------------------------------------------------  1017
RR_7Ba          ------------------------------------------------------------  1017
RR_7Bb          ------------------------------------------------------------  1017

Brilliance      TTTTTCTATTATAAACATGAACGGTTCAAGTTCGACAGAATTATGTTTATCTATTGTTTT  1259
FL 16.33-8-Hap1 ------------------------------------------------------------  1017
RR_7Ba          ------------------------------------------------------------  1017
RR_7Bb          ------------------------------------------------------------  1017

Brilliance      AATCGAGTTTGAGTGTCTTAATGATAACCAAATTTGCTAAAATTTTGCATAGATGATCTA  1319
FL 16.33-8-Hap1 ------------------------------------------------------------  1017
RR_7Ba          ------------------------------------------------------------  1017
RR_7Bb          ------------------------------------------------------------  1017

Brilliance      TACTTTAGTATCTAAAGAGTAGACAGTGGAGATGTGAAATTTCGATTGAAAAATGAGTGT  1379
FL 16.33-8-Hap1 ------------------------------------------------------------  1017
RR_7Ba          ------------------------------------------------------------  1017
RR_7Bb          ------------------------------------------------------------  1017

Brilliance      AAAGTGAAAATCCACACCGTAAGAATAAATACGGACGTCCGCAGCTGAGAAGCTCTGGAT  1439
FL 16.33-8-Hap1 ------------------------------------------------------------  1017
RR_7Ba          ------------------------------------------------------------  1017
RR_7Bb          ------------------------------------------------------------  1017

Brilliance      ATATATGTAGTTTTTAGATTTGATTTATATAGCTTCCCCTAAATGTAATATATGTCTTTT  1499
FL 16.33-8-Hap1 ------ATTGTTTCTATATTTGCTTTATATATAGCTTCC----CCTAATGTGATGTCTTT  1067
RR_7Ba          ------ATTGTTTCTATATTTGCTTTATATATAGCTTCC----CCTAATGTGATGTCTTT  1067
RR_7Bb          ------ATTGTTTCTATATTTGCTTTATATATAGCTTCC----CCTAATGTGATGTCTTT  1067

Brilliance      TTGTTTCAGATATCAATGAGTGCAATAAGAACTCAACCCTTTGCAGCGGTCCTGCAACTT  1559
FL 16.33-8-Hap1 TTGTTTCAGATATCAATGAGTGC---AAGCATTCAACCCTTTGCAGTGGTCCTGCAACTT  1124
RR_7Ba          TTGTTTCAGATATCAATGAGTGC---AAGCATTCAACCCTTTGCAGTGGTCCTGCAACTT  1124
RR_7Bb          TTGTTTCAGATATCAATGAGTGC---AAGCATTCAACCCTTTGCAGTGGTCCTGCAACTT  1124

Brilliance      GCATAAACTCAATTGGAGGTTACACCTGTAAATGTCACAAGGGCTATAGAAATGATGACC  1619
FL 16.33-8-Hap1 GCATAAACTCAATTGGAAGTTACACCTGTAAATGTCACAAGGGCCATAGAAATGATGACC  1184
RR_7Ba          GCATAAACTCAATTGGAAGTTACACCTGTAAATGTCACAAGGGCCATAGAAATGATGACC  1184
RR_7Bb          GCATAAACTCAATTGGAAGTTACACCTGTAAATGTCACAAGGGCCATAGAAATGATGACC  1184

Brilliance      ACGACAAGAATAAGTGTGTCCAAATCACTGAAACCTCCAGCAAAAATGACAAAGAAATGA  1679
FL 16.33-8-Hap1 ACGACAAGAATAAGTGTGTCCTAATCACTGAAACCTCCAGCAAAAATGACAAAGAAATGA  1244
RR_7Ba          ACGACAAGAATAAGTGTGTCCTAATCACTGAAACCTCCAGCAAAAATGACAAAGAAATGA  1244
RR_7Bb          ACGACAAGAATAAGTGTGTCCTAATCACTGAAACCTCCAGCAAAAATGACAAAGAAATGA  1244

Brilliance      AAATTTCCCTGGGTATGCACTCTCATCTTTTTTTTCTTTTTC------------------  1721
FL 16.33-8-Hap1 AAATTTCCCTGGGTATGCGCACTCATTTTTTTTTTTCTTTTCTCTTTTCTCAGAGCACTT  1304
RR_7Ba          AAATTTCCCTGGGTATGCGCACTCATTTTTTTTTTTCTTTTCTCTTTTCTCAGAGCACTT  1304
RR_7Bb          AAATTTCCCTGGGTATGCGCACTCATTTTTTTTTTTCTTTTCTCTTTTCTCAGAGCACTT  1304

Brilliance      ------------------------------------------------------------  1721
FL 16.33-8-Hap1 TCATCTCTGTTAATCTATTATTGACAAGTTTGTCAATTCTATATATCAATTTCTAAAATT  1364
RR_7Ba          TCATCTCTGTTAATCTATTATTGACAAGTTTGTCAATTCTATATATCAATTTCTAAAATT  1364
RR_7Bb          TCATCTCTGTTAATCTATTATTGACAAGTTTGTCAATTCTATATATCAATTTCTAAAATT  1364

Brilliance      ------------------------------------------------------------  1721
FL 16.33-8-Hap1 GTCGACTTTTTAAGTATTATATTTTTTTTATACGAACTCTTTAACTATCGTGTTGGAATT  1424
RR_7Ba          GTCGACTTTTTAAGTATTATATTTTTTTTATACGAACTCTTTAACTATCGTGTTGGAATT  1424
RR_7Bb          GTCGACTTTTTAAGTATTATATTTTTTTTATACGAACTCTTTAACTATCGTGTTGGAATT  1424

Brilliance      --------------------------------------TTTTTTTAAATAACACCTCTGC  1743
FL 16.33-8-Hap1 TATATTTACCTTAAAAATTGATAAATTTTAACAGTTTTTTTTTTTTTATAACAAATCTGC  1484
RR_7Ba          TATATTTACCTTAAAAATTGATAAATTTTAACAGTTTTTTTTTTTTTATAACAAATCTGC  1484
```

```
RR_7Bb             TATATTTACCTTAAAAATTGATAAATTTTAACAGTTTTTTTTTTTTTATAACAAATCTGC  1484

Brilliance         TAGAGATGCTC----AATAGTCAAGTTGTAGGAATGGATTTGAGTACGTTTGGGCTGATC  1799
FL 16.33-8-Hap1    TGGAGATGCTCCTAATATAGTCAAGTTGTAGGAGTGGATTTGAGTACG-TTGTGTTGATC  1543
RR_7Ba             TGGAGATGCTCCTAATATAGTCAAGTTGTAGGAGTGGATTTGAGTACG-TTGTGTTGATC  1543
RR_7Bb             TGGAGATGCTCCTAATATAGTCAAGTTGTAGGAGTGGATTTGAGTACG-TTGTGTTGATC  1543

Brilliance         CGATCTTTTGAATTTTTGCAGGTGTCTCTTTGAGCTTCCTAGTCATATTGATTATAACAT  1859
FL 16.33-8-Hap1    CGTTCTTTTGAATTTTTGTAGGTGTCTCTTTGACCTTCCTAGTCATACTGATTATAACTT  1603
RR_7Ba             CGTTCTTTTGAATTTTTGTAGGTGTCTCTTTGACCTTCCTAGTCATACTGATTATAACTT  1603
RR_7Bb             CGTTCTTTTGAATTTTTGTAGGTGTCTCTTTGACCTTCCTAGTCATACTGATTATAACTT  1603

Brilliance         TTTCGATATACTGCGTAATGAAAAGAAGAAAGTTCAAGCAACTCTGCGACAAGTACTACA  1919
FL 16.33-8-Hap1    TTTCGATATACTGCGTACTGAAAAGAAGAAAGTTCAAGCATCTCTGTGACAGGTACTACA  1663
RR_7Ba             TTTCGATATACTGCGTACTGAAAAGAAGAAAGTTCAAGCATCTCTGTGACAGGTACTACA  1663
RR_7Bb             TTTCGATATACTGCGTACTGAAAAGAAGAAAGTTCAAGCATCTCTGTGACAGGTACTACA  1663

Brilliance         AAGACAATGGGGGCTTCTTGTTACCACAAGAAATGCAAAAGTACAGAGGGTCTCAGGCAC  1979
FL 16.33-8-Hap1    AAGACAATGGGGGCTTCTTGTTACCACAAGAAATGCAGAAGTACAAAGGGTCTCAGGCAC  1723
RR_7Ba             AAGACAATGGGGGCTTCTTGTTACCACAAGAAATGCAGAAGTACAAAGGGTCTCAGGCAC  1723
RR_7Bb             AAGACAATGGGGGCTTCTTGTTACCACAAGAAATGCAGAAGTACAAAGGGTCTCAGGCAC  1723

Brilliance         CCAGAATCTTTAAATTAGAAGAGCTTAACAAGGCAACCAACAAGTTTGATGTCACTAATA  2039
FL 16.33-8-Hap1    CCAGAATCTTTAAATTAGAAGAGCTTAACAAGGCAACCAACAAGTTTGATCCCCATGAAA  1783
RR_7Ba             CCAGAATCTTTAAATTAGAAGAGCTTAACAAGGCAACCAACAAGTTTGATCCCCATGAAA  1783
RR_7Bb             CCAGAATCTTTAAATTAGAAGAGCTTAACAAGGCAACCAACAAGTTTGATCCCCATGAAA  1783

Brilliance         TAATTGGAGAGGGAGGCTTTGGATTGGTTTACAAGGGAACATTGCCGGACAACAAGCAGG  2099
FL 16.33-8-Hap1    TAATTGGAGAAGGAGGCTTTGGATTGGTTTACAAGGGAACATTGCCGGACAACAAGCAGG  1843
RR_7Ba             TAATTGGAGAAGGAGGCTTTGGATTGGTTTACAAGGGAACATTGCCGGACAACAAGCAGG  1843
RR_7Bb             TAATTGGAGAAGGAGGCTTTGGATTGGTTTACAAGGGAACATTGCCGGACAACAAGCAGG  1843

Brilliance         TGGTTGCCATAAAGAAGTCAAAAACTGATGCTCCCACCATGACCCCTGAAAATAGGATCG  2159
FL 16.33-8-Hap1    AGGTTGCCATAAAGAAGTCAAAAACTGATGCTCCCACCATGACCCCCGAAAGTAGGATCG  1903
RR_7Ba             AGGTTGCCATAAAGAAGTCAAAAACTGATGCTCCCACCATGACCCCCGAAAGTAGGATCG  1903
RR_7Bb             AGGTTGCCATAAAGAAGTCAAAAACTGATGCTCCCACCATGACCCCCGAAAGTAGGATCG  1903

Brilliance         CCCACACTAAGCAGTTCATTAATGAGATGATTGTTCTCTCTGGAATCAAGCATAGAAATG  2219
FL 16.33-8-Hap1    CCCACACTAAGCAGTTCATTAATGAGATGATTGTTCTCTCTGGAATCAAGCATAGAAATG  1963
RR_7Ba             CCCACACTAAGCAGTTCATTAATGAGATGATTGTTCTCTCTGGAATCAAGCATAGAAATG  1963
RR_7Bb             CCCACACTAAGCAGTTCATTAATGAGATGATTGTTCTCTCTGGAATCAAGCATAGAAATG  1963

Brilliance         TCGTGAGGCTCTTAGGTTGTTGTTTGGAGACCAAAACGCCTATATTAGTGTACGAATTCG  2279
FL 16.33-8-Hap1    TCGTGAGGCTCTTAGGTTGTTGTTTGGAGACCAAAACGCCTATATTAGTGTACGAGTTTG  2023
RR_7Ba             TCGTGAGGCTCTTAGGTTGTTGTTTGGAGACCAAAACGCCTATATTAGTGTACGAGTTTG  2023
RR_7Bb             TCGTGAGGCTCTTAGGTTGTTGTTTGGAGACCAAAACGCCTATATTAGTGTACGAGTTTG  2023

Brilliance         TCAGCAATGGCACACTTTATGAACACATTCATAAAAAGAAAGGCCAAGGACCTCTTTCCT  2339
FL 16.33-8-Hap1    TCTGCAATGGCACACTTTATGAGCACATTCATAAACAGAAAGGCAAAGGACCTCTTTCCT  2083
RR_7Ba             TCTGCAATGGCACACTTTATGAGCACATTCATAAACAGAAAGGCAAAGGACCTCTTTCCT  2083
RR_7Bb             TCTGCAATGGCACACTTTATGAGCACATTCATAAACAGAAAGGCAAAGGACCTCTTTCCT  2083

Brilliance         TTGAGCAACGAGTGAAGATAGCAGCAGAAACAGCAGGATCCCTAGCATACTTGCACTACA  2399
FL 16.33-8-Hap1    TTAGGCAACGAGTGAAGATAGCAGCAGAAACAGCAGGATCCCTAGCATACTTGCACTACA  2143
RR_7Ba             TTAGGCAACGAGTGAAGATAGCAGCAGAAACAGCAGGATCCCTAGCATACTTGCACTACA  2143
RR_7Bb             TTAGGCAACGAGTGAAGATAGCAGCAGAAACAGCAGGATCCCTAGCATACTTGCACTACA  2143

Brilliance         GCGCTTCTTCCACGCCAATTCTGCATCGAGATGTCAAGGCATCTAACATCCTACTGGATG  2459
FL 16.33-8-Hap1    ACGCTTCTTCCACGCCAATTCTGCATCGAGACGTAAAGGCATCTAACATCCTACTGGATG  2203
RR_7Ba             ACGCTTCTTCCACGCCAATTCTGCATCGAGACGTAAAGGCATCTAACATCCTACTGGATG  2203
RR_7Bb             ACGCTTCTTCCACGCCAATTCTGCATCGAGACGTAAAGGCATCTAACATCCTACTGGATG  2203

Brilliance         AAAATTGCATAGCCAAAGTATCAGACTTTGGAGCTTCAAAATTGGTTCCTGAAGATGAAA  2519
FL 16.33-8-Hap1    AGAATTGCACAGCCAAAATATCAGACTTTGGAGCTTCAAAATTGGTTCCCGAAGATGAAA  2263
RR_7Ba             AGAATTGCACAGCCAAAATATCAGACTTTGGAGCTTCAAAATTGGTTCCCGAAGATGAAA  2263
RR_7Bb             AGAATTGCACAGCCAAAATATCAGACTTTGGAGCTTCAAAATTGGTTCCCGAAGATGAAA  2263

Brilliance         ATACTCAAATGGCTACTTTAGTGCAAGGGACGCTAGGGTACTTGGACCCTGAATATCTTC  2579
FL 16.33-8-Hap1    ACACTCAACTGGCTACTTTGGTGCAAGGGACGCTAGGGTACTTGGATCCTCAATATATTC  2323
RR_7Ba             ACACTCAACTGGCTACTTTGGTGCAAGGGACGCTAGGGTACTTGGATCCTCAATATATTC  2323
RR_7Bb             ACACTCAACTGGCTACTTTGGTGCAAGGGACGCTAGGGTACTTGGATCCTCAATATATTC  2323

Brilliance         AGTCACACGCACTGACGGAGAAGAGTGATGTCTATAGTTTCGGGGTTGTCCTTGTGGAGC  2639
FL 16.33-8-Hap1    AAACACACACACTGACGGAGAAGAGTGATGTCTACAGTTTCGGGGTTGTCCTTGTGGAAC  2383
RR_7Ba             AAACACACACACTGACGGAGAAGAGTGATGTCTACAGTTTCGGGGTTGTCCTTGTGGAAC  2383
RR_7Bb             AAACACACACACTGACGGAGAAGAGTGATGTCTACAGTTTCGGGGTTGTCCTTGTGGAAC  2383

Brilliance         TAATAACAAGTCAAGTGGCAAT---TTCTAACAAGAAGCTTGAGGCAGAGAAAAACCTAG  2696
FL 16.33-8-Hap1    TGATAACGAGTCAAGCGGCAATTAATTCTAACAAGAAGCTTGAGGCAGAGAAAAACCTAG  2443
RR_7Ba             TGATAACGAGTCAAGCGGCAATTAATTCTAACAAGAAGCTTGAGGCAGAGAAAAACCTAG  2443
RR_7Bb             TGATAACGAGTCAAGCGGCAATTAATTCTAACAAGAAGCTTGAGGCAGAGAAAAACCTAG  2443
```

```
Brilliance       CAAACTGTTTTTTAAAGTCGGTGGCAGATAATAACTTGGATCAGATTCTTGATCATGAAA  2756
FL 16.33-8-Hap1  CCAATTGTTTTTTAAAGTCGGTCGAAGATAATCACTTGGATCAGATTCTTGATCATGAAA  2503
RR_7Ba           CCAATTGTTTTTTAAAGTCGGTCGAAGATAATCACTTGGATCAGATTCTTGATCATGAAA  2503
RR_7Bb           CCAATTGTTTTTTAAAGTCGGTCGAAGATAATCACTTGGATCAGATTCTTGATCATGAAA  2503

Brilliance       TTATCAAAGAAGAAAGCTTCGAAATAGCCGAACAAGTAGCTCATCTCGCCAAAAGATGTT  2816
FL 16.33-8-Hap1  TTATCA---AAGATAGCTTAGAAATAGCTGAACAAGTAGCCCATCTCGCCAAAAGATGTT  2560
RR_7Ba           TTATCA---AAGATAGCTTAGAAATAGCTGAACAAGTAGCCCATCTCGCCAAAAGATGTT  2560
RR_7Bb           TTATCA---AAGATAGCTTAGAAATAGCTGAACAAGTAGCCCATCTCGCCAAAAGATGTT  2560

Brilliance       TAAGTTTGAAAGGAGAAGATAGGCCTACCATGGAAGAAGTAGAAACAGAGCTAAAGGTGA  2876
FL 16.33-8-Hap1  TGAGTTTGAAAGGAGATGATAGGCCTGCCATGAGAGAAGTAGAAACGGAGCTAGTGGCAA  2620
RR_7Ba           TGAGTTTGAAAGGAGATGATAGGCCTGCCATGAGAGAAGTAGAAACGGAGCTAGTGGCAA  2620
RR_7Bb           TGAGTTTGAAAGGAGATGATAGGCCTGCCATGAGAGAAGTAGAAACGGAGCTAGTGGCAA  2620

Brilliance       TATTGGCAGTTATGGCAAAGCATCCAGGAGGAAAGCCCGACTCCTCCCCCAAAGAGACAG  2936
FL 16.33-8-Hap1  TATTGGCAGTTATGGAAAAGCGTCCCGGAGGAAAGCCTGACTCCTCCCCCAAAGAGACCG  2680
RR_7Ba           TATTGGCAGTTATGGAAAAGCGTCCCGGAGGAAAGCCTGACTCCTCCCCCAAAGAGACCG  2680
RR_7Bb           TATTGGCAGTTATGGAAAAGCGTCCCGGAGGAAAGCCTGACTCCTCCCCCAAAGAGACCG  2680

Brilliance       ATTACTTGCTTGCAGCGTCACCTTCAAACGCTTACGTTGTTGATGTTAGAAGTGATGAAG  2996
FL 16.33-8-Hap1  ATTACTTGCTTGCAGCGTCACCTTCAAATGCTTTCGTTGTTGATGTTAGAAGTGATGAAG  2740
RR_7Ba           ATTACTTGCTTGCAGCGTCACCTTCAAATGCTTTCGTTGTTGATGTTAGAAGTGATGAAG  2740
RR_7Bb           ATTACTTGCTTGCAGCGTCACCTTCAAATGCTTTCGTTGTTGATGTTAGAAGTGATGAAG  2740

Brilliance       GTGAAGTCATAACTAGCATAGATTATGACAAGAGCATGCAGAATCAAGCCCAGATGATGA  3056
FL 16.33-8-Hap1  GTGAACTCACAACCAGCATAGATTATGACCAGAGCATGCAGAATCAATCCCAGATGACGA  2800
RR_7Ba           GTGAACTCACAACCAGCATAGATTATGACCAGAGCATGCAGAATCAATCCCAGATGACGA  2800
RR_7Bb           GTGAACTCACAACCAGCATAGATTATGACCAGAGCATGCAGAATCAATCCCAGATGACGA  2800

Brilliance       AGCATTATGATGGTGGGAGATAG                                       3079
FL 16.33-8-Hap1  GGCCTTATGATAGTGGGAGATAG                                       2823
RR_7Ba           GGCCTTATGATAGTGGGAGATAG                                       2823
RR_7Bb           GGCCTTATGATAGTGGGAGATAG                                       2823
```

**CDS of WAK**

```
Brilliance       ATGGCCTTACATGGGAGAATGCTTCTTTTGCAACTCTCTTTGGCCGCAGTGCTATTAGCA    60
FL 16.33-8-Hap1  ATGGCCTTACATGGGAGAATGCTTCTTTTGCAACTCTCTTTGGCCGCAGTGCTATTAGCA    60
RR_7Ba           ATGGCCTTACATGGGAGAATGCTTCTTTTGCAACTCTCTTTGGCCGCAGTGCTATTAGCA    60
RR_7Bb           ATGGCCTTACATGGGAGAATGCTTCTTTTGCAACTCTCTTTGGCCGCAGTGCTATTAGCA    60

Brilliance       ACGACAATGCTAGCAGCTGCTCAAGCCCCGCCGCCTGACTGCAATGAGAGCTGCGGTGGT   120
FL 16.33-8-Hap1  ACGACAATGCTAGCAGCTGCTCAAGCCTCACCGCCTGTCTGCAATGAGAGCTGCGGTGGT   120
RR_7Ba           ACGACAATGCTAGCAGCTGCTCAAGCCTCACCGCCTGTCTGCAATGAGAGCTGCGGTGGT   120
RR_7Bb           ACGACAATGCTAGCAGCTGCTCAAGCCTCACCGCCTGTCTGCAATGAGAGCTGCGGTGGT   120

Brilliance       GTCTCAGTTCCATATCCATTTGGTCTCACTGAGGGTTGTTACCTACATGTGCCGGGACAA   180
FL 16.33-8-Hap1  GTCTCAGTTCCATATCCATTTGGTCTCACTGATGGTTGTTACCTACATGTGCCGGGACAA   180
RR_7Ba           GTCTCAGTTCCATATCCATTTGGTCTCACTGATGGTTGTTACCTACATGTGCCGGGACAA   180
RR_7Bb           GTCTCAGTTCCATATCCATTTGGTCTCACTGATGGTTGTTACCTACATGTGCCGGGACAA   180

Brilliance       GATTCTCAGCAGCCATTCAAGATCACTTGCAACACCACCACCTCGCAACCATCCCTACAG   240
FL 16.33-8-Hap1  GATTCTCAGCAGCCATTCAAGATCACTTGCAACACCACCACCTCGCAACCATCCCTACAG   240
RR_7Ba           GATTCTCAGCAGCCATTCAAGATCACTTGCAACACCACCACCTCGCAACCATCCCTACAG   240
RR_7Bb           GATTCTCAGCAGCCATTCAAGATCACTTGCAACACCACCACCTCGCAACCATCCCTACAG   240

Brilliance       TTCTCGGACAGCGATAACTTCCCCACAAATATTGCCAACATTTTTGTGGAAGAGAGTGCG   300
FL 16.33-8-Hap1  TTCTCGGACAGCGATAACTTCCCCACAAATATTGCCAACATTTTTTTGGAAGAGAGTGCG   300
RR_7Ba           TTCTCGGACAGCGATAACTTCCCCACAAATATTGCCAACATTTTTTTGGAAGAGAGTGCG   300
RR_7Bb           TTCTCGGACAGCGATAACTTCCCCACAAATATTGCCAACATTTTTTTGGAAGAGAGTGCG   300

Brilliance       CTGCAAGTAATGATGGCTCCGTCCTATAACTGCTACACCAACAATACATACGACTCCTCC   360
FL 16.33-8-Hap1  CTGCAAGTAATGATGGCCCCGTCCTATAACTGCTACACCAACAATACATACAACCCCTCC   360
RR_7Ba           CTGCAAGTAATGATGGCCCCGTCCTATAACTGCTACACCAACAATACATACAACCCCTCC   360
RR_7Bb           CTGCAAGTAATGATGGCCCCGTCCTATAACTGCTACACCAACAATACATACAACCCCTCC   360

Brilliance       ATGGACAAAGTCATGGACCTCAATCTTCCGCCTTCTTTCACCCTCTCCGACAGAAACAAG   420
FL 16.33-8-Hap1  ATGGACAAAGTCATGGACCTCAATCTTCCGCCTTCTTACACCCTCTCCAACAGAAACAAG   420
RR_7Ba           ATGGACAAAGTCATGGACCTCAATCTTCCGCCTTCTTACACCCTCTCCAACAGAAACAAG   420
RR_7Bb           ATGGACAAAGTCATGGACCTCAATCTTCCGCCTTCTTACACCCTCTCCAACAGAAACAAG   420

Brilliance       GTTTATAACCTTGGGTGCAACAAAGTAACGAAGCTCATAGGTTATCCTCTTGAAGTTACA   480
FL 16.33-8-Hap1  GTTTATAACCTTGGGTGCAACAAAGTAACGAAACTCATAGGTTATCCTCTTGAAGTTACA   480
RR_7Ba           GTTTATAACCTTGGGTGCAACAAAGTAACGAAACTCATAGGTTATCCTCTTGAAGTTACA   480
RR_7Bb           GTTTATAACCTTGGGTGCAACAAAGTAACGAAACTCATAGGTTATCCTCTTGAAGTTACA   480

Brilliance       GACCCTGACCAGCTCTTTGAAGTAGGGTTCAGCGGTGTAAGCTTGTGCCAAGATGAATTT   540
FL 16.33-8-Hap1  GACCCTGAGCAGCTCTTTCAAGTACGGTTCAGCGGTGTAAGCTTGTGCCAAGATGAATTC   540
RR_7Ba           GACCCTGAGCAGCTCTTTCAAGTACGGTTCAGCGGTGTAAGCTTGTGCCAAGATGAATTC   540
```

```
RR_7Bb            GACCCTGAGCAGCTCTTTCAAGTACGGTTCAGCGGTGTAAGCTTGTGCCAAGATGAATTC   540

Brilliance        GGCAAAAGATTCCCCGACACTTGCACCGGATTTGGGTGCTCCGTAAATTCTATCCCTATC   600
FL 16.33-8-Hap1   GGCAAGAGATTCCCCGACACTTGCACCGGATTCGGGTGCTCCGTAAATTCTATCCCTAGC   600
RR_7Ba            GGCAAGAGATTCCCCGACACTTGCACCGGATTCGGGTGCTCCGTAAATTCTATCCCTAGC   600
RR_7Bb            GGCAAGAGATTCCCCGACACTTGCACCGGATTCGGGTGCTCCGTAAATTCTATCCCTAGC   600

Brilliance        GGACTGCAGAATATTACGTTGGGTGTGTGGACACTCGGTACCAGCCTCGGAACAACGGCG   660
FL 16.33-8-Hap1   GGACTGCAGAATATTACGGTGGATGTGTGGACACTCGGTACCGGCCTCGGAAAAACGGAG   660
RR_7Ba            GGACTGCAGAATATTACGGTGGATGTGTGGACACTCGGTACCGGCCTCGGAAAAACGGAG   660
RR_7Bb            GGACTGCAGAATATTACGGTGGATGTGTGGACACTCGGTACCGGCCTCGGAAAAACGGAG   660

Brilliance        CAGTGGGGCCTGAGCTACCCTTGCAGTTACGGTTTCGTTGTGGATGAACGCAACTTCACA   720
FL 16.33-8-Hap1   CAGTGGGGACTGAGCTACCCTTGCAGTTACGGCTTCGTTGTGGATGAACGCAACTTCACA   720
RR_7Ba            CAGTGGGGACTGAGCTACCCTTGCAGTTACGGCTTCGTTGTGGATGAACGCAACTTCACA   720
RR_7Bb            CAGTGGGGACTGAGCTACCCTTGCAGTTACGGCTTCGTTGTGGATGAACGCAACTTCACA   720

Brilliance        TTCGCCGGGAATCAAAGTTTTGATGCACTGAGGTCCGGCGAAATAAAGAAGCTTCCGGTG   780
FL 16.33-8-Hap1   TTCGCCGGGAATCAAAGTTTTGTTGCACTGAGGTCCGGCGCAATAAAGAAGCTTCCGGTG   780
RR_7Ba            TTCGCCGGGAATCAAAGTTTTGTTGCACTGAGGTCCGGCGCAATAAAGAAGCTTCCGGTG   780
RR_7Bb            TTCGCCGGGAATCAAAGTTTTGTTGCACTGAGGTCCGGCGCAATAAAGAAGCTTCCGGTG   780

Brilliance        CTTGCTAATTGGGCAATTGGGAATGACAACTGCGAAGCAGCAAAGAAGAAGAACGAGACT   840
FL 16.33-8-Hap1   CTTGCTAATTGGGCGATTGGGAATGACAACTGCGAAGCAGCAAAGAAGAAGAACGAGACT   840
RR_7Ba            CTTGCTAATTGGGCGATTGGGAATGACAACTGCGAAGCAGCAAAGAAGAAGAACGAGACT   840
RR_7Bb            CTTGCTAATTGGGCGATTGGGAATGACAACTGCGAAGCAGCAAAGAAGAAGAACGAGACT   840

Brilliance        GCTTTTGCATGCAAGACTGTGAATTCCAAGTGTGTCGACCGTGCTGGTGGTGGTTACTTT   900
FL 16.33-8-Hap1   GCTTTTGCATGCAAGACTGTGAATTCCAAGTGTGTCGACCGTGCTGGTGGTGGTTACTTT   900
RR_7Ba            GCTTTTGCATGCAAGACTGTGAATTCCAAGTGTGTCGACCGTGCTGGTGGTGGTTACTTT   900
RR_7Bb            GCTTTTGCATGCAAGACTGTGAATTCCAAGTGTGTCGACCGTGCTGGTGGTGGTTACTTT   900

Brilliance        TGCCAGTGTGAGGATGGTTACGAAGGGAACCCATACCTACTCGATAATTGCCTAGATATC   960
FL 16.33-8-Hap1   TGCCAGTGTGAGGATGGTTACGAAGGGAACCCATACCTCCCCGATAATTGCCTAGATATC   960
RR_7Ba            TGCCAGTGTGAGGATGGTTACGAAGGGAACCCATACCTCCCCGATAATTGCCTAGATATC   960
RR_7Bb            TGCCAGTGTGAGGATGGTTACGAAGGGAACCCATACCTCCCCGATAATTGCCTAGATATC   960

Brilliance        AATGAGTGCAATAAGAACTCAACCCTTTGCAGCGGTCCTGCAACTTGCATAAACTCAATT  1020
FL 16.33-8-Hap1   AATGAGTGC---AAGCATTCAACCCTTTGCAGTGGTCCTGCAACTTGCATAAACTCAATT  1017
RR_7Ba            AATGAGTGC---AAGCATTCAACCCTTTGCAGTGGTCCTGCAACTTGCATAAACTCAATT  1017
RR_7Bb            AATGAGTGC---AAGCATTCAACCCTTTGCAGTGGTCCTGCAACTTGCATAAACTCAATT  1017

Brilliance        GGAGGTTACACCTGTAAATGTCACAAGGGCTATAGAAATGATGACCACGACAAGAATAAG  1080
FL 16.33-8-Hap1   GGAAGTTACACCTGTAAATGTCACAAGGGCCATAGAAATGATGACCACGACAAGAATAAG  1077
RR_7Ba            GGAAGTTACACCTGTAAATGTCACAAGGGCCATAGAAATGATGACCACGACAAGAATAAG  1077
RR_7Bb            GGAAGTTACACCTGTAAATGTCACAAGGGCCATAGAAATGATGACCACGACAAGAATAAG  1077

Brilliance        TGTGTCCAAATCACTGAAACCTCCAGCAAAAATGACAAAGAAATGAAAATTTCCCTGGGT  1140
FL 16.33-8-Hap1   TGTGTCCTAATCACTGAAACCTCCAGCAAAAATGACAAAGAAATGAAAATTTCCCTGGGT  1137
RR_7Ba            TGTGTCCTAATCACTGAAACCTCCAGCAAAAATGACAAAGAAATGAAAATTTCCCTGGGT  1137
RR_7Bb            TGTGTCCTAATCACTGAAACCTCCAGCAAAAATGACAAAGAAATGAAAATTTCCCTGGGT  1137

Brilliance        GTCTCTTTGAGCTTCCTAGTCATATTGATTATAACATTTTCGATATACTGCGTAATGAAA  1200
FL 16.33-8-Hap1   GTCTCTTTGACCTTCCTAGTCATACTGATTATAACTTTTTCGATATACTGCGTACTGAAA  1197
RR_7Ba            GTCTCTTTGACCTTCCTAGTCATACTGATTATAACTTTTTCGATATACTGCGTACTGAAA  1197
RR_7Bb            GTCTCTTTGACCTTCCTAGTCATACTGATTATAACTTTTTCGATATACTGCGTACTGAAA  1197

Brilliance        AGAAGAAAGTTCAAGCAACTCTGCGACAAGTACTACAAAGACAATGGGGGCTTCTTGTTA  1260
FL 16.33-8-Hap1   AGAAGAAAGTTCAAGCATCTCTGTGACAGGTACTACAAAGACAATGGGGGCTTCTTGTTA  1257
RR_7Ba            AGAAGAAAGTTCAAGCATCTCTGTGACAGGTACTACAAAGACAATGGGGGCTTCTTGTTA  1257
RR_7Bb            AGAAGAAAGTTCAAGCATCTCTGTGACAGGTACTACAAAGACAATGGGGGCTTCTTGTTA  1257

Brilliance        CCACAAGAAATGCAAAAGTACAGAGGGTCTCAGGCACCCAGAATCTTTAAATTAGAAGAG  1320
FL 16.33-8-Hap1   CCACAAGAAATGCAGAAGTACAAAGGGTCTCAGGCACCCAGAATCTTTAAATTAGAAGAG  1317
RR_7Ba            CCACAAGAAATGCAGAAGTACAAAGGGTCTCAGGCACCCAGAATCTTTAAATTAGAAGAG  1317
RR_7Bb            CCACAAGAAATGCAGAAGTACAAAGGGTCTCAGGCACCCAGAATCTTTAAATTAGAAGAG  1317

Brilliance        CTTAACAAGGCAACCAACAAGTTTGATGTCACTAATATAATTGGAGAGGGAGGCTTTGGA  1380
FL 16.33-8-Hap1   CTTAACAAGGCAACCAACAAGTTTGATCCCCATGAAATAATTGGAGAAGGAGGCTTTGGA  1377
RR_7Ba            CTTAACAAGGCAACCAACAAGTTTGATCCCCATGAAATAATTGGAGAAGGAGGCTTTGGA  1377
RR_7Bb            CTTAACAAGGCAACCAACAAGTTTGATCCCCATGAAATAATTGGAGAAGGAGGCTTTGGA  1377

Brilliance        TTGGTTTACAAGGGAACATTGCCGGACAACAAGCAGGTGGTTGCCATAAAGAAGTCAAAA  1440
FL 16.33-8-Hap1   TTGGTTTACAAGGGAACATTGCCGGACAACAAGCAGGAGGTTGCCATAAAGAAGTCAAAA  1437
RR_7Ba            TTGGTTTACAAGGGAACATTGCCGGACAACAAGCAGGAGGTTGCCATAAAGAAGTCAAAA  1437
RR_7Bb            TTGGTTTACAAGGGAACATTGCCGGACAACAAGCAGGAGGTTGCCATAAAGAAGTCAAAA  1437

Brilliance        ACTGATGCTCCCACCATGACCCCTGAAAATAGGATCGCCCACACTAAGCAGTTCATTAAT  1500
FL 16.33-8-Hap1   ACTGATGCTCCCACCATGACCCCCGAAAGTAGGATCGCCCACACTAAGCAGTTCATTAAT  1497
RR_7Ba            ACTGATGCTCCCACCATGACCCCCGAAAGTAGGATCGCCCACACTAAGCAGTTCATTAAT  1497
RR_7Bb            ACTGATGCTCCCACCATGACCCCCGAAAGTAGGATCGCCCACACTAAGCAGTTCATTAAT  1497
```

```
Brilliance        GAGATGATTGTTCTCTCTGGAATCAAGCATAGAAATGTCGTGAGGCTCTTAGGTTGTTGT  1560
FL 16.33-8-Hap1   GAGATGATTGTTCTCTCTGGAATCAAGCATAGAAATGTCGTGAGGCTCTTAGGTTGTTGT  1557
RR_7Ba            GAGATGATTGTTCTCTCTGGAATCAAGCATAGAAATGTCGTGAGGCTCTTAGGTTGTTGT  1557
RR_7Bb            GAGATGATTGTTCTCTCTGGAATCAAGCATAGAAATGTCGTGAGGCTCTTAGGTTGTTGT  1557

Brilliance        TTGGAGACCAAAACGCCTATATTAGTGTACGAATTCGTCAGCAATGGCACACTTTATGAA  1620
FL 16.33-8-Hap1   TTGGAGACCAAAACGCCTATATTAGTGTACGAGTTTGTCTGCAATGGCACACTTTATGAG  1617
RR_7Ba            TTGGAGACCAAAACGCCTATATTAGTGTACGAGTTTGTCTGCAATGGCACACTTTATGAG  1617
RR_7Bb            TTGGAGACCAAAACGCCTATATTAGTGTACGAGTTTGTCTGCAATGGCACACTTTATGAG  1617

Brilliance        CACATTCATAAAAGAAAGGCCAAGGACCTCTTTCCTTTGAGCAACGAGTGAAGATAGCA  1680
FL 16.33-8-Hap1   CACATTCATAAACAGAAAGGCAAAGGACCTCTTTCCTTTAGGCAACGAGTGAAGATAGCA  1677
RR_7Ba            CACATTCATAAACAGAAAGGCAAAGGACCTCTTTCCTTTAGGCAACGAGTGAAGATAGCA  1677
RR_7Bb            CACATTCATAAACAGAAAGGCAAAGGACCTCTTTCCTTTAGGCAACGAGTGAAGATAGCA  1677

Brilliance        GCAGAAACAGCAGGATCCCTAGCATACTTGCACTACAGCGCTTCTTCCACGCCAATTCTG  1740
FL 16.33-8-Hap1   GCAGAAACAGCAGGATCCCTAGCATACTTGCACTACAACGCTTCTTCCACGCCAATTCTG  1737
RR_7Ba            GCAGAAACAGCAGGATCCCTAGCATACTTGCACTACAACGCTTCTTCCACGCCAATTCTG  1737
RR_7Bb            GCAGAAACAGCAGGATCCCTAGCATACTTGCACTACAACGCTTCTTCCACGCCAATTCTG  1737

Brilliance        CATCGAGATGTCAAGGCATCTAACATCCTACTGGATGAAAATTGCATAGCCAAAGTATCA  1800
FL 16.33-8-Hap1   CATCGAGACGTAAAGGCATCTAACATCCTACTGGATGAGAATTGCACAGCCAAAATATCA  1797
RR_7Ba            CATCGAGACGTAAAGGCATCTAACATCCTACTGGATGAGAATTGCACAGCCAAAATATCA  1797
RR_7Bb            CATCGAGACGTAAAGGCATCTAACATCCTACTGGATGAGAATTGCACAGCCAAAATATCA  1797

Brilliance        GACTTTGGAGCTTCAAAATTGGTTCCTGAAGATGAAAATACTCAAATGGCTACTTTAGTG  1860
FL 16.33-8-Hap1   GACTTTGGAGCTTCAAAATTGGTTCCCGAAGATGAAAACACTCAACTGGCTACTTTGGTG  1857
RR_7Ba            GACTTTGGAGCTTCAAAATTGGTTCCCGAAGATGAAAACACTCAACTGGCTACTTTGGTG  1857
RR_7Bb            GACTTTGGAGCTTCAAAATTGGTTCCCGAAGATGAAAACACTCAACTGGCTACTTTGGTG  1857

Brilliance        CAAGGGACGCTAGGGTACTTGGACCCTGAATATCTTCAGTCACACGCACTGACGGAGAAG  1920
FL 16.33-8-Hap1   CAAGGGACGCTAGGGTACTTGGATCCTCAATATATTCAAACACACACACTGACGGAGAAG  1917
RR_7Ba            CAAGGGACGCTAGGGTACTTGGATCCTCAATATATTCAAACACACACACTGACGGAGAAG  1917
RR_7Bb            CAAGGGACGCTAGGGTACTTGGATCCTCAATATATTCAAACACACACACTGACGGAGAAG  1917

Brilliance        AGTGATGTCTATAGTTTCGGGGTTGTCCTTGTGGAGCTAATAACAAGTCAAGTGGCAAT-  1979
FL 16.33-8-Hap1   AGTGATGTCTACAGTTTCGGGGTTGTCCTTGTGGAACTGATAACGAGTCAAGCGGCAATT  1977
RR_7Ba            AGTGATGTCTACAGTTTCGGGGTTGTCCTTGTGGAACTGATAACGAGTCAAGCGGCAATT  1977
RR_7Bb            AGTGATGTCTACAGTTTCGGGGTTGTCCTTGTGGAACTGATAACGAGTCAAGCGGCAATT  1977

Brilliance        --TTCTAACAAGAAGCTTGAGGCAGAGAAAAACCTAGCAAACTGTTTTTTAAAGTCGGTG  2037
FL 16.33-8-Hap1   AATTCTAACAAGAAGCTTGAGGCAGAGAAAAACCTAGCCAATTGTTTTTTAAAGTCGGTC  2037
RR_7Ba            AATTCTAACAAGAAGCTTGAGGCAGAGAAAAACCTAGCCAATTGTTTTTTAAAGTCGGTC  2037
RR_7Bb            AATTCTAACAAGAAGCTTGAGGCAGAGAAAAACCTAGCCAATTGTTTTTTAAAGTCGGTC  2037

Brilliance        GCAGATAATAACTTGGATCAGATTCTTGATCATGAAATTATCAAAGAAGAAAGCTTCGAA  2097
FL 16.33-8-Hap1   GAAGATAATCACTTGGATCAGATTCTTGATCATGAAATTATCA---AAGATAGCTTAGAA  2094
RR_7Ba            GAAGATAATCACTTGGATCAGATTCTTGATCATGAAATTATCA---AAGATAGCTTAGAA  2094
RR_7Bb            GAAGATAATCACTTGGATCAGATTCTTGATCATGAAATTATCA---AAGATAGCTTAGAA  2094

Brilliance        ATAGCCGAACAAGTAGCTCATCTCGCCAAAAGATGTTTAAGTTTGAAAGGAGAAGATAGG  2157
FL 16.33-8-Hap1   ATAGCTGAACAAGTAGCCCATCTCGCCAAAAGATGTTTGAGTTTGAAAGGAGATGATAGG  2154
RR_7Ba            ATAGCTGAACAAGTAGCCCATCTCGCCAAAAGATGTTTGAGTTTGAAAGGAGATGATAGG  2154
RR_7Bb            ATAGCTGAACAAGTAGCCCATCTCGCCAAAAGATGTTTGAGTTTGAAAGGAGATGATAGG  2154

Brilliance        CCTACCATGGAAGAAGTAGAAACAGAGCTAAAGGTGATATTGGCAGTTATGGCAAAGCAT  2217
FL 16.33-8-Hap1   CCTGCCATGAGAGAAGTAGAAACGGAGCTAGTGGCAATATTGGCAGTTATGGAAAAGCGT  2214
RR_7Ba            CCTGCCATGAGAGAAGTAGAAACGGAGCTAGTGGCAATATTGGCAGTTATGGAAAAGCGT  2214
RR_7Bb            CCTGCCATGAGAGAAGTAGAAACGGAGCTAGTGGCAATATTGGCAGTTATGGAAAAGCGT  2214

Brilliance        CCAGGAGGAAAGCCCGACTCCTCCCCCAAAGAGACAGATTACTTGCTTGCAGCGTCACCT  2277
FL 16.33-8-Hap1   CCCGGAGGAAAGCCTGACTCCTCCCCCAAAGAGACCGATTACTTGCTTGCAGCGTCACCT  2274
RR_7Ba            CCCGGAGGAAAGCCTGACTCCTCCCCCAAAGAGACCGATTACTTGCTTGCAGCGTCACCT  2274
RR_7Bb            CCCGGAGGAAAGCCTGACTCCTCCCCCAAAGAGACCGATTACTTGCTTGCAGCGTCACCT  2274

Brilliance        TCAAACGCTTACGTTGTTGATGTTAGAAGTGATGAAGGTGAAGTCATAACTAGCATAGAT  2337
FL 16.33-8-Hap1   TCAAATGCTTTCGTTGTTGATGTTAGAAGTGATGAAGGTGAACTCACAACCAGCATAGAT  2334
RR_7Ba            TCAAATGCTTTCGTTGTTGATGTTAGAAGTGATGAAGGTGAACTCACAACCAGCATAGAT  2334
RR_7Bb            TCAAATGCTTTCGTTGTTGATGTTAGAAGTGATGAAGGTGAACTCACAACCAGCATAGAT  2334

Brilliance        TATGACAAGAGCATGCAGAATCAAGCCCAGATGATGAAGCATTATGATGGTGGGAGATAG  2397
FL 16.33-8-Hap1   TATGACCAGAGCATGCAGAATCAATCCCAGATGACGAGGCCTTATGATAGTGGGAGATAG  2394
RR_7Ba            TATGACCAGAGCATGCAGAATCAATCCCAGATGACGAGGCCTTATGATAGTGGGAGATAG  2394
RR_7Bb            TATGACCAGAGCATGCAGAATCAATCCCAGATGACGAGGCCTTATGATAGTGGGAGATAG  2394
```

**Peptide sequence of WAK**

```
Brilliance        MALHGRMLLLQLSLAAVLLATTMLAAAQAPPPDCNESCGGVSVPYPFGLTEGCYLHVPGQ   60
FL 16.33-8-Hap1   MALHGRMLLLQLSLAAVLLATTMLAAAQASPPVCNESCGGVSVPYPFGLTDGCYLHVPGQ   60
RR_7Ba            MALHGRMLLLQLSLAAVLLATTMLAAAQASPPVCNESCGGVSVPYPFGLTDGCYLHVPGQ   60
```

```
RR_7Bb              MALHGRMLLLQLSLAAVLLATTMLAAAQASPPVCNESCGGVSVPYPFGLTDGCYLHVPGQ    60

Brilliance          DSQQPFKITCNTTTSQPSLQFSDSDNFPTNIANIFVEESALQVMMAPSYNCYTNNTYDSS   120
FL 16.33-8-Hap1     DSQQPFKITCNTTTSQPSLQFSDSDNFPTNIANIFLEESALQVMMAPSYNCYTNNTYNPS   120
RR_7Ba              DSQQPFKITCNTTTSQPSLQFSDSDNFPTNIANIFLEESALQVMMAPSYNCYTNNTYNPS   120
RR_7Bb              DSQQPFKITCNTTTSQPSLQFSDSDNFPTNIANIFLEESALQVMMAPSYNCYTNNTYNPS   120

Brilliance          MDKVMDLNLPPSFTLSDRNKVYNLGCNKVTKLIGYPLEVTDPDQLFEVGFSGVSLCQDEF   180
FL 16.33-8-Hap1     MDKVMDLNLPPSYTLSNRNKVYNLGCNKVTKLIGYPLEVTDPEQLFQVRFSGVSLCQDEF   180
RR_7Ba              MDKVMDLNLPPSYTLSNRNKVYNLGCNKVTKLIGYPLEVTDPEQLFQVRFSGVSLCQDEF   180
RR_7Bb              MDKVMDLNLPPSYTLSNRNKVYNLGCNKVTKLIGYPLEVTDPEQLFQVRFSGVSLCQDEF   180

Brilliance          GKRFPDTCTGFGCSVNSIPIGLQNITLGVWTLGTSLGTTAQWGLSYPCSYGFVVDERNFT   240
FL 16.33-8-Hap1     GKRFPDTCTGFGCSVNSIPSGLQNITVDVWTLGTGLGKTEQWGLSYPCSYGFVVDERNFT   240
RR_7Ba              GKRFPDTCTGFGCSVNSIPSGLQNITVDVWTLGTGLGKTEQWGLSYPCSYGFVVDERNFT   240
RR_7Bb              GKRFPDTCTGFGCSVNSIPSGLQNITVDVWTLGTGLGKTEQWGLSYPCSYGFVVDERNFT   240

Brilliance          FAGNQSFDALRSGEIKKLPVLANWAIGNDNCEAAKKKNETAFACKTVNSKCVDRAGGGYF   300
FL 16.33-8-Hap1     FAGNQSFVALRSGAIKKLPVLANWAIGNDNCEAAKKKNETAFACKTVNSKCVDRAGGGYF   300
RR_7Ba              FAGNQSFVALRSGAIKKLPVLANWAIGNDNCEAAKKKNETAFACKTVNSKCVDRAGGGYF   300
RR_7Bb              FAGNQSFVALRSGAIKKLPVLANWAIGNDNCEAAKKKNETAFACKTVNSKCVDRAGGGYF   300

Brilliance          CQCEDGYEGNPYLLDNCLDINECNKNSTLCSGPATCINSIGGYTCKCHKGYRNDDHDKNK   360
FL 16.33-8-Hap1     CQCEDGYEGNPYLPDNCLDINEC-KHSTLCSGPATCINSIGSYTCKCHKGHRNDDHDKNK   359
RR_7Ba              CQCEDGYEGNPYLPDNCLDINEC-KHSTLCSGPATCINSIGSYTCKCHKGHRNDDHDKNK   359
RR_7Bb              CQCEDGYEGNPYLPDNCLDINEC-KHSTLCSGPATCINSIGSYTCKCHKGHRNDDHDKNK   359

Brilliance          CVQITETSSKNDKEMKISLGVSLSFLVILIITFSIYCVMKRRKFKQLCDKYYKDNGGFLL   420
FL 16.33-8-Hap1     CVLITETSSKNDKEMKISLGVSLTFLVILIITFSIYCVLKRRKFKHLCDRYYKDNGGFLL   419
RR_7Ba              CVLITETSSKNDKEMKISLGVSLTFLVILIITFSIYCVLKRRKFKHLCDRYYKDNGGFLL   419
RR_7Bb              CVLITETSSKNDKEMKISLGVSLTFLVILIITFSIYCVLKRRKFKHLCDRYYKDNGGFLL   419

Brilliance          PQEMQKYRGSQAPRIFKLEELNKATNKFDVTNIIGEGGFGLVYKGTLPDNKQVVAIKKSK   480
FL 16.33-8-Hap1     PQEMQKYKGSQAPRIFKLEELNKATNKFDPHEIIGEGGFGLVYKGTLPDNKQEVAIKKSK   479
RR_7Ba              PQEMQKYKGSQAPRIFKLEELNKATNKFDPHEIIGEGGFGLVYKGTLPDNKQEVAIKKSK   479
RR_7Bb              PQEMQKYKGSQAPRIFKLEELNKATNKFDPHEIIGEGGFGLVYKGTLPDNKQEVAIKKSK   479

Brilliance          TDAPTMTPENRIAHTKQFINEMIVLSGIKHRNVVRLLGCCLETKTPILVYEFVSNGTLYE   540
FL 16.33-8-Hap1     TDAPTMTPESRIAHTKQFINEMIVLSGIKHRNVVRLLGCCLETKTPILVYEFVCNGTLYE   539
RR_7Ba              TDAPTMTPESRIAHTKQFINEMIVLSGIKHRNVVRLLGCCLETKTPILVYEFVCNGTLYE   539
RR_7Bb              TDAPTMTPESRIAHTKQFINEMIVLSGIKHRNVVRLLGCCLETKTPILVYEFVCNGTLYE   539

Brilliance          HIHKKKGQGPLSFEQRVKIAAETAGSLAYLHYSASSTPILHRDVKASNILLDENCIAKVS   600
FL 16.33-8-Hap1     HIHKQKGKGPLSFRQRVKIAAETAGSLAYLHYNASSTPILHRDVKASNILLDENCTAKIS   599
RR_7Ba              HIHKQKGKGPLSFRQRVKIAAETAGSLAYLHYNASSTPILHRDVKASNILLDENCTAKIS   599
RR_7Bb              HIHKQKGKGPLSFRQRVKIAAETAGSLAYLHYNASSTPILHRDVKASNILLDENCTAKIS   599

Brilliance          DFGASKLVPEDENTQMATLVQGTLGYLDPEYLQSHALTEKSDVYSFGVVLVELITSQVAI   660
FL 16.33-8-Hap1     DFGASKLVPEDENTQLATLVQGTLGYLDPQYIQTHTLTEKSDVYSFGVVLVELITSQAAI   659
RR_7Ba              DFGASKLVPEDENTQLATLVQGTLGYLDPQYIQTHTLTEKSDVYSFGVVLVELITSQAAI   659
RR_7Bb              DFGASKLVPEDENTQLATLVQGTLGYLDPQYIQTHTLTEKSDVYSFGVVLVELITSQAAI   659

Brilliance          -SNKKLEAEKNLANCFLKSVADNNLDQILDHEIIKEESFEIAEQVAHLAKRCLSLKGEDR   719
FL 16.33-8-Hap1     NSNKKLEAEKNLANCFLKSVEDNHLDQILDHEIIK-DSLEIAEQVAHLAKRCLSLKGDDR   718
RR_7Ba              NSNKKLEAEKNLANCFLKSVEDNHLDQILDHEIIK-DSLEIAEQVAHLAKRCLSLKGDDR   718
RR_7Bb              NSNKKLEAEKNLANCFLKSVEDNHLDQILDHEIIK-DSLEIAEQVAHLAKRCLSLKGDDR   718

Brilliance          PTMEEVETELKVILAVMAKHPGGKPDSSPKETDYLLAASPSNAYVVDVRSDEGEVITSID   779
FL 16.33-8-Hap1     PAMREVETELVAILAVMEKRPGGKPDSSPKETDYLLAASPSNAFVVDVRSDEGELTTSID   778
RR_7Ba              PAMREVETELVAILAVMEKRPGGKPDSSPKETDYLLAASPSNAFVVDVRSDEGELTTSID   778
RR_7Bb              PAMREVETELVAILAVMEKRPGGKPDSSPKETDYLLAASPSNAFVVDVRSDEGELTTSID   778

Brilliance          YDKSMQNQAQMMKHYDGGR*                                           799
FL 16.33-8-Hap1     YDQSMQNQSQMTRPYDSGR*                                           798
RR_7Ba              YDQSMQNQSQMTRPYDSGR*                                           798
RR_7Bb              YDQSMQNQSQMTRPYDSGR*                                           798
```

**Supplementary Fig. S8. Genomic, CDS, and peptide sequence alignment of *CNGC* genes derived from FL 16.33-8-phase1, and 'Royal Royce' phase1 and 2 assemblies.**

#### Genomic sequence of *CNGC*

```
FL16.33-8-CNGC1 ATGGCTAAACATCAGGGACCAAACAATGAAATCCAAATGAGGTTTGTTTTATTAACCTAG    60
RR_7Ba-CNGC1    ATGGCTAAACATCAGGGACCAAACAATGAAATCCAAATGAGGTTTGTTTTATTAACCTAG    60
RR_7Bb-CNGC1    ATGGCTAAACATCAGGGACCAAACAATGAAATCCAAATGAGGTTTGTTTTATTAACCTAG    60
FL16.33-8-CNGC2 ATGGCTAAACATCAGGGACCAACCAATGAAATCCAAATGAGGTTTGTTTTATTAACCTAG    60
RR_7Ba-CNGC2    ATGGCTAAACATCAGGGACCAACCAATGAAATCCAAATGAGGTTTGTTTTATTAACCTAG    60
RR_7Bb-CNGC2    ATGGCTAAACATCAGGGACCAACCAATGAAATCCAAATGAGGTTTGTTTTATTAACCTAG    60

FL16.33-8-CNGC1 CAATAAGTTAGTTTTCTGCAATGGTACTGTTTCTTTTCCTTTAATTTGGAAAGGATCGAC   120
RR_7Ba-CNGC1    CAATAAGTTAGTTTTCTGCAATGGTACTGTTTCTTTTCCTTTAATTTGGAAAGGATCGAC   120
RR_7Bb-CNGC1    CAATAAGTTAGTTTTCTGCAATGGTACTGTTTCTTTTCCTTTAATTTGGAAAGGATCGAC   120
FL16.33-8-CNGC2 CAATAAGTTAGTTTTCTGCAATAGTACTGTTTCTTTTCCTTTAATTTGGAAAGGATCGAC   120
RR_7Ba-CNGC2    CAATAAGTTAGTTTTCTGCAATAGTACTGTTTCTTTTCCTTTAATTTGGAAAGGATCGAC   120
RR_7Bb-CNGC2    CAATAAGTTAGTTTTCTGCAATAGTACTGTTTCTTTTCCTTTAATTTGGAAAGGATCGAC   120

FL16.33-8-CNGC1 TTCTCCGTTCGAGAGGGTATCGGCTCTAGATCTGATAAATTTGACGGCTTCGATTCATTT   180
RR_7Ba-CNGC1    TTCTCCGTTCGAGAGGGTATCGGCTCTAGATCTGATAAATTTGACGGCTTCGATTCATTT   180
RR_7Bb-CNGC1    TTCTCCGTTCGAGAGGGTATCGGCTCTAGATCTGATAAATTTGACGGCTTCGATTCATTT   180
FL16.33-8-CNGC2 TTCTCCGTTCGAGAGGGTATCGGCTCTAGATCTGATAAATTTGACGGCTTCGATTCATTT   180
RR_7Ba-CNGC2    TTCTCCGTTCGAGAGGGTATCGGCTCTAGATCTGATAAATTTGACGGCTTCGATTCATTT   180
RR_7Bb-CNGC2    TTCTCCGTTCGAGAGGGTATCGGCTCTAGATCTGATAAATTTGACGGCTTCGATTCATTT   180

FL16.33-8-CNGC1 AATTTACATTTTTTTTGTTTTTCTTCTATCAGCATTACACCAGATGTGTCTTATGATTTC   240
RR_7Ba-CNGC1    AATTTACATTTTTTTTGTTTTTCTTCTATCAGCATTACACCAGATGTGTCTTATGATTTC   240
RR_7Bb-CNGC1    AATTTACATTTTTTTTGTTTTTCTTCTATCAGCATTACACCAGATGTGTCTTATGATTTC   240
FL16.33-8-CNGC2 AATTTACATTTTTTTTGTTTTTCTTCTATCAGCATTACACCAGATGTGTCTTATGATTTC   240
RR_7Ba-CNGC2    AATTTACATTTTTTTTGTTTTTCTTCTATCAGCATTACACCAGATGTGTCTTATGATTTC   240
RR_7Bb-CNGC2    AATTTACATTTTTTTTGTTTTTCTTCTATCAGCATTACACCAGATGTGTCTTATGATTTC   240

FL16.33-8-CNGC1 CATAGTCCTGAGGCTCTAAAACCGACACGCGAACATGGACAACCTCAAGCAAGCAGCAAA   300
RR_7Ba-CNGC1    CATAGTCCTGAGGCTCTAAAACCGACACGCGAACATGGACAACCTCAAGCAAGCAGCAAA   300
RR_7Bb-CNGC1    CATAGTCCTGAGGCTCTAAAACCGACACGCGAACATGGACAACCTCAAGCAAGCAGCAAA   300
FL16.33-8-CNGC2 CATAGTCCTGAGGCTCTAAAACCGACACCCGAACATGGACAACCTCAAGCAAGCAGCAAA   300
RR_7Ba-CNGC2    CATAGTCCTGAGGCTCTAAAACCGACACCCGAACATGGACAACCTCAAGCAAGCAGCAAA   300
RR_7Bb-CNGC2    CATAGTCCTGAGGCTCTAAAACCGACACCCGAACATGGACAACCTCAAGCAAGCAGCAAA   300

FL16.33-8-CNGC1 AGATGGAATTGGCCAGCAGTTGCCGGAGTTTCTTCAGCTTTGAACCCATGGTGGAATAAG   360
RR_7Ba-CNGC1    AGATGGAATTGGCCAGCAGTTGCCGGAGTTTCTTCAGCTTTGAACCCATGGTGGAATAAG   360
RR_7Bb-CNGC1    AGATGGAATTGGCCAGCAGTTGCCGGAGTTTCTTCAGCTTTGAACCCATGGTGGAATAAG   360
FL16.33-8-CNGC2 AGACGGAATTGGCCAACAGTTGCCGGAGTTTCTTCAGCTTTGAACCCATGGTGGAATAAG   360
RR_7Ba-CNGC2    AGACGGAATTGGCCAACAGTTGCCGGAGTTTCTTCAGCTTTGAACCCATGGTGGAATAAG   360
RR_7Bb-CNGC2    AGACGGAATTGGCCAACAGTTGCCGGAGTTTCTTCAGCTTTGAACCCATGGTGGAATAAG   360

FL16.33-8-CNGC1 ATACTGATAACTTCATGTGTGATTGCAGTTTTTTTTGATCCCTTGTTCTTTTACATTCCA   420
RR_7Ba-CNGC1    ATACTGATAACTTCATGTGTGATTGCAGTTTTTTTTGATCCCTTGTTCTTTTACATTCCA   420
RR_7Bb-CNGC1    ATACTGATAACTTCATGTGTGATTGCAGTTTTTTTTGATCCCTTGTTCTTTTACATTCCA   420
FL16.33-8-CNGC2 ATACTGGTAACTTCATGTGTGATTGCAGTTTTATTTGATCCCTTGTTCTTTTACATTCCA   420
RR_7Ba-CNGC2    ATACTGGTAACTTCATGTGTGATTGCAGTTTTATTTGATCCCTTGTTCTTTTACATTCCA   420
RR_7Bb-CNGC2    ATACTGGTAACTTCATGTGTGATTGCAGTTTTATTTGATCCCTTGTTCTTTTACATTCCA   420

FL16.33-8-CNGC1 TACACCAGCGAGGAAAAAAAGTGTATGGGAAACGACGAAAGAGTGCAGACTCCAGCTCTG   480
RR_7Ba-CNGC1    TACACCAGCGAGGAAAAAAAGTGTATGGGAAACGACGAAAGAGTGCAGACTCCAGCTCTG   480
RR_7Bb-CNGC1    TACACCAGCGAGGAAAAAAAGTGTATGGGAAACGACGAAAGAGTGCAGACTCCAGCTCTG   480
FL16.33-8-CNGC2 TACACCAGCGAGGAAAAAAAGTGTATGGGAAACGACGAAAGAGTGCAGACTCCAGCTCTG   480
RR_7Ba-CNGC2    TACACCAGCGAGGAAAAAAAGTGTATGGGAAACGACGAAAGAGTGCAGACTCCAGCTCTG   480
RR_7Bb-CNGC2    TACACCAGCGAGGAAAAAAAGTGTATGGGAAACGACGAAAGAGTGCAGACTCCAGCTCTG   480

FL16.33-8-CNGC1 ATTTTCCGATCAGTCACAGACATCATTTTCCTGGTGCATATCATATACCAACTATATGTG   540
RR_7Ba-CNGC1    ATTTTCCGATCAGTCACAGACATCATTTTCCTGGTGCATATCATATACCAACTATATGTG   540
RR_7Bb-CNGC1    ATTTTCCGATCAGTCACAGACATCATTTTCCTGGTGCATATCATATACCAACTATATGTG   540
FL16.33-8-CNGC2 ATTTTCCGATCAGTCACAGACATCATTTTCCTGGTGCATATCATATACCAACTATATGTG   540
RR_7Ba-CNGC2    ATTTTCCGATCAGTCACAGACATCATTTTCCTGGTGCATATCATATACCAACTATATGTG   540
RR_7Bb-CNGC2    ATTTTCCGATCAGTCACAGACATCATTTTCCTGGTGCATATCATATACCAACTATATGTG   540

FL16.33-8-CNGC1 GCTATCAAATTTGCACACTCAAGGGTTTCCGAGGATGAAAGTTTGGTTTTTAGATATGAA   600
RR_7Ba-CNGC1    GCTATCAAATTTGCACACTCAAGGGTTTCCGAGGATGAAAGTTTGGTTTTTAGATATGAA   600
RR_7Bb-CNGC1    GCTATCAAATTTGCACACTCAAGGGTTTCCGAGGATGAAAGTTTGGTTTTTAGATATGAA   600
FL16.33-8-CNGC2 GCTATCAAATTTGCACTCTCAAGGGTTTCCGAGGATGAAAGTTTGGTTTTTAGATATGAA   600
RR_7Ba-CNGC2    GCTATCAAATTTGCACTCTCAAGGGTTTCCGAGGATGAAAGTTTGGTTTTTAGATATGAA   600
RR_7Bb-CNGC2    GCTATCAAATTTGCACTCTCAAGGGTTTCCGAGGATGAAAGTTTGGTTTTTAGATATGAA   600

FL16.33-8-CNGC1 TCGGTATCCCGAGAGCAAATGATTCAGTTTGCGAAGGCATTTTCTCACAAGTTGTCATGG   660
RR_7Ba-CNGC1    TCGGTATCCCGAGAGCAAATGATTCAGTTTGCGAAGGCATTTTCTCACAAGTTGTCATGG   660
RR_7Bb-CNGC1    TCGGTATCCCGAGAGCAAATGATTCAGTTTGCGAAGGCATTTTCTCACAAGTTGTCATGG   660
FL16.33-8-CNGC2 TCGGTATCCCGAGAGCAAATGATTCAGTTTGCGAAGGCATTTTCTCACAAGTTGTCATGG   660
```

```
RR_7Ba-CNGC2      TCGGTATCCCGAGAGCAAATGATTCAGTTTGCGAAGGCATTTTCTCACAAGTTGTCATGG    660
RR_7Bb-CNGC2      TCGGTATCCCGAGAGCAAATGATTCAGTTTGCGAAGGCATTTTCTCACAAGTTGTCATGG    660

FL16.33-8-CNGC1   CGATCTTTCCTAACTGACGTTTTCGCTATTTTGCCCATACCACAAGTAAGACATATCGTA    720
RR_7Ba-CNGC1      CGATCTTTCCTAACTGACGTTTTCGCTATTTTGCCCATACCACAAGTAAGACATATCGTA    720
RR_7Bb-CNGC1      CGATCTTTCCTAACTGACGTTTTCGCTATTTTGCCCATACCACAAGTAAGACATATCGTA    720
FL16.33-8-CNGC2   CGATCTTTCCTAACTGACGTTTTCGCTATTTTGCCCATACCACAAGTAAGACATATCGTA    720
RR_7Ba-CNGC2      CGATCTTTCCTAACTGACGTTTTCGCTATTTTGCCCATACCACAAGTAAGACATATCGTA    720
RR_7Bb-CNGC2      CGATCTTTCCTAACTGACGTTTTCGCTATTTTGCCCATACCACAAGTAAGACATATCGTA    720

FL16.33-8-CNGC1   ACTCTCTTTCTAGTTATTAGCTAGAGATTGGTTAATTAACTGGCTGTAATTTATGGTCTT    780
RR_7Ba-CNGC1      ACTCTCTTTCTAGTTATTAGCTAGAGATTGGTTAATTAACTGGCTGTAATTTATGGTCTT    780
RR_7Bb-CNGC1      ACTCTCTTTCTAGTTATTAGCTAGAGATTGGTTAATTAACTGGCTGTAATTTATGGTCTT    780
FL16.33-8-CNGC2   ACTCTCTTTCTAGTTATTAGCTAGAGATTGGTTAATTAACTGGCTGTAATTTATGGTCTT    780
RR_7Ba-CNGC2      ACTCTCTTTCTAGTTATTAGCTAGAGATTGGTTAATTAACTGGCTGTAATTTATGGTCTT    780
RR_7Bb-CNGC2      ACTCTCTTTCTAGTTATTAGCTAGAGATTGGTTAATTAACTGGCTGTAATTTATGGTCTT    780

FL16.33-8-CNGC1   TTTAAAGTTTTAACCAACGATTTTTTGTCCAGCTTCTAATAGTAGGATTGTTTTTCAACA    840
RR_7Ba-CNGC1      TTTAAAGTTTTAACCAACGATTTTTTGTCCAGCTTCTAATAGTAGGATTGTTTTTCAACA    840
RR_7Bb-CNGC1      TTTAAAGTTTTAACCAACGATTTTTTGTCCAGCTTCTAATAGTAGGATTGTTTTTCAACA    840
FL16.33-8-CNGC2   TTTAAAGTTTTAACCAACGATTTTTTGTCCAGCTGCTAATAGTAAGATTGTTTTTCAACA    840
RR_7Ba-CNGC2      TTTAAAGTTTTAACCAACGATTTTTTGTCCAGCTGCTAATAGTAAGATTGTTTTTCAACA    840
RR_7Bb-CNGC2      TTTAAAGTTTTAACCAACGATTTTTTGTCCAGCTGCTAATAGTAAGATTGTTTTTCAACA    840

FL16.33-8-CNGC1   TCGGAGGCAATAACTACTTGTTAAAAAGAAAGATCGTGGGCGGTTTTCTTCTAATCCAAT    900
RR_7Ba-CNGC1      TCGGAGGCAATAACTACTTGTTAAAAAGAAAGATCGTGGGCGGTTTTCTTCTAATCCAAT    900
RR_7Bb-CNGC1      TCGGAGGCAATAACTACTTGTTAAAAAGAAAGATCGTGGGCGGTTTTCTTCTAATCCAAT    900
FL16.33-8-CNGC2   TTGGAGGCAATAACTACTTGTTAAAAAGAAAGATTGTGAGCGGTTTTCTTCTAATCCAAT    900
RR_7Ba-CNGC2      TTGGAGGCAATAACTACTTGTTAAAAAGAAAGATTGTGAGCGGTTTTCTTCTAATCCAAT    900
RR_7Bb-CNGC2      TTGGAGGCAATAACTACTTGTTAAAAAGAAAGATTGTGAGCGGTTTTCTTCTAATCCAAT    900

FL16.33-8-CNGC1   ATGTGCCAAGAATCTATCGAATTTATCTTTCATCTGTCGCGGTCACAGAATCTGCATTGT    960
RR_7Ba-CNGC1      ATGTGCCAAGAATCTATCGAATTTATCTTTCATCTGTCGCGGTCACAGAATCTGCATTGT    960
RR_7Bb-CNGC1      ATGTGCCAAGAATCTATCGAATTTATCTTTCATCTGTCGCGGTCACAGAATCTGCATTGT    960
FL16.33-8-CNGC2   ATGTGCCAAGAATCTATCGAATTTATCTTTCATCTGTCGCGGTCACAGAATCTGCAATGT    960
RR_7Ba-CNGC2      ATGTGCCAAGAATCTATCGAATTTATCTTTCATCTGTCGCGGTCACAGAATCTGCAATGT    960
RR_7Bb-CNGC2      ATGTGCCAAGAATCTATCGAATTTATCTTTCATCTGTCGCGGTCACAGAATCTGCAATGT    960

FL16.33-8-CNGC1   GGGTTAGAGGTGTATTTAACTTCTTCCTCTACATTCTTGCGAGTCATGTAAGTTTATCCA   1020
RR_7Ba-CNGC1      GGGTTAGAGGTGTATTTAACTTCTTCCTCTACATTCTTGCGAGTCATGTAAGTTTATCCA   1020
RR_7Bb-CNGC1      GGGTTAGAGGTGTATTTAACTTCTTCCTCTACATTCTTGCGAGTCATGTAAGTTTATCCA   1020
FL16.33-8-CNGC2   GGGTTAGAGGTGTATTTAACATTTTTCTCTACATTCTTGCGAGTCATGTAAGTTTATCCA   1020
RR_7Ba-CNGC2      GGGTTAGAGGTGTATTTAACATTTTTCTCTACATTCTTGCGAGTCATGTAAGTTTATCCA   1020
RR_7Bb-CNGC2      GGGTTAGAGGTGTATTTAACATTTTTCTCTACATTCTTGCGAGTCATGTAAGTTTATCCA   1020

FL16.33-8-CNGC1   TCTCACCTCTTCTCATCAATTCATTATGTTTTTTTTTTCATTGAATTAGGTGAGTTGCTT   1080
RR_7Ba-CNGC1      TCTCACCTCTTCTCATCAATTCATTATGTTTTTTTTTTCATTGAATTAGGTGAGTTGCTT   1080
RR_7Bb-CNGC1      TCTCACCTCTTCTCATCAATTCATTATGTTTTTTTTTTCATTGAATTAGGTGAGTTGCTT   1080
FL16.33-8-CNGC2   TCTCACCACTTCTCATCAATTCATTATG--TTTTTTTTCATTGAATTAGGTGAGTTGCTT   1078
RR_7Ba-CNGC2      TCTCACCACTTCTCATCAATTCATTATG--TTTTTTTTCATTGAATTAGGTGAGTTGCTT   1078
RR_7Bb-CNGC2      TCTCACCACTTCTCATCAATTCATTATG--TTTTTTTTCATTGAATTAGGTGAGTTGCTT   1078

FL16.33-8-CNGC1   GAGTTAAGTTTGTGTTGATCAAAAATGTTTGTTGTTAATTATAGGCACTTGGAGCCTTTT   1140
RR_7Ba-CNGC1      GAGTTAAGTTTGTGTTGATCAAAAATGTTTGTTGTTAATTATAGGCACTTGGAGCCTTTT   1140
RR_7Bb-CNGC1      GAGTTAAGTTTGTGTTGATCAAAAATGTTTGTTGTTAATTATAGGCACTTGGAGCCTTTT   1140
FL16.33-8-CNGC2   GAGTTAAGTTTGTGTTGATCAAAAATGTTTGTTGTTAATTATAGGCACTTGGAGCCTTTT   1138
RR_7Ba-CNGC2      GAGTTAAGTTTGTGTTGATCAAAAATGTTTGTTGTTAATTATAGGCACTTGGAGCCTTTT   1138
RR_7Bb-CNGC2      GAGTTAAGTTTGTGTTGATCAAAAATGTTTGTTGTTAATTATAGGCACTTGGAGCCTTTT   1138

FL16.33-8-CNGC1   GGTACTTTTGTTCTATTCAGCGAGAGACATCATGTTGGTATCGAACTTGCGTCGCTACAA   1200
RR_7Ba-CNGC1      GGTACTTTTGTTCTATTCAGCGAGAGACATCATGTTGGTATCGAACTTGCGTCGCTACAA   1200
RR_7Bb-CNGC1      GGTACTTTTGTTCTATTCAGCGAGAGACATCATGTTGGTATCGAACTTGCGTCGCTACAA   1200
FL16.33-8-CNGC2   GGTACTTTTGTTCTATTCAGCGAGAGACATTATGTTGGTATCGAACTTGCCTCGCTACAA   1198
RR_7Ba-CNGC2      GGTACTTTTGTTCTATTCAGCGAGAGACATTATGTTGGTATCGAACTTGCCTCGCTACAA   1198
RR_7Bb-CNGC2      GGTACTTTTGTTCTATTCAGCGAGAGACATTATGTTGGTATCGAACTTGCCTCGCTACAA   1198

FL16.33-8-CNGC1   ATATATCAGAATCAAAATGCACCTTTTACTGCCATAACAGTACTGCTGCACTTATAACCC   1260
RR_7Ba-CNGC1      ATATATCAGAATCAAAATGCACCTTTTACTGCCATAACAGTACTGCTGCACTTATAACCC   1260
RR_7Bb-CNGC1      ATATATCAGAATCAAAATGCACCTTTTACTGCCATAACAGTACTGCTGCACTTATAACCC   1260
FL16.33-8-CNGC2   ATATATCAGAATCAAAATGCACCTTTTACTGCCATAACAGTGCTGCTGCACTTATGACCC   1258
RR_7Ba-CNGC2      ATATATCAGAATCAAAATGCACCTTTTACTGCCATAACAGTGCTGCTGCACTTATGACCC   1258
RR_7Bb-CNGC2      ATATATCAGAATCAAAATGCACCTTTTACTGCCATAACAGTGCTGCTGCACTTATGACCC   1258

FL16.33-8-CNGC1   CACAATTCAAGACAACTCTAGACGAGAAATGCATTCTAAAAGTTCCATATAACATGACCG   1320
RR_7Ba-CNGC1      CACAATTCAAGACAACTCTAGACGAGAAATGCATTCTAAAAGTTCCATATAACATGACCG   1320
RR_7Bb-CNGC1      CACAATTCAAGACAACTCTAGACGAGAAATGCATTCTAAAAGTTCCATATAACATGACCG   1320
FL16.33-8-CNGC2   CACAATTCAAGACAACTCTAGACGAGAAATGCATTCTAAAAGTTCCATATAACATGACCG   1318
RR_7Ba-CNGC2      CACAATTCAAGACAACTCTAGACGAGAAATGCATTCTAAAAGTTCCATATAACATGACCG   1318
RR_7Bb-CNGC2      CACAATTCAAGACAACTCTAGACGAGAAATGCATTCTAAAAGTTCCATATAACATGACCG   1318

FL16.33-8-CNGC1   ATCCACCATTTGATTTTGGAATATTTTTTGATGCCCTCAAAAATGATATCCAAGGGAAAA   1380
RR_7Ba-CNGC1      ATCCACCATTTGATTTTGGAATATTTTTTGATGCCCTCAAAAATGATATCCAAGGGAAAA   1380
```

```
RR_7Bb-CNGC1     ATCCACCATTTGATTTTGGAATATTTTTTGATGCCCTCAAAAATGATATCCAAGGGAAAA  1380
FL16.33-8-CNGC2  ATCCACCATATGATTTTGGAATATTTTTTGATGCCCTCAAAAATGATATCCAAGGGAAAA  1378
RR_7Ba-CNGC2     ATCCACCATATGATTTTGGAATATTTTTTGATGCCCTCAAAAATGATATCCAAGGGAAAA  1378
RR_7Bb-CNGC2     ATCCACCATATGATTTTGGAATATTTTTTGATGCCCTCAAAAATGATATCCAAGGGAAAA  1378

FL16.33-8-CNGC1  TAAATGTTCCACAAAAGATTGGTTACTGTTTTTGGTGGGGCTTGCGAAACTTGAGGTAAA  1440
RR_7Ba-CNGC1     TAAATGTTCCACAAAAGATTGGTTACTGTTTTTGGTGGGGCTTGCGAAACTTGAGGTAAA  1440
RR_7Bb-CNGC1     TAAATGTTCCACAAAAGATTGGTTACTGTTTTTGGTGGGGCTTGCGAAACTTGAGGTAAA  1440
FL16.33-8-CNGC2  TAAATGTTCCACAAAAGATTGGTTACTGTTTTTGGTGGGGCTTGCGAAACTTGAGGTAAA  1438
RR_7Ba-CNGC2     TAAATGTTCCACAAAAGATTGGTTACTGTTTTTGGTGGGGCTTGCGAAACTTGAGGTAAA  1438
RR_7Bb-CNGC2     TAAATGTTCCACAAAAGATTGGTTACTGTTTTTGGTGGGGCTTGCGAAACTTGAGGTAAA  1438

FL16.33-8-CNGC1  TCTTCATTTATAACTTAATTCAGTCGGGTCATTCATCAACCAGCATTTTACAATTACTAA  1500
RR_7Ba-CNGC1     TCTTCATTTATAACTTAATTCAGTCGGGTCATTCATCAACCAGCATTTTACAATTACTAA  1500
RR_7Bb-CNGC1     TCTTCATTTATAACTTAATTCAGTCGGGTCATTCATCAACCAGCATTTTACAATTACTAA  1500
FL16.33-8-CNGC2  TCTTCATTTATAACTTAATTCAGTCGGGTCATTCATCAACCAGCATTTTACAATTACTAA  1498
RR_7Ba-CNGC2     TCTTCATTTATAACTTAATTCAGTCGGGTCATTCATCAACCAGCATTTTACAATTACTAA  1498
RR_7Bb-CNGC2     TCTTCATTTATAACTTAATTCAGTCGGGTCATTCATCAACCAGCATTTTACAATTACTAA  1498

FL16.33-8-CNGC1  TCATTTAACAATTTCTTCTGTGTCCGGCAGTAATTTTGGTACAAGTTTGGAAACAAGTAC  1560
RR_7Ba-CNGC1     TCATTTAACAATTTCTTCTGTGTCCGGCAGTAATTTTGGTACAAGTTTGGAAACAAGTAC  1560
RR_7Bb-CNGC1     TCATTTAACAATTTCTTCTGTGTCCGGCAGTAATTTTGGTACAAGTTTGGAAACAAGTAC  1560
FL16.33-8-CNGC2  TCATTTAACAATTTCTTCTGTGTCCGGCAGTAATTTTGGTACAAATTTGGAAACAAGTAC  1558
RR_7Ba-CNGC2     TCATTTAACAATTTCTTCTGTGTCCGGCAGTAATTTTGGTACAAATTTGGAAACAAGTAC  1558
RR_7Bb-CNGC2     TCATTTAACAATTTCTTCTGTGTCCGGCAGTAATTTTGGTACAAATTTGGAAACAAGTAC  1558

FL16.33-8-CNGC1  CTATCTGTGGGAAAACAGCTTTGCCATTTTAATTTCTATCATCGGCTTGCTGTTGTTTTT  1620
RR_7Ba-CNGC1     CTATCTGTGGGAAAACAGCTTTGCCATTTTAATTTCTATCATCGGCTTGCTGTTGTTTTT  1620
RR_7Bb-CNGC1     CTATCTGTGGGAAAACAGCTTTGCCATTTTAATTTCTATCATCGGCTTGCTGTTGTTTTT  1620
FL16.33-8-CNGC2  CTATCTGTGGGAAAACAGCTTTGCCATTTTAATTTCTATCATCGGCTTGCTGTTGTTTTT  1618
RR_7Ba-CNGC2     CTATCTGTGGGAAAACAGCTTTGCCATTTTAATTTCTATCATCGGCTTGCTGTTGTTTTT  1618
RR_7Bb-CNGC2     CTATCTGTGGGAAAACAGCTTTGCCATTTTAATTTCTATCATCGGCTTGCTGTTGTTTTT  1618

FL16.33-8-CNGC1  ATACCTCATTGGAAATGTACAGGTTAGTAACCTAATCGATCACCCATATATGTTATGTTC  1680
RR_7Ba-CNGC1     ATACCTCATTGGAAATGTACAGGTTAGTAACCTAATCGATCACCCATATATGTTATGTTC  1680
RR_7Bb-CNGC1     ATACCTCATTGGAAATGTACAGGTTAGTAACCTAATCGATCACCCATATATGTTATGTTC  1680
FL16.33-8-CNGC2  ATACCTCATTGGAAATGTACAGGTTGGTAACCTAATCGATCACCCATATATGTTATGTTC  1678
RR_7Ba-CNGC2     ATACCTCATTGGAAATGTACAGGTTGGTAACCTAATCGATCACCCATATATGTTATGTTC  1678
RR_7Bb-CNGC2     ATACCTCATTGGAAATGTACAGGTTGGTAACCTAATCGATCACCCATATATGTTATGTTC  1678

FL16.33-8-CNGC1  TTTTTAGGATTTGTTCATTTTAATTTTCCGTAAATTATTAACTTTGTTTTTCATGAAGAA  1740
RR_7Ba-CNGC1     TTTTTAGGATTTGTTCATTTTAATTTTCCGTAAATTATTAACTTTGTTTTTCATGAAGAA  1740
RR_7Bb-CNGC1     TTTTTAGGATTTGTTCATTTTAATTTTCCGTAAATTATTAACTTTGTTTTTCATGAAGAA  1740
FL16.33-8-CNGC2  TTTTTAGGATTTGTTCATTTTAATTTTCCACAAATTCTTAACTTTGTTTTTCATGAAGAA  1738
RR_7Ba-CNGC2     TTTTTAGGATTTGTTCATTTTAATTTTCCACAAATTCTTAACTTTGTTTTTCATGAAGAA  1738
RR_7Bb-CNGC2     TTTTTAGGATTTGTTCATTTTAATTTTCCACAAATTCTTAACTTTGTTTTTCATGAAGAA  1738

FL16.33-8-CNGC1  AAGGTTTAATTTTAGATTTCTTTCAAAAGAAAAAAGTTCAATTTTAGATTATAAATATGT  1800
RR_7Ba-CNGC1     AAGGTTTAATTTTAGATTTCTTTCAAAAGAAAAAAGTTCAATTTTAGATTATAAATATGT  1800
RR_7Bb-CNGC1     AAGGTTTAATTTTAGATTTCTTTCAAAAGAAAAAAGTTCAATTTTAGATTATAAATATGT  1800
FL16.33-8-CNGC2  AAGATTTAATTTTAGATTTCTTTCAAAAGAAAAAAGTTTAATTTTAGATTATAAATATGT  1798
RR_7Ba-CNGC2     AAGATTTAATTTTAGATTTCTTTCAAAAGAAAAAAGTTTAATTTTAGATTATAAATATGT  1798
RR_7Bb-CNGC2     AAGATTTAATTTTAGATTTCTTTCAAAAGAAAAAAGTTTAATTTTAGATTATAAATATGT  1798

FL16.33-8-CNGC1  GATGATGCCATGTTATTTTCACATCACTTGGTGTTGGGTGTTTACATCATTGAAACAATA  1860
RR_7Ba-CNGC1     GATGATGCCATGTTATTTTCACATCACTTGGTGTTGGGTGTTTACATCATTGAAACAATA  1860
RR_7Bb-CNGC1     GATGATGCCATGTTATTTTCACATCACTTGGTGTTGGGTGTTTACATCATTGAAACAATA  1860
FL16.33-8-CNGC2  GATGATGCCATGTTATTTTCACATCAGTTGGTGTTGGGTGTTTACATCATTCAAACAATA  1858
RR_7Ba-CNGC2     GATGATGCCATGTTATTTTCACATCAGTTGGTGTTGGGTGTTTACATCATTCAAACAATA  1858
RR_7Bb-CNGC2     GATGATGCCATGTTATTTTCACATCAGTTGGTGTTGGGTGTTTACATCATTCAAACAATA  1858

FL16.33-8-CNGC1  AATCTAAACCGTTGAACATTTTAAAATTTAATGGCTCAGAAACGGTGTCAAAATTTATGT  1920
RR_7Ba-CNGC1     AATCTAAACCGTTGAACATTTTAAAATTTAATGGCTCAGAAACGGTGTCAAAATTTATGT  1920
RR_7Bb-CNGC1     AATCTAAACCGTTGAACATTTTAAAATTTAATGGCTCAGAAACGGTGTCAAAATTTATGT  1920
FL16.33-8-CNGC2  AATCTAAACCGTTGAACATTTTAAAATTTAATGGCTCAGAAACGGTGTCAAAATTTATGT  1918
RR_7Ba-CNGC2     AATCTAAACCGTTGAACATTTTAAAATTTAATGGCTCAGAAACGGTGTCAAAATTTATGT  1918
RR_7Bb-CNGC2     AATCTAAACCGTTGAACATTTTAAAATTTAATGGCTCAGAAACGGTGTCAAAATTTATGT  1918

FL16.33-8-CNGC1  AAAATTTTATGTCTTTTGAACCCGACCCTATATATATATATATATATATATATATACACA  1980
RR_7Ba-CNGC1     AAAATTTTATGTCTTTTGAACCCGACCCTATATATATATATATATATATATATATACACA  1980
RR_7Bb-CNGC1     AAAATTTTATGTCTTTTGAACCCGACCCTATATATATATATATATATATATATATACACA  1980
FL16.33-8-CNGC2  AAAATTTTATGTCCTTTGAACCGGACCCTATATATATATAT----------------ACA  1962
RR_7Ba-CNGC2     AAAATTTTATGTCCTTTGAACCGGACCCTATATATATATAT----------------ACA  1962
RR_7Bb-CNGC2     AAAATTTTATGTCCTTTGAACCGGACCCTATATATATATAT----------------ACA  1962

FL16.33-8-CNGC1  CGCTATTGTGCATAGCAATGGTTAATTTAATTTGATCGAGGAGGTGATTAAATTGATGGA  2040
RR_7Ba-CNGC1     CGCTATTGTGCATAGCAATGGTTAATTTAATTTGATCGAGGAGGTGATTAAATTGATGGA  2040
RR_7Bb-CNGC1     CGCTATTGTGCATAGCAATGGTTAATTTAATTTGATCGAGGAGGTGATTAAATTGATGGA  2040
FL16.33-8-CNGC2  CGATATTGTGCATAGCAATGGTTAATTTAATTTGATCGAGGAGGTGATTAAATTGATGGA  2022
RR_7Ba-CNGC2     CGATATTGTGCATAGCAATGGTTAATTTAATTTGATCGAGGAGGTGATTAAATTGATGGA  2022
RR_7Bb-CNGC2     CGATATTGTGCATAGCAATGGTTAATTTAATTTGATCGAGGAGGTGATTAAATTGATGGA  2022
```

```
FL16.33-8-CNGC1 TGGAGTT-----------------------------------------------------  2047
RR_7Ba-CNGC1    TGGAGTT-----------------------------------------------------  2047
RR_7Bb-CNGC1    TGGAGTT-----------------------------------------------------  2047
FL16.33-8-CNGC2 TGGAGTTATTAATTACAGGGGCGGAGCCACACCAAGACCAAGGGGTGCCATGGCACCCCC  2082
RR_7Ba-CNGC2    TGGAGTTATTAATTACAGGGGCGGAGCCACACCAAGACCAAGGGGTGCCATGGCACCCCC  2082
RR_7Bb-CNGC2    TGGAGTTATTAATTACAGGGGCGGAGCCACACCAAGACCAAGGGGTGCCATGGCACCCCC  2082

FL16.33-8-CNGC1 ------------------------------------------------------------  2047
RR_7Ba-CNGC1    ------------------------------------------------------------  2047
RR_7Bb-CNGC1    ------------------------------------------------------------  2047
FL16.33-8-CNGC2 ACACATTTTGATGATGGTCAATTACGCCACTGTTGTAGGATTAAATGGTTACAATTGTAT  2142
RR_7Ba-CNGC2    ACACATTTTGATGATGGTCAATTACGCCACTGTTGTAGGATTAAATGGTTACAATTGTAT  2142
RR_7Bb-CNGC2    ACACATTTTGATGATGGTCAATTACGCCACTGTTGTAGGATTAAATGGTTACAATTGTAT  2142

FL16.33-8-CNGC1 ------------------------------------------------------------  2047
RR_7Ba-CNGC1    ------------------------------------------------------------  2047
RR_7Bb-CNGC1    ------------------------------------------------------------  2047
FL16.33-8-CNGC2 AATCTGGCCCCCCTTTTTATGTAATAATATGTCACGTAATAATTACTAATTTGATATTTT  2202
RR_7Ba-CNGC2    AATCTGGCCCCCCTTTTTATGTAATAATATGTCACGTAATAATTACTAATTTGATATTTT  2202
RR_7Bb-CNGC2    AATCTGGCCCCCCTTTTTATGTAATAATATGTCACGTAATAATTACTAATTTGATATTTT  2202

FL16.33-8-CNGC1 ------------------------------------------------------------  2047
RR_7Ba-CNGC1    ------------------------------------------------------------  2047
RR_7Bb-CNGC1    ------------------------------------------------------------  2047
FL16.33-8-CNGC2 GGACTATATAATCTCCTTTTTTGTTGCATCTTGCCATATTCATCACCATATCGATTAGTT  2262
RR_7Ba-CNGC2    GGACTATATAATCTCCTTTTTTGTTGCATCTTGCCATATTCATCACCATATCGATTAGTT  2262
RR_7Bb-CNGC2    GGACTATATAATCTCCTTTTTTGTTGCATCTTGCCATATTCATCACCATATCGATTAGTT  2262

FL16.33-8-CNGC1 ------------------------------------------------------------  2047
RR_7Ba-CNGC1    ------------------------------------------------------------  2047
RR_7Bb-CNGC1    ------------------------------------------------------------  2047
FL16.33-8-CNGC2 ATTAACTAATCATTTTACATGCATGTAAACTTTAAATATATAATCATTTACATATACATA  2322
RR_7Ba-CNGC2    ATTAACTAATCATTTTACATGCATGTAAACTTTAAATATATAATCATTTACATATACATA  2322
RR_7Bb-CNGC2    ATTAACTAATCATTTTACATGCATGTAAACTTTAAATATATAATCATTTACATATACATA  2322

FL16.33-8-CNGC1 ------------------------------------------------------------  2047
RR_7Ba-CNGC1    ------------------------------------------------------------  2047
RR_7Bb-CNGC1    ------------------------------------------------------------  2047
FL16.33-8-CNGC2 AATTAAACTATATATACATCAACAAAAACAATAAATTCAATTTTAATTAACATATAATTT  2382
RR_7Ba-CNGC2    AATTAAACTATATATACATCAACAAAAACAATAAATTCAATTTTAATTAACATATAATTT  2382
RR_7Bb-CNGC2    AATTAAACTATATATACATCAACAAAAACAATAAATTCAATTTTAATTAACATATAATTT  2382

FL16.33-8-CNGC1 ------------------------------------------------------------  2047
RR_7Ba-CNGC1    ------------------------------------------------------------  2047
RR_7Bb-CNGC1    ------------------------------------------------------------  2047
FL16.33-8-CNGC2 TGACAGACAATTTAAAATTTTATATATATTTTTTCGTATTTTATATCTCGGCCCCCCTAG  2442
RR_7Ba-CNGC2    TGACAGACAATTTAAAATTTTATATATATTTTTTCGTATTTTATATCTCGGCCCCCCTAG  2442
RR_7Bb-CNGC2    TGACAGACAATTTAAAATTTTATATATATTTTTTCGTATTTTATATCTCGGCCCCCCTAG  2442

FL16.33-8-CNGC1 -------------------------ATTAATTATATGCAACGTGCAGATTTATATGCAGA  2082
RR_7Ba-CNGC1    -------------------------ATTAATTATATGCAACGTGCAGATTTATATGCAGA  2082
RR_7Bb-CNGC1    -------------------------ATTAATTATATGCAACGTGCAGATTTATATGCAGA  2082
FL16.33-8-CNGC2 ACTTATAGTTCTGGCTCCGCCCCTGATTAATTATATGCAACATGCAGATTTATATGCAGA  2502
RR_7Ba-CNGC2    ACTTATAGTTCTGGCTCCGCCCCTGATTAATTATATGCAACATGCAGATTTATATGCAGA  2502
RR_7Bb-CNGC2    ACTTATAGTTCTGGCTCCGCCCCTGATTAATTATATGCAACATGCAGATTTATATGCAGA  2502

FL16.33-8-CNGC1 GGGCAACTACCAGAGCGGAGGAGGTAAGAGAGAAGCTTCGGATCAAAAAGAACGATATAG  2142
RR_7Ba-CNGC1    GGGCAACTACCAGAGCGGAGGAGGTAAGAGAGAAGCTTCGGATCAAAAAGAACGATATAG  2142
RR_7Bb-CNGC1    GGGCAACTACCAGAGCGGAGGAGGTAAGAGAGAAGCTTCGGATCAAAAAGAACGATATAG  2142
FL16.33-8-CNGC2 GGGCAGTTACCAGAGCGGAGGAGGTAAGAGAGAAGCTTCGGATCAAAAAGAACGATATAG  2562
RR_7Ba-CNGC2    GGGCAGTTACCAGAGCGGAGGAGGTAAGAGAGAAGCTTCGGATCAAAAAGAACGATATAG  2562
RR_7Bb-CNGC2    GGGCAGTTACCAGAGCGGAGGAGGTAAGAGAGAAGCTTCGGATCAAAAAGAACGATATAG  2562

FL16.33-8-CNGC1 ATAATTGGGTCCAAAATAATGGTATCGAGAAGGACATGAAGGAAGAGATCATGAAAAACA  2202
RR_7Ba-CNGC1    ATAATTGGGTCCAAAATAATGGTATCGAGAAGGACATGAAGGAAGAGATCATGAAAAACA  2202
RR_7Bb-CNGC1    ATAATTGGGTCCAAAATAATGGTATCGAGAAGGACATGAAGGAAGAGATCATGAAAAACA  2202
FL16.33-8-CNGC2 ATAATTGGATCCAAAATAATGGTATCGAGAATGACATGAAGGAAGAGATCATGAAAAACA  2622
RR_7Ba-CNGC2    ATAATTGGATCCAAAATAATGGTATCGAGAATGACATGAAGGAAGAGATCATGAAAAACA  2622
RR_7Bb-CNGC2    ATAATTGGATCCAAAATAATGGTATCGAGAATGACATGAAGGAAGAGATCATGAAAAACA  2622

FL16.33-8-CNGC1 TCGTTGAAAAGTTGGAAAACGACATAAGTGCAGATCTGGATGATATATACTCTATTCTTC  2262
RR_7Ba-CNGC1    TCGTTGAAAAGTTGGAAAACGACATAAGTGCAGATCTGGATGATATATACTCTATTCTTC  2262
RR_7Bb-CNGC1    TCGTTGAAAAGTTGGAAAACGACATAAGTGCAGATCTGGATGATATATACTCTATTCTTC  2262
FL16.33-8-CNGC2 TCGTTGAAAAGTTGGAAAACGACATAAGTGCAGATCTGGATGATATATACTCTATTCTTC  2682
RR_7Ba-CNGC2    TCGTTGAAAAGTTGGAAAACGACATAAGTGCAGATCTGGATGATATATACTCTATTCTTC  2682
RR_7Bb-CNGC2    TCGTTGAAAAGTTGGAAAACGACATAAGTGCAGATCTGGATGATATATACTCTATTCTTC  2682

FL16.33-8-CNGC1 CCCCGTATACCAGGAAAGTTGTGAAGCGATTTGTCGGCATGAGGACGCTAAGAAATGTAT  2322
RR_7Ba-CNGC1    CCCCGTATACCAGGAAAGTTGTGAAGCGATTTGTCGGCATGAGGACGCTAAGAAATGTAT  2322
RR_7Bb-CNGC1    CCCCGTATACCAGGAAAGTTGTGAAGCGATTTGTCGGCATGAGGACGCTAAGAAATGTAT  2322
FL16.33-8-CNGC2 CCCCGTATACCAGGAAAGTTGTGAAGCGATTTGTCGGCATGAGGACGCTAAGAAATGTAT  2742
RR_7Ba-CNGC2    CCCCGTATACCAGGAAAGTTGTGAAGCGATTTGTCGGCATGAGGACGCTAAGAAATGTAT  2742
```

```
RR_7Bb-CNGC2     CCCCGTATACCAGGAAAGTTGTGAAGCGATTTGTCGGCATGAGGACGCTAAGAAATGTAT  2742

FL16.33-8-CNGC1  GTCACAGTACCCTTGTTTGTGACTTGTGTCATCAGTACGTTGTGTGTGTTGTTGAAGCTA  2382
RR_7Ba-CNGC1     GTCACAGTACCCTTGTTTGTGACTTGTGTCATCAGTACGTTGTGTGTGTTGTTGAAGCTA  2382
RR_7Bb-CNGC1     GTCACAGTACCCTTGTTTGTGACTTGTGTCATCAGTACGTTGTGTGTGTTGTTGAAGCTA  2382
FL16.33-8-CNGC2  GTCACAGTACCCTTGTTTGTGACTTGTGTCATCAGTACGTTGTGTGTGTTGTTGAAGCTA  2802
RR_7Ba-CNGC2     GTCACAGTACCCTTGTTTGTGACTTGTGTCATCAGTACGTTGTGTGTGTTGTTGAAGCTA  2802
RR_7Bb-CNGC2     GTCACAGTACCCTTGTTTGTGACTTGTGTCATCAGTACGTTGTGTGTGTTGTTGAAGCTA  2802

FL16.33-8-CNGC1  GCTTACTAATCAGTGGTCTGCTTATACATACATATGTAGGTACCAATGCTTAGCCAGGTG  2442
RR_7Ba-CNGC1     GCTTACTAATCAGTGGTCTGCTTATACATACATATGTAGGTACCAATGCTTAGCCAGGTG  2442
RR_7Bb-CNGC1     GCTTACTAATCAGTGGTCTGCTTATACATACATATGTAGGTACCAATGCTTAGCCAGGTG  2442
FL16.33-8-CNGC2  GCTTACTAATCAGTGGTCTGCTTATACATACATATGTAGGTACCAATGCTTAGCCAGGTG  2862
RR_7Ba-CNGC2     GCTTACTAATCAGTGGTCTGCTTATACATACATATGTAGGTACCAATGCTTAGCCAGGTG  2862
RR_7Bb-CNGC2     GCTTACTAATCAGTGGTCTGCTTATACATACATATGTAGGTACCAATGCTTAGCCAGGTG  2862

FL16.33-8-CNGC1  GATGGAAGAGTTTTGAAGATGATGTGCGACTATCTGAAGCCGGTCAAGTACCAGGAGAAT  2502
RR_7Ba-CNGC1     GATGGAAGAGTTTTGAAGATGATGTGCGACTATCTGAAGCCGGTCAAGTACCAGGAGAAT  2502
RR_7Bb-CNGC1     GATGGAAGAGTTTTGAAGATGATGTGCGACTATCTGAAGCCGGTCAAGTACCAGGAGAAT  2502
FL16.33-8-CNGC2  GATGGAAGAGTTTTGAAGATGATGTGCGACTATCTGAAGCCGGTCAAGTACCAGGAGAAT  2922
RR_7Ba-CNGC2     GATGGAAGAGTTTTGAAGATGATGTGCGACTATCTGAAGCCGGTCAAGTACCAGGAGAAT  2922
RR_7Bb-CNGC2     GATGGAAGAGTTTTGAAGATGATGTGCGACTATCTGAAGCCGGTCAAGTACCAGGAGAAT  2922

FL16.33-8-CNGC1  ACAATGGTTTTTGAAACGGGAGTACCATTTGATAGAATGGTCTTCATTACAGATGGGTCA  2562
RR_7Ba-CNGC1     ACAATGGTTTTTGAAACGGGAGTACCATTTGATAGAATGGTCTTCATTACAGATGGGTCA  2562
RR_7Bb-CNGC1     ACAATGGTTTTTGAAACGGGAGTACCATTTGATAGAATGGTCTTCATTACAGATGGGTCA  2562
FL16.33-8-CNGC2  ACAATGGTTTTTGAAACGGGAGTACCATTTGATAGAATGGTCTTCATTACAGATGGGTCA  2982
RR_7Ba-CNGC2     ACAATGGTTTTTGAAACGGGAGTACCATTTGATAGAATGGTCTTCATTACAGATGGGTCA  2982
RR_7Bb-CNGC2     ACAATGGTTTTTGAAACGGGAGTACCATTTGATAGAATGGTCTTCATTACAGATGGGTCA  2982

FL16.33-8-CNGC1  ATGTGGACTTACACGATTGCTACTGATCATCAGACTAGTCATTCTGGAAAAGCAACGTCA  2622
RR_7Ba-CNGC1     ATGTGGACTTACACGATTGCTACTGATCATCAGACTAGTCATTCTGGAAAAGCAACGTCA  2622
RR_7Bb-CNGC1     ATGTGGACTTACACGATTGCTACTGATCATCAGACTAGTCATTCTGGAAAAGCAACGTCA  2622
FL16.33-8-CNGC2  ATGTGGACTTACACGATTGCTACTGATCATCAGACTAGTCATTCTGGAAAAGCAACGTCA  3042
RR_7Ba-CNGC2     ATGTGGACTTACACGATTGCTACTGATCATCAGACTAGTCATTCTGGAAAAGCAACGTCA  3042
RR_7Bb-CNGC2     ATGTGGACTTACACGATTGCTACTGATCATCAGACTAGTCATTCTGGAAAAGCAACGTCA  3042

FL16.33-8-CNGC1  GGAGGACTAGTTACTTCTTCGTCCACGGAAGCTCACTTCCTCAAGAAGGGTGATGCTTAT  2682
RR_7Ba-CNGC1     GGAGGACTAGTTACTTCTTCGTCCACGGAAGCTCACTTCCTCAAGAAGGGTGATGCTTAT  2682
RR_7Bb-CNGC1     GGAGGACTAGTTACTTCTTCGTCCACGGAAGCTCACTTCCTCAAGAAGGGTGATGCTTAT  2682
FL16.33-8-CNGC2  GGAGGACTAGTTACTTCTTCGTCCACGGAAGCTCACTTCCTCAAGAAGGGTGATGCTTAT  3102
RR_7Ba-CNGC2     GGAGGACTAGTTACTTCTTCGTCCACGGAAGCTCACTTCCTCAAGAAGGGTGATGCTTAT  3102
RR_7Bb-CNGC2     GGAGGACTAGTTACTTCTTCGTCCACGGAAGCTCACTTCCTCAAGAAGGGTGATGCTTAT  3102

FL16.33-8-CNGC1  GGACACATGCTTCTGCAATTAGCATCATCATCTTCTACGGCGGTTCCTATCTCAGTTGCA  2742
RR_7Ba-CNGC1     GGACACATGCTTCTGCAATTAGCATCATCATCTTCTACGGCGGTTCCTATCTCAGTTGCA  2742
RR_7Bb-CNGC1     GGACACATGCTTCTGCAATTAGCATCATCATCTTCTACGGCGGTTCCTATCTCAGTTGCA  2742
FL16.33-8-CNGC2  GGACACATGCTTCTGCAATTAGCATCATCATCTTCTACGGCGGTTCCTATCTCAGTTGCA  3162
RR_7Ba-CNGC2     GGACACATGCTTCTGCAATTAGCATCATCATCTTCTACGGCGGTTCCTATCTCAGTTGCA  3162
RR_7Bb-CNGC2     GGACACATGCTTCTGCAATTAGCATCATCATCTTCTACGGCGGTTCCTATCTCAGTTGCA  3162

FL16.33-8-CNGC1  AATGTCAGGTGCCACACAAAGGTAGAGGCGTTTGTCCTCATGGCCAATGACTTGAGAAAC  2802
RR_7Ba-CNGC1     AATGTCAGGTGCCACACAAAGGTAGAGGCGTTTGTCCTCATGGCCAATGACTTGAGAAAC  2802
RR_7Bb-CNGC1     AATGTCAGGTGCCACACAAAGGTAGAGGCGTTTGTCCTCATGGCCAATGACTTGAGAAAC  2802
FL16.33-8-CNGC2  AATGTCAGGTGCCACACAAAGGTAGAGGCGTTTGTCCTCATGGCCAATGACTTGAGAAAC  3222
RR_7Ba-CNGC2     AATGTCAGGTGCCACACAAAGGTAGAGGCGTTTGTCCTCATGGCCAATGACTTGAGAAAC  3222
RR_7Bb-CNGC2     AATGTCAGGTGCCACACAAAGGTAGAGGCGTTTGTCCTCATGGCCAATGACTTGAGAAAC  3222

FL16.33-8-CNGC1  ATAGTAACTGCGTGCGGAGATTTGTGGCCGGCTTTTGACTACAAGGCTTCTCAAGATCAG  2862
RR_7Ba-CNGC1     ATAGTAACTGCGTGCGGAGATTTGTGGCCGGCTTTTGACTACAAGGCTTCTCAAGATCAG  2862
RR_7Bb-CNGC1     ATAGTAACTGCGTGCGGAGATTTGTGGCCGGCTTTTGACTACAAGGCTTCTCAAGATCAG  2862
FL16.33-8-CNGC2  ATAGTAACTGCGTGCGGAGATTTGTGGCCGGCTTTTGACTACAAGGCTTCTCAAGATCAG  3282
RR_7Ba-CNGC2     ATAGTAACTGCGTGCGGAGATTTGTGGCCGGCTTTTGACTACAAGGCTTCTCAAGATCAG  3282
RR_7Bb-CNGC2     ATAGTAACTGCGTGCGGAGATTTGTGGCCGGCTTTTGACTACAAGGCTTCTCAAGATCAG  3282

FL16.33-8-CNGC1  GCGGCTGATCAGATGGCACCGCCGGGGCATTTTCAGCAACAACAAGGGCCAAAGAAGCGT  2922
RR_7Ba-CNGC1     GCGGCTGATCAGATGGCACCGCCGGGGCATTTTCAGCAACAACAAGGGCCAAAGAAGCGT  2922
RR_7Bb-CNGC1     GCGGCTGATCAGATGGCACCGCCGGGGCATTTTCAGCAACAACAAGGGCCAAAGAAGCGT  2922
FL16.33-8-CNGC2  GCGGCTGATCAGATGGCACCGCCGGGGCATTTTCAGCAACAACAAGGGCCAAAGAAGCGT  3342
RR_7Ba-CNGC2     GCGGCTGATCAGATGGCACCGCCGGGGCATTTTCAGCAACAACAAGGGCCAAAGAAGCGT  3342
RR_7Bb-CNGC2     GCGGCTGATCAGATGGCACCGCCGGGGCATTTTCAGCAACAACAAGGGCCAAAGAAGCGT  3342

FL16.33-8-CNGC1  ACTGTAGCATCATCAAATATCCATCCTACCGGCTACAGTGGGTGA                 2967
RR_7Ba-CNGC1     ACTGTAGCATCATCAAATATCCATCCTACCGGCTACAGTGGGTGA                 2967
RR_7Bb-CNGC1     ACTGTAGCATCATCAAATATCCATCCTACCGGCTACAGTGGGTGA                 2967
FL16.33-8-CNGC2  ACTGTAGCATCATCAAATATCCATCCTACCGGCTACAGTGGGTGA                 3387
RR_7Ba-CNGC2     ACTGTAGCATCATCAAATATCCATCCTACCGGCTACAGTGGGTGA                 3387
RR_7Bb-CNGC2     ACTGTAGCATCATCAAATATCCATCCTACCGGCTACAGTGGGTGA                 3387
```

**CDS of CNGC**

```
FL16.33-8-CNGC1 ATGGCTAAACATCAGGGACCAAACAATGAAATCCAAATGAGCATTACACCAGATGTGTCT  60
RR_7Ba-CNGC1    ATGGCTAAACATCAGGGACCAAACAATGAAATCCAAATGAGCATTACACCAGATGTGTCT  60
RR_7Bb-CNGC1    ATGGCTAAACATCAGGGACCAAACAATGAAATCCAAATGAGCATTACACCAGATGTGTCT  60
FL16.33-8-CNGC2 ATGGCTAAACATCAGGGACCAACCAATGAAATCCAAATGAGCATTACACCAGATGTGTCT  60
RR_7Ba-CNGC2    ATGGCTAAACATCAGGGACCAACCAATGAAATCCAAATGAGCATTACACCAGATGTGTCT  60
RR_7Bb-CNGC2    ATGGCTAAACATCAGGGACCAACCAATGAAATCCAAATGAGCATTACACCAGATGTGTCT  60

FL16.33-8-CNGC1 TATGATTTCCATAGTCCTGAGGCTCTAAAACCGACACGCGAACATGGACAACCTCAAGCA  120
RR_7Ba-CNGC1    TATGATTTCCATAGTCCTGAGGCTCTAAAACCGACACGCGAACATGGACAACCTCAAGCA  120
RR_7Bb-CNGC1    TATGATTTCCATAGTCCTGAGGCTCTAAAACCGACACGCGAACATGGACAACCTCAAGCA  120
FL16.33-8-CNGC2 TATGATTTCCATAGTCCTGAGGCTCTAAAACCGACACCCGAACATGGACAACCTCAAGCA  120
RR_7Ba-CNGC2    TATGATTTCCATAGTCCTGAGGCTCTAAAACCGACACCCGAACATGGACAACCTCAAGCA  120
RR_7Bb-CNGC2    TATGATTTCCATAGTCCTGAGGCTCTAAAACCGACACCCGAACATGGACAACCTCAAGCA  120

FL16.33-8-CNGC1 AGCAGCAAAAGATGGAATTGGCCAGCAGTTGCCGGAGTTTCTTCAGCTTTGAACCCATGG  180
RR_7Ba-CNGC1    AGCAGCAAAAGATGGAATTGGCCAGCAGTTGCCGGAGTTTCTTCAGCTTTGAACCCATGG  180
RR_7Bb-CNGC1    AGCAGCAAAAGATGGAATTGGCCAGCAGTTGCCGGAGTTTCTTCAGCTTTGAACCCATGG  180
FL16.33-8-CNGC2 AGCAGCAAAAGACGGAATTGGCCAACAGTTGCCGGAGTTTCTTCAGCTTTGAACCCATGG  180
RR_7Ba-CNGC2    AGCAGCAAAAGACGGAATTGGCCAACAGTTGCCGGAGTTTCTTCAGCTTTGAACCCATGG  180
RR_7Bb-CNGC2    AGCAGCAAAAGACGGAATTGGCCAACAGTTGCCGGAGTTTCTTCAGCTTTGAACCCATGG  180

FL16.33-8-CNGC1 TGGAATAAGATACTGATAACTTCATGTGTGATTGCAGTTTTTTTTGATCCCTTGTTCTTT  240
RR_7Ba-CNGC1    TGGAATAAGATACTGATAACTTCATGTGTGATTGCAGTTTTTTTTGATCCCTTGTTCTTT  240
RR_7Bb-CNGC1    TGGAATAAGATACTGATAACTTCATGTGTGATTGCAGTTTTTTTTGATCCCTTGTTCTTT  240
FL16.33-8-CNGC2 TGGAATAAGATACTGGTAACTTCATGTGTGATTGCAGTTTTATTTGATCCCTTGTTCTTT  240
RR_7Ba-CNGC2    TGGAATAAGATACTGGTAACTTCATGTGTGATTGCAGTTTTATTTGATCCCTTGTTCTTT  240
RR_7Bb-CNGC2    TGGAATAAGATACTGGTAACTTCATGTGTGATTGCAGTTTTATTTGATCCCTTGTTCTTT  240

FL16.33-8-CNGC1 TACATTCCATACACCAGCGAGGAAAAAAAGTGTATGGGAAACGACGAAAGAGTGCAGACT  300
RR_7Ba-CNGC1    TACATTCCATACACCAGCGAGGAAAAAAAGTGTATGGGAAACGACGAAAGAGTGCAGACT  300
RR_7Bb-CNGC1    TACATTCCATACACCAGCGAGGAAAAAAAGTGTATGGGAAACGACGAAAGAGTGCAGACT  300
FL16.33-8-CNGC2 TACATTCCATACACCAGCGAGGAAAAAAAGTGTATGGGAAACGACGAAAGAGTGCAGACT  300
RR_7Ba-CNGC2    TACATTCCATACACCAGCGAGGAAAAAAAGTGTATGGGAAACGACGAAAGAGTGCAGACT  300
RR_7Bb-CNGC2    TACATTCCATACACCAGCGAGGAAAAAAAGTGTATGGGAAACGACGAAAGAGTGCAGACT  300

FL16.33-8-CNGC1 CCAGCTCTGATTTTCCGATCAGTCACAGACATCATTTTCCTGGTGCATATCATATACCAA  360
RR_7Ba-CNGC1    CCAGCTCTGATTTTCCGATCAGTCACAGACATCATTTTCCTGGTGCATATCATATACCAA  360
RR_7Bb-CNGC1    CCAGCTCTGATTTTCCGATCAGTCACAGACATCATTTTCCTGGTGCATATCATATACCAA  360
FL16.33-8-CNGC2 CCAGCTCTGATTTTCCGATCAGTCACAGACATCATTTTCCTGGTGCATATCATATACCAA  360
RR_7Ba-CNGC2    CCAGCTCTGATTTTCCGATCAGTCACAGACATCATTTTCCTGGTGCATATCATATACCAA  360
RR_7Bb-CNGC2    CCAGCTCTGATTTTCCGATCAGTCACAGACATCATTTTCCTGGTGCATATCATATACCAA  360

FL16.33-8-CNGC1 CTATATGTGGCTATCAAATTTGCACACTCAAGGGTTTCCGAGGATGAAAGTTTGGTTTTT  420
RR_7Ba-CNGC1    CTATATGTGGCTATCAAATTTGCACACTCAAGGGTTTCCGAGGATGAAAGTTTGGTTTTT  420
RR_7Bb-CNGC1    CTATATGTGGCTATCAAATTTGCACACTCAAGGGTTTCCGAGGATGAAAGTTTGGTTTTT  420
FL16.33-8-CNGC2 CTATATGTGGCTATCAAATTTGCACTCTCAAGGGTTTCCGAGGATGAAAGTTTGGTTTTT  420
RR_7Ba-CNGC2    CTATATGTGGCTATCAAATTTGCACTCTCAAGGGTTTCCGAGGATGAAAGTTTGGTTTTT  420
RR_7Bb-CNGC2    CTATATGTGGCTATCAAATTTGCACTCTCAAGGGTTTCCGAGGATGAAAGTTTGGTTTTT  420

FL16.33-8-CNGC1 AGATATGAATCGGTATCCCGAGAGCAAATGATTCAGTTTGCGAAGGCATTTTCTCACAAG  480
RR_7Ba-CNGC1    AGATATGAATCGGTATCCCGAGAGCAAATGATTCAGTTTGCGAAGGCATTTTCTCACAAG  480
RR_7Bb-CNGC1    AGATATGAATCGGTATCCCGAGAGCAAATGATTCAGTTTGCGAAGGCATTTTCTCACAAG  480
FL16.33-8-CNGC2 AGATATGAATCGGTATCCCGAGAGCAAATGATTCAGTTTGCGAAGGCATTTTCTCACAAG  480
RR_7Ba-CNGC2    AGATATGAATCGGTATCCCGAGAGCAAATGATTCAGTTTGCGAAGGCATTTTCTCACAAG  480
RR_7Bb-CNGC2    AGATATGAATCGGTATCCCGAGAGCAAATGATTCAGTTTGCGAAGGCATTTTCTCACAAG  480

FL16.33-8-CNGC1 TTGTCATGGCGATCTTTCCTAACTGACGTTTTCGCTATTTTGCCCATACCACAACTTCTA  540
RR_7Ba-CNGC1    TTGTCATGGCGATCTTTCCTAACTGACGTTTTCGCTATTTTGCCCATACCACAACTTCTA  540
RR_7Bb-CNGC1    TTGTCATGGCGATCTTTCCTAACTGACGTTTTCGCTATTTTGCCCATACCACAACTTCTA  540
FL16.33-8-CNGC2 TTGTCATGGCGATCTTTCCTAACTGACGTTTTCGCTATTTTGCCCATACCACAACTGCTA  540
RR_7Ba-CNGC2    TTGTCATGGCGATCTTTCCTAACTGACGTTTTCGCTATTTTGCCCATACCACAACTGCTA  540
RR_7Bb-CNGC2    TTGTCATGGCGATCTTTCCTAACTGACGTTTTCGCTATTTTGCCCATACCACAACTGCTA  540

FL16.33-8-CNGC1 ATAGTAGGATTGTTTTTCAACATCGGAGGCAATAACTACTTGTTAAAAAGAAAGATCGTG  600
RR_7Ba-CNGC1    ATAGTAGGATTGTTTTTCAACATCGGAGGCAATAACTACTTGTTAAAAAGAAAGATCGTG  600
RR_7Bb-CNGC1    ATAGTAGGATTGTTTTTCAACATCGGAGGCAATAACTACTTGTTAAAAAGAAAGATCGTG  600
FL16.33-8-CNGC2 ATAGTAAGATTGTTTTTCAACATTGGAGGCAATAACTACTTGTTAAAAAGAAAGATTGTG  600
RR_7Ba-CNGC2    ATAGTAAGATTGTTTTTCAACATTGGAGGCAATAACTACTTGTTAAAAAGAAAGATTGTG  600
RR_7Bb-CNGC2    ATAGTAAGATTGTTTTTCAACATTGGAGGCAATAACTACTTGTTAAAAAGAAAGATTGTG  600

FL16.33-8-CNGC1 GGCGGTTTTCTTCTAATCCAATATGTGCCAAGAATCTATCGAATTTATCTTTCATCTGTC  660
RR_7Ba-CNGC1    GGCGGTTTTCTTCTAATCCAATATGTGCCAAGAATCTATCGAATTTATCTTTCATCTGTC  660
RR_7Bb-CNGC1    GGCGGTTTTCTTCTAATCCAATATGTGCCAAGAATCTATCGAATTTATCTTTCATCTGTC  660
FL16.33-8-CNGC2 AGCGGTTTTCTTCTAATCCAATATGTGCCAAGAATCTATCGAATTTATCTTTCATCTGTC  660
RR_7Ba-CNGC2    AGCGGTTTTCTTCTAATCCAATATGTGCCAAGAATCTATCGAATTTATCTTTCATCTGTC  660
RR_7Bb-CNGC2    AGCGGTTTTCTTCTAATCCAATATGTGCCAAGAATCTATCGAATTTATCTTTCATCTGTC  660

FL16.33-8-CNGC1 GCGGTCACAGAATCTGCATTGTGGGTTAGAGGTGTATTTAACTTCTTCCTCTACATTCTT  720
RR_7Ba-CNGC1    GCGGTCACAGAATCTGCATTGTGGGTTAGAGGTGTATTTAACTTCTTCCTCTACATTCTT  720
```

```
RR_7Bb-CNGC1     GCGGTCACAGAATCTGCATTGTGGGTTAGAGGTGTATTTAACTTCTTCCTCTACATTCTT    720
FL16.33-8-CNGC2  GCGGTCACAGAATCTGCAATGTGGGTTAGAGGTGTATTTAACATTTTTCTCTACATTCTT    720
RR_7Ba-CNGC2     GCGGTCACAGAATCTGCAATGTGGGTTAGAGGTGTATTTAACATTTTTCTCTACATTCTT    720
RR_7Bb-CNGC2     GCGGTCACAGAATCTGCAATGTGGGTTAGAGGTGTATTTAACATTTTTCTCTACATTCTT    720

FL16.33-8-CNGC1  GCGAGTCATGCACTTGGAGCCTTTTGGTACTTTTGTTCTATTCAGCGAGAGACATCATGT    780
RR_7Ba-CNGC1     GCGAGTCATGCACTTGGAGCCTTTTGGTACTTTTGTTCTATTCAGCGAGAGACATCATGT    780
RR_7Bb-CNGC1     GCGAGTCATGCACTTGGAGCCTTTTGGTACTTTTGTTCTATTCAGCGAGAGACATCATGT    780
FL16.33-8-CNGC2  GCGAGTCATGCACTTGGAGCCTTTTGGTACTTTTGTTCTATTCAGCGAGAGACATTATGT    780
RR_7Ba-CNGC2     GCGAGTCATGCACTTGGAGCCTTTTGGTACTTTTGTTCTATTCAGCGAGAGACATTATGT    780
RR_7Bb-CNGC2     GCGAGTCATGCACTTGGAGCCTTTTGGTACTTTTGTTCTATTCAGCGAGAGACATTATGT    780

FL16.33-8-CNGC1  TGGTATCGAACTTGCGTCGCTACAAATATATCAGAATCAAAATGCACCTTTTACTGCCAT    840
RR_7Ba-CNGC1     TGGTATCGAACTTGCGTCGCTACAAATATATCAGAATCAAAATGCACCTTTTACTGCCAT    840
RR_7Bb-CNGC1     TGGTATCGAACTTGCGTCGCTACAAATATATCAGAATCAAAATGCACCTTTTACTGCCAT    840
FL16.33-8-CNGC2  TGGTATCGAACTTGCCTCGCTACAAATATATCAGAATCAAAATGCACCTTTTACTGCCAT    840
RR_7Ba-CNGC2     TGGTATCGAACTTGCCTCGCTACAAATATATCAGAATCAAAATGCACCTTTTACTGCCAT    840
RR_7Bb-CNGC2     TGGTATCGAACTTGCCTCGCTACAAATATATCAGAATCAAAATGCACCTTTTACTGCCAT    840

FL16.33-8-CNGC1  AACAGTACTGCTGCACTTATAACCCCACAATTCAAGACAACTCTAGACGAGAAATGCATT    900
RR_7Ba-CNGC1     AACAGTACTGCTGCACTTATAACCCCACAATTCAAGACAACTCTAGACGAGAAATGCATT    900
RR_7Bb-CNGC1     AACAGTACTGCTGCACTTATAACCCCACAATTCAAGACAACTCTAGACGAGAAATGCATT    900
FL16.33-8-CNGC2  AACAGTGCTGCTGCACTTATGACCCCACAATTCAAGACAACTCTAGACGAGAAATGCATT    900
RR_7Ba-CNGC2     AACAGTGCTGCTGCACTTATGACCCCACAATTCAAGACAACTCTAGACGAGAAATGCATT    900
RR_7Bb-CNGC2     AACAGTGCTGCTGCACTTATGACCCCACAATTCAAGACAACTCTAGACGAGAAATGCATT    900

FL16.33-8-CNGC1  CTAAAAGTTCCATATAACATGACCGATCCACCATTTGATTTTGGAATATTTTTTGATGCC    960
RR_7Ba-CNGC1     CTAAAAGTTCCATATAACATGACCGATCCACCATTTGATTTTGGAATATTTTTTGATGCC    960
RR_7Bb-CNGC1     CTAAAAGTTCCATATAACATGACCGATCCACCATTTGATTTTGGAATATTTTTTGATGCC    960
FL16.33-8-CNGC2  CTAAAAGTTCCATATAACATGACCGATCCACCATATGATTTTGGAATATTTTTTGATGCC    960
RR_7Ba-CNGC2     CTAAAAGTTCCATATAACATGACCGATCCACCATATGATTTTGGAATATTTTTTGATGCC    960
RR_7Bb-CNGC2     CTAAAAGTTCCATATAACATGACCGATCCACCATATGATTTTGGAATATTTTTTGATGCC    960

FL16.33-8-CNGC1  CTCAAAAATGATATCCAAGGGAAAATAAATGTTCCACAAAAGATTGGTTACTGTTTTTGG   1020
RR_7Ba-CNGC1     CTCAAAAATGATATCCAAGGGAAAATAAATGTTCCACAAAAGATTGGTTACTGTTTTTGG   1020
RR_7Bb-CNGC1     CTCAAAAATGATATCCAAGGGAAAATAAATGTTCCACAAAAGATTGGTTACTGTTTTTGG   1020
FL16.33-8-CNGC2  CTCAAAAATGATATCCAAGGGAAAATAAATGTTCCACAAAAGATTGGTTACTGTTTTTGG   1020
RR_7Ba-CNGC2     CTCAAAAATGATATCCAAGGGAAAATAAATGTTCCACAAAAGATTGGTTACTGTTTTTGG   1020
RR_7Bb-CNGC2     CTCAAAAATGATATCCAAGGGAAAATAAATGTTCCACAAAAGATTGGTTACTGTTTTTGG   1020

FL16.33-8-CNGC1  TGGGGCTTGCGAAACTTGAGTAATTTTGGTACAAGTTTGGAAACAAGTACCTATCTGTGG   1080
RR_7Ba-CNGC1     TGGGGCTTGCGAAACTTGAGTAATTTTGGTACAAGTTTGGAAACAAGTACCTATCTGTGG   1080
RR_7Bb-CNGC1     TGGGGCTTGCGAAACTTGAGTAATTTTGGTACAAGTTTGGAAACAAGTACCTATCTGTGG   1080
FL16.33-8-CNGC2  TGGGGCTTGCGAAACTTGAGTAATTTTGGTACAAATTTGGAAACAAGTACCTATCTGTGG   1080
RR_7Ba-CNGC2     TGGGGCTTGCGAAACTTGAGTAATTTTGGTACAAATTTGGAAACAAGTACCTATCTGTGG   1080
RR_7Bb-CNGC2     TGGGGCTTGCGAAACTTGAGTAATTTTGGTACAAATTTGGAAACAAGTACCTATCTGTGG   1080

FL16.33-8-CNGC1  GAAAACAGCTTTGCCATTTTAATTTCTATCATCGGCTTGCTGTTGTTTTTATACCTCATT   1140
RR_7Ba-CNGC1     GAAAACAGCTTTGCCATTTTAATTTCTATCATCGGCTTGCTGTTGTTTTTATACCTCATT   1140
RR_7Bb-CNGC1     GAAAACAGCTTTGCCATTTTAATTTCTATCATCGGCTTGCTGTTGTTTTTATACCTCATT   1140
FL16.33-8-CNGC2  GAAAACAGCTTTGCCATTTTAATTTCTATCATCGGCTTGCTGTTGTTTTTATACCTCATT   1140
RR_7Ba-CNGC2     GAAAACAGCTTTGCCATTTTAATTTCTATCATCGGCTTGCTGTTGTTTTTATACCTCATT   1140
RR_7Bb-CNGC2     GAAAACAGCTTTGCCATTTTAATTTCTATCATCGGCTTGCTGTTGTTTTTATACCTCATT   1140

FL16.33-8-CNGC1  GGAAATGTACAGATTTATATGCAGAGGGCAACTACCAGAGCGGAGGAGGTAAGAGAGAAG   1200
RR_7Ba-CNGC1     GGAAATGTACAGATTTATATGCAGAGGGCAACTACCAGAGCGGAGGAGGTAAGAGAGAAG   1200
RR_7Bb-CNGC1     GGAAATGTACAGATTTATATGCAGAGGGCAACTACCAGAGCGGAGGAGGTAAGAGAGAAG   1200
FL16.33-8-CNGC2  GGAAATGTACAGATTTATATGCAGAGGGCAGTTACCAGAGCGGAGGAGGTAAGAGAGAAG   1200
RR_7Ba-CNGC2     GGAAATGTACAGATTTATATGCAGAGGGCAGTTACCAGAGCGGAGGAGGTAAGAGAGAAG   1200
RR_7Bb-CNGC2     GGAAATGTACAGATTTATATGCAGAGGGCAGTTACCAGAGCGGAGGAGGTAAGAGAGAAG   1200

FL16.33-8-CNGC1  CTTCGGATCAAAAAGAACGATATAGATAATTGGGTCCAAAATAATGGTATCGAGAAGGAC   1260
RR_7Ba-CNGC1     CTTCGGATCAAAAAGAACGATATAGATAATTGGGTCCAAAATAATGGTATCGAGAAGGAC   1260
RR_7Bb-CNGC1     CTTCGGATCAAAAAGAACGATATAGATAATTGGGTCCAAAATAATGGTATCGAGAAGGAC   1260
FL16.33-8-CNGC2  CTTCGGATCAAAAAGAACGATATAGATAATTGGATCCAAAATAATGGTATCGAGAATGAC   1260
RR_7Ba-CNGC2     CTTCGGATCAAAAAGAACGATATAGATAATTGGATCCAAAATAATGGTATCGAGAATGAC   1260
RR_7Bb-CNGC2     CTTCGGATCAAAAAGAACGATATAGATAATTGGATCCAAAATAATGGTATCGAGAATGAC   1260

FL16.33-8-CNGC1  ATGAAGGAAGAGATCATGAAAAACATCGTTGAAAAGTTGGAAAACGACATAAGTGCAGAT   1320
RR_7Ba-CNGC1     ATGAAGGAAGAGATCATGAAAAACATCGTTGAAAAGTTGGAAAACGACATAAGTGCAGAT   1320
RR_7Bb-CNGC1     ATGAAGGAAGAGATCATGAAAAACATCGTTGAAAAGTTGGAAAACGACATAAGTGCAGAT   1320
FL16.33-8-CNGC2  ATGAAGGAAGAGATCATGAAAAACATCGTTGAAAAGTTGGAAAACGACATAAGTGCAGAT   1320
RR_7Ba-CNGC2     ATGAAGGAAGAGATCATGAAAAACATCGTTGAAAAGTTGGAAAACGACATAAGTGCAGAT   1320
RR_7Bb-CNGC2     ATGAAGGAAGAGATCATGAAAAACATCGTTGAAAAGTTGGAAAACGACATAAGTGCAGAT   1320

FL16.33-8-CNGC1  CTGGATGATATATACTCTATTCTTCCCCCGTATACCAGGAAAGTTGTGAAGCGATTTGTC   1380
RR_7Ba-CNGC1     CTGGATGATATATACTCTATTCTTCCCCCGTATACCAGGAAAGTTGTGAAGCGATTTGTC   1380
RR_7Bb-CNGC1     CTGGATGATATATACTCTATTCTTCCCCCGTATACCAGGAAAGTTGTGAAGCGATTTGTC   1380
FL16.33-8-CNGC2  CTGGATGATATATACTCTATTCTTCCCCCGTATACCAGGAAAGTTGTGAAGCGATTTGTC   1380
RR_7Ba-CNGC2     CTGGATGATATATACTCTATTCTTCCCCCGTATACCAGGAAAGTTGTGAAGCGATTTGTC   1380
RR_7Bb-CNGC2     CTGGATGATATATACTCTATTCTTCCCCCGTATACCAGGAAAGTTGTGAAGCGATTTGTC   1380
```

```
FL16.33-8-CNGC1 GGCATGAGGACGCTAAGAAATGTACCAATGCTTAGCCAGGTGGATGGAAGAGTTTTGAAG  1440
RR_7Ba-CNGC1    GGCATGAGGACGCTAAGAAATGTACCAATGCTTAGCCAGGTGGATGGAAGAGTTTTGAAG  1440
RR_7Bb-CNGC1    GGCATGAGGACGCTAAGAAATGTACCAATGCTTAGCCAGGTGGATGGAAGAGTTTTGAAG  1440
FL16.33-8-CNGC2 GGCATGAGGACGCTAAGAAATGTACCAATGCTTAGCCAGGTGGATGGAAGAGTTTTGAAG  1440
RR_7Ba-CNGC2    GGCATGAGGACGCTAAGAAATGTACCAATGCTTAGCCAGGTGGATGGAAGAGTTTTGAAG  1440
RR_7Bb-CNGC2    GGCATGAGGACGCTAAGAAATGTACCAATGCTTAGCCAGGTGGATGGAAGAGTTTTGAAG  1440

FL16.33-8-CNGC1 ATGATGTGCGACTATCTGAAGCCGGTCAAGTACCAGGAGAATACAATGGTTTTTGAAACG  1500
RR_7Ba-CNGC1    ATGATGTGCGACTATCTGAAGCCGGTCAAGTACCAGGAGAATACAATGGTTTTTGAAACG  1500
RR_7Bb-CNGC1    ATGATGTGCGACTATCTGAAGCCGGTCAAGTACCAGGAGAATACAATGGTTTTTGAAACG  1500
FL16.33-8-CNGC2 ATGATGTGCGACTATCTGAAGCCGGTCAAGTACCAGGAGAATACAATGGTTTTTGAAACG  1500
RR_7Ba-CNGC2    ATGATGTGCGACTATCTGAAGCCGGTCAAGTACCAGGAGAATACAATGGTTTTTGAAACG  1500
RR_7Bb-CNGC2    ATGATGTGCGACTATCTGAAGCCGGTCAAGTACCAGGAGAATACAATGGTTTTTGAAACG  1500

FL16.33-8-CNGC1 GGAGTACCATTTGATAGAATGGTCTTCATTACAGATGGGTCAATGTGGACTTACACGATT  1560
RR_7Ba-CNGC1    GGAGTACCATTTGATAGAATGGTCTTCATTACAGATGGGTCAATGTGGACTTACACGATT  1560
RR_7Bb-CNGC1    GGAGTACCATTTGATAGAATGGTCTTCATTACAGATGGGTCAATGTGGACTTACACGATT  1560
FL16.33-8-CNGC2 GGAGTACCATTTGATAGAATGGTCTTCATTACAGATGGGTCAATGTGGACTTACACGATT  1560
RR_7Ba-CNGC2    GGAGTACCATTTGATAGAATGGTCTTCATTACAGATGGGTCAATGTGGACTTACACGATT  1560
RR_7Bb-CNGC2    GGAGTACCATTTGATAGAATGGTCTTCATTACAGATGGGTCAATGTGGACTTACACGATT  1560

FL16.33-8-CNGC1 GCTACTGATCATCAGACTAGTCATTCTGGAAAAGCAACGTCAGGAGGACTAGTTACTTCT  1620
RR_7Ba-CNGC1    GCTACTGATCATCAGACTAGTCATTCTGGAAAAGCAACGTCAGGAGGACTAGTTACTTCT  1620
RR_7Bb-CNGC1    GCTACTGATCATCAGACTAGTCATTCTGGAAAAGCAACGTCAGGAGGACTAGTTACTTCT  1620
FL16.33-8-CNGC2 GCTACTGATCATCAGACTAGTCATTCTGGAAAAGCAACGTCAGGAGGACTAGTTACTTCT  1620
RR_7Ba-CNGC2    GCTACTGATCATCAGACTAGTCATTCTGGAAAAGCAACGTCAGGAGGACTAGTTACTTCT  1620
RR_7Bb-CNGC2    GCTACTGATCATCAGACTAGTCATTCTGGAAAAGCAACGTCAGGAGGACTAGTTACTTCT  1620

FL16.33-8-CNGC1 TCGTCCACGGAAGCTCACTTCCTCAAGAAGGGTGATGCTTATGGACACATGCTTCTGCAA  1680
RR_7Ba-CNGC1    TCGTCCACGGAAGCTCACTTCCTCAAGAAGGGTGATGCTTATGGACACATGCTTCTGCAA  1680
RR_7Bb-CNGC1    TCGTCCACGGAAGCTCACTTCCTCAAGAAGGGTGATGCTTATGGACACATGCTTCTGCAA  1680
FL16.33-8-CNGC2 TCGTCCACGGAAGCTCACTTCCTCAAGAAGGGTGATGCTTATGGACACATGCTTCTGCAA  1680
RR_7Ba-CNGC2    TCGTCCACGGAAGCTCACTTCCTCAAGAAGGGTGATGCTTATGGACACATGCTTCTGCAA  1680
RR_7Bb-CNGC2    TCGTCCACGGAAGCTCACTTCCTCAAGAAGGGTGATGCTTATGGACACATGCTTCTGCAA  1680

FL16.33-8-CNGC1 TTAGCATCATCATCTTCTACGGCGGTTCCTATCTCAGTTGCAAATGTCAGGTGCCACACA  1740
RR_7Ba-CNGC1    TTAGCATCATCATCTTCTACGGCGGTTCCTATCTCAGTTGCAAATGTCAGGTGCCACACA  1740
RR_7Bb-CNGC1    TTAGCATCATCATCTTCTACGGCGGTTCCTATCTCAGTTGCAAATGTCAGGTGCCACACA  1740
FL16.33-8-CNGC2 TTAGCATCATCATCTTCTACGGCGGTTCCTATCTCAGTTGCAAATGTCAGGTGCCACACA  1740
RR_7Ba-CNGC2    TTAGCATCATCATCTTCTACGGCGGTTCCTATCTCAGTTGCAAATGTCAGGTGCCACACA  1740
RR_7Bb-CNGC2    TTAGCATCATCATCTTCTACGGCGGTTCCTATCTCAGTTGCAAATGTCAGGTGCCACACA  1740

FL16.33-8-CNGC1 AAGGTAGAGGCGTTTGTCCTCATGGCCAATGACTTGAGAAACATAGTAACTGCGTGCGGA  1800
RR_7Ba-CNGC1    AAGGTAGAGGCGTTTGTCCTCATGGCCAATGACTTGAGAAACATAGTAACTGCGTGCGGA  1800
RR_7Bb-CNGC1    AAGGTAGAGGCGTTTGTCCTCATGGCCAATGACTTGAGAAACATAGTAACTGCGTGCGGA  1800
FL16.33-8-CNGC2 AAGGTAGAGGCGTTTGTCCTCATGGCCAATGACTTGAGAAACATAGTAACTGCGTGCGGA  1800
RR_7Ba-CNGC2    AAGGTAGAGGCGTTTGTCCTCATGGCCAATGACTTGAGAAACATAGTAACTGCGTGCGGA  1800
RR_7Bb-CNGC2    AAGGTAGAGGCGTTTGTCCTCATGGCCAATGACTTGAGAAACATAGTAACTGCGTGCGGA  1800

FL16.33-8-CNGC1 GATTTGTGGCCGGCTTTTGACTACAAGGCTTCTCAAGATCAGGCGGCTGATCAGATGGCA  1860
RR_7Ba-CNGC1    GATTTGTGGCCGGCTTTTGACTACAAGGCTTCTCAAGATCAGGCGGCTGATCAGATGGCA  1860
RR_7Bb-CNGC1    GATTTGTGGCCGGCTTTTGACTACAAGGCTTCTCAAGATCAGGCGGCTGATCAGATGGCA  1860
FL16.33-8-CNGC2 GATTTGTGGCCGGCTTTTGACTACAAGGCTTCTCAAGATCAGGCGGCTGATCAGATGGCA  1860
RR_7Ba-CNGC2    GATTTGTGGCCGGCTTTTGACTACAAGGCTTCTCAAGATCAGGCGGCTGATCAGATGGCA  1860
RR_7Bb-CNGC2    GATTTGTGGCCGGCTTTTGACTACAAGGCTTCTCAAGATCAGGCGGCTGATCAGATGGCA  1860

FL16.33-8-CNGC1 CCGCCGGGGCATTTTCAGCAACAACAAGGGCCAAAGAAGCGTACTGTAGCATCATCAAAT  1920
RR_7Ba-CNGC1    CCGCCGGGGCATTTTCAGCAACAACAAGGGCCAAAGAAGCGTACTGTAGCATCATCAAAT  1920
RR_7Bb-CNGC1    CCGCCGGGGCATTTTCAGCAACAACAAGGGCCAAAGAAGCGTACTGTAGCATCATCAAAT  1920
FL16.33-8-CNGC2 CCGCCGGGGCATTTTCAGCAACAACAAGGGCCAAAGAAGCGTACTGTAGCATCATCAAAT  1920
RR_7Ba-CNGC2    CCGCCGGGGCATTTTCAGCAACAACAAGGGCCAAAGAAGCGTACTGTAGCATCATCAAAT  1920
RR_7Bb-CNGC2    CCGCCGGGGCATTTTCAGCAACAACAAGGGCCAAAGAAGCGTACTGTAGCATCATCAAAT  1920

FL16.33-8-CNGC1 ATCCATCCTACCGGCTACAGTGGGTGA                                   1947
RR_7Ba-CNGC1    ATCCATCCTACCGGCTACAGTGGGTGA                                   1947
RR_7Bb-CNGC1    ATCCATCCTACCGGCTACAGTGGGTGA                                   1947
FL16.33-8-CNGC2 ATCCATCCTACCGGCTACAGTGGGTGA                                   1947
RR_7Ba-CNGC2    ATCCATCCTACCGGCTACAGTGGGTGA                                   1947
RR_7Bb-CNGC2    ATCCATCCTACCGGCTACAGTGGGTGA                                   1947
```

**Peptide sequence of CNGC**

```
FL16.33-8-CNGC1 MAKHQGPNNEIQMSITPDVSYDFHSPEALKPTREHGQPQASSKRWNWPAVAGVSSALNPW   60
RR_7Ba-CNGC1    MAKHQGPNNEIQMSITPDVSYDFHSPEALKPTREHGQPQASSKRWNWPAVAGVSSALNPW   60
RR_7Bb-CNGC1    MAKHQGPNNEIQMSITPDVSYDFHSPEALKPTREHGQPQASSKRWNWPAVAGVSSALNPW   60
FL16.33-8-CNGC2 MAKHQGPTNEIQMSITPDVSYDFHSPEALKPTPEHGQPQASSKRRNWPTVAGVSSALNPW   60
RR_7Ba-CNGC2    MAKHQGPTNEIQMSITPDVSYDFHSPEALKPTPEHGQPQASSKRRNWPTVAGVSSALNPW   60
RR_7Bb-CNGC2    MAKHQGPTNEIQMSITPDVSYDFHSPEALKPTPEHGQPQASSKRRNWPTVAGVSSALNPW   60

FL16.33-8-CNGC1 WNKILITSCVIAVFFDPLFFYIPYTSEEKKCMGNDERVQTPALIFRSVTDIIFLVHIIYQ  120
```

```
RR_7Ba-CNGC1     WNKILITSCVIAVFFDPLFFYIPYTSEEKKCMGNDERVQTPALIFRSVTDIIFLVHIIYQ   120
RR_7Bb-CNGC1     WNKILITSCVIAVFFDPLFFYIPYTSEEKKCMGNDERVQTPALIFRSVTDIIFLVHIIYQ   120
FL16.33-8-CNGC2  WNKILVTSCVIAVLFDPLFFYIPYTSEEKKCMGNDERVQTPALIFRSVTDIIFLVHIIYQ   120
RR_7Ba-CNGC2     WNKILVTSCVIAVLFDPLFFYIPYTSEEKKCMGNDERVQTPALIFRSVTDIIFLVHIIYQ   120
RR_7Bb-CNGC2     WNKILVTSCVIAVLFDPLFFYIPYTSEEKKCMGNDERVQTPALIFRSVTDIIFLVHIIYQ   120

FL16.33-8-CNGC1  LYVAIKFAHSRVSEDESLVFRYESVSREQMIQFAKAFSHKLSWRSFLTDVFAILPIPQLL   180
RR_7Ba-CNGC1     LYVAIKFAHSRVSEDESLVFRYESVSREQMIQFAKAFSHKLSWRSFLTDVFAILPIPQLL   180
RR_7Bb-CNGC1     LYVAIKFAHSRVSEDESLVFRYESVSREQMIQFAKAFSHKLSWRSFLTDVFAILPIPQLL   180
FL16.33-8-CNGC2  LYVAIKFALSRVSEDESLVFRYESVSREQMIQFAKAFSHKLSWRSFLTDVFAILPIPQLL   180
RR_7Ba-CNGC2     LYVAIKFALSRVSEDESLVFRYESVSREQMIQFAKAFSHKLSWRSFLTDVFAILPIPQLL   180
RR_7Bb-CNGC2     LYVAIKFALSRVSEDESLVFRYESVSREQMIQFAKAFSHKLSWRSFLTDVFAILPIPQLL   180

FL16.33-8-CNGC1  IVGLFFNIGGNNYLLKRKIVGGFLLIQYVPRIYRIYLSSVAVTESALWVRGVFNFFLYIL   240
RR_7Ba-CNGC1     IVGLFFNIGGNNYLLKRKIVGGFLLIQYVPRIYRIYLSSVAVTESALWVRGVFNFFLYIL   240
RR_7Bb-CNGC1     IVGLFFNIGGNNYLLKRKIVGGFLLIQYVPRIYRIYLSSVAVTESALWVRGVFNFFLYIL   240
FL16.33-8-CNGC2  IVRLFFNIGGNNYLLKRKIVSGFLLIQYVPRIYRIYLSSVAVTESAMWVRGVFNIFLYIL   240
RR_7Ba-CNGC2     IVRLFFNIGGNNYLLKRKIVSGFLLIQYVPRIYRIYLSSVAVTESAMWVRGVFNIFLYIL   240
RR_7Bb-CNGC2     IVRLFFNIGGNNYLLKRKIVSGFLLIQYVPRIYRIYLSSVAVTESAMWVRGVFNIFLYIL   240

FL16.33-8-CNGC1  ASHALGAFWYFCSIQRETSCWYRTCVATNISESKCTFYCHNSTAALITPQFKTTLDEKCI   300
RR_7Ba-CNGC1     ASHALGAFWYFCSIQRETSCWYRTCVATNISESKCTFYCHNSTAALITPQFKTTLDEKCI   300
RR_7Bb-CNGC1     ASHALGAFWYFCSIQRETSCWYRTCVATNISESKCTFYCHNSTAALITPQFKTTLDEKCI   300
FL16.33-8-CNGC2  ASHALGAFWYFCSIQRETLCWYRTCLATNISESKCTFYCHNSAAALMTPQFKTTLDEKCI   300
RR_7Ba-CNGC2     ASHALGAFWYFCSIQRETLCWYRTCLATNISESKCTFYCHNSAAALMTPQFKTTLDEKCI   300
RR_7Bb-CNGC2     ASHALGAFWYFCSIQRETLCWYRTCLATNISESKCTFYCHNSAAALMTPQFKTTLDEKCI   300

FL16.33-8-CNGC1  LKVPYNMTDPPFDFGIFFDALKNDIQGKINVPQKIGYCFWWGLRNLSNFGTSLETSTYLW   360
RR_7Ba-CNGC1     LKVPYNMTDPPFDFGIFFDALKNDIQGKINVPQKIGYCFWWGLRNLSNFGTSLETSTYLW   360
RR_7Bb-CNGC1     LKVPYNMTDPPFDFGIFFDALKNDIQGKINVPQKIGYCFWWGLRNLSNFGTSLETSTYLW   360
FL16.33-8-CNGC2  LKVPYNMTDPPYDFGIFFDALKNDIQGKINVPQKIGYCFWWGLRNLSNFGTNLETSTYLW   360
RR_7Ba-CNGC2     LKVPYNMTDPPYDFGIFFDALKNDIQGKINVPQKIGYCFWWGLRNLSNFGTNLETSTYLW   360
RR_7Bb-CNGC2     LKVPYNMTDPPYDFGIFFDALKNDIQGKINVPQKIGYCFWWGLRNLSNFGTNLETSTYLW   360

FL16.33-8-CNGC1  ENSFAILISIIGLLLFLYLIGNVQIYMQRATTRAEEVREKLRIKKNDIDNWVQNNGIEKD   420
RR_7Ba-CNGC1     ENSFAILISIIGLLLFLYLIGNVQIYMQRATTRAEEVREKLRIKKNDIDNWVQNNGIEKD   420
RR_7Bb-CNGC1     ENSFAILISIIGLLLFLYLIGNVQIYMQRATTRAEEVREKLRIKKNDIDNWVQNNGIEKD   420
FL16.33-8-CNGC2  ENSFAILISIIGLLLFLYLIGNVQIYMQRAVTRAEEVREKLRIKKNDIDNWIQNNGIEND   420
RR_7Ba-CNGC2     ENSFAILISIIGLLLFLYLIGNVQIYMQRAVTRAEEVREKLRIKKNDIDNWIQNNGIEND   420
RR_7Bb-CNGC2     ENSFAILISIIGLLLFLYLIGNVQIYMQRAVTRAEEVREKLRIKKNDIDNWIQNNGIEND   420

FL16.33-8-CNGC1  MKEEIMKNIVEKLENDISADLDDIYSILPPYTRKVVKRFVGMRTLRNVPMLSQVDGRVLK   480
RR_7Ba-CNGC1     MKEEIMKNIVEKLENDISADLDDIYSILPPYTRKVVKRFVGMRTLRNVPMLSQVDGRVLK   480
RR_7Bb-CNGC1     MKEEIMKNIVEKLENDISADLDDIYSILPPYTRKVVKRFVGMRTLRNVPMLSQVDGRVLK   480
FL16.33-8-CNGC2  MKEEIMKNIVEKLENDISADLDDIYSILPPYTRKVVKRFVGMRTLRNVPMLSQVDGRVLK   480
RR_7Ba-CNGC2     MKEEIMKNIVEKLENDISADLDDIYSILPPYTRKVVKRFVGMRTLRNVPMLSQVDGRVLK   480
RR_7Bb-CNGC2     MKEEIMKNIVEKLENDISADLDDIYSILPPYTRKVVKRFVGMRTLRNVPMLSQVDGRVLK   480

FL16.33-8-CNGC1  MMCDYLKPVKYQENTMVFETGVPFDRMVFITDGSMWTYTIATDHQTSHSGKATSGGLVTS   540
RR_7Ba-CNGC1     MMCDYLKPVKYQENTMVFETGVPFDRMVFITDGSMWTYTIATDHQTSHSGKATSGGLVTS   540
RR_7Bb-CNGC1     MMCDYLKPVKYQENTMVFETGVPFDRMVFITDGSMWTYTIATDHQTSHSGKATSGGLVTS   540
FL16.33-8-CNGC2  MMCDYLKPVKYQENTMVFETGVPFDRMVFITDGSMWTYTIATDHQTSHSGKATSGGLVTS   540
RR_7Ba-CNGC2     MMCDYLKPVKYQENTMVFETGVPFDRMVFITDGSMWTYTIATDHQTSHSGKATSGGLVTS   540
RR_7Bb-CNGC2     MMCDYLKPVKYQENTMVFETGVPFDRMVFITDGSMWTYTIATDHQTSHSGKATSGGLVTS   540

FL16.33-8-CNGC1  SSTEAHFLKKGDAYGHMLLQLASSSSTAVPISVANVRCHTKVEAFVLMANDLRNIVTACG   600
RR_7Ba-CNGC1     SSTEAHFLKKGDAYGHMLLQLASSSSTAVPISVANVRCHTKVEAFVLMANDLRNIVTACG   600
RR_7Bb-CNGC1     SSTEAHFLKKGDAYGHMLLQLASSSSTAVPISVANVRCHTKVEAFVLMANDLRNIVTACG   600
FL16.33-8-CNGC2  SSTEAHFLKKGDAYGHMLLQLASSSSTAVPISVANVRCHTKVEAFVLMANDLRNIVTACG   600
RR_7Ba-CNGC2     SSTEAHFLKKGDAYGHMLLQLASSSSTAVPISVANVRCHTKVEAFVLMANDLRNIVTACG   600
RR_7Bb-CNGC2     SSTEAHFLKKGDAYGHMLLQLASSSSTAVPISVANVRCHTKVEAFVLMANDLRNIVTACG   600

FL16.33-8-CNGC1  DLWPAFDYKASQDQAADQMAPPGHFQQQQGPKKRTVASSNIHPTGYSG*              649
RR_7Ba-CNGC1     DLWPAFDYKASQDQAADQMAPPGHFQQQQGPKKRTVASSNIHPTGYSG*              649
RR_7Bb-CNGC1     DLWPAFDYKASQDQAADQMAPPGHFQQQQGPKKRTVASSNIHPTGYSG*              649
FL16.33-8-CNGC2  DLWPAFDYKASQDQAADQMAPPGHFQQQQGPKKRTVASSNIHPTGYSG*              649
RR_7Ba-CNGC2     DLWPAFDYKASQDQAADQMAPPGHFQQQQGPKKRTVASSNIHPTGYSG*              649
RR_7Bb-CNGC2     DLWPAFDYKASQDQAADQMAPPGHFQQQQGPKKRTVASSNIHPTGYSG*              649
```

**Supplementary Fig. S9. Genomic, CDS, and peptide sequence alignment of *WAK* and homoeologous genes from FL 16.33-8-phase1 assembly.**

#### Genomic sequence of *WAK*

```
WAK       ----------------------------------------------------------------------------------------------------
WAK_7-1   ----------------------------------------------------------------------------------------------------
WAK_7-4   TCACAACTTCATACACAAACAATCAAATT-------------------------------------------------------------AGCCGTGCCA    39

WAK       ATGGCCTTACATGGGAGAATGCTTCTTTTGCAAC---TCTCTTTGGCCGCAG---TGCTATTAGCAACGACAATGCTAGCAGCTGCTCAAGCCTCACCGC    94
WAK_7-1   ----------------------------------------------------------------------------------------------------
WAK_7-4   ATGGCCTTACAGGGGAGAATGCATCTTTTGCAACTGATCTCTTTGGCCGCAGTACTGCTGTTAGCATCCACAATGCTAGCAGCTGCTCAATCCCCGCCGC   139

WAK       CTGTCTGCAATGAGAGCTGCGGTGGTGTCTCAGTTCCATATCCATTTGGTCTCACTGATGGTTGTTACCTACATGTGCCGGGACAAGATTCTCAGCAGCC   194
WAK_7-1   ----------------------------------------------------------------------------------------------------
WAK_7-4   CTGACTGCAAGGAAACCTGTGGTGGCGTCTCAATTCCATATCCATTTGGTCTCACTGACGGTTGTTACCTACATGTGCCGGGACAAGATTCACAGCA---   236

WAK       ATTCAAGATCACTTGCAACACCA---CCACCTCGCAACCATCCCTACAGTTCTCGGACAGCGATAACTTCCCCACAAATATTGCCAACATTTTTTTGGAA   291
WAK_7-1   ----------------------------------------------------------------AACTTCCCCACAAATATTGCCAACATTTTTGTGGAA    36
WAK_7-4   ATTCAAGCTCGCTTGCGACACCACCGCCACCTCGCAACCGTCCCTAAATTTCTTGGATAGCTATAACTTCCCCACAAACATTTCCAACATTTTCGCGGGT   336

WAK       GAGAGTGCGCTGCAAGTAATGATGGCCCCGTCCTATAACTGCTACACCAACAATACATACAACCCCTCCATGGACAAAGTCATGGACCTCAATCTTCCGC   391
WAK_7-1   GAGAGTGCGCTGCAAGTAATGATGGCCCCGTCCTATAACTGCTACACCAACAATACATACAACTCCTCCATGGACAAAGTCATGGACCTCAATCTTCCGC   136
WAK_7-4   GAGAGTGCGCTGCAAGTAATGTTGACCACGTCCCGAAACTGCTACAACGACACTGTATACAACTCCTCCTTGCACAGAGCCAGTTACCTATATCTTCCGC   436

WAK       CTTCTTACACCCTCTCCAACAGAAACAAGGTTTATAACCTTGGGTGCAACAAAGTAACGAAACTCATAGGTTATCCTCTTGAAGTTACAGACCCTGAGCA   491
WAK_7-1   CTTCTTACACCCTCTCCGACAGAAACAAGGTTTACAACCTTGGGTGCAACAAAGTAACGAAGCTAATAGGTTATCCTCTTGAAGTAACAGACCCTGAGCA   236
WAK_7-4   CTTCTTACACCCTCGCCGACAAAAACAAGGTTTATAACATTGGCTGCAACAAAGCAACGAAGCTCGTAGGCTATCCTCTTCATGTTACAGACCCTGAGCA   536

WAK       GCTCTTTCAAGTACGGTTCAGCGGTGTAAGCTTGTGCCAAGATGAATTCGGCAAGAGATTCCCCGACACTTGCACCGGATTCGGGTGCTCCGTAAATTCT   591
WAK_7-1   GCTCTTTGAAGTAGGGTTCAGCGGTGTAAGCTTGTGCCAAGATGAATTTGGCAAACGATTCCCCGACACTTGCACCGGATTTGGGTGCTCCGTAAATTCT   336
WAK_7-4   GCTCTTTGAAGTAGCTCTCAGCGGTGTAAGCTTGTGCCAAGATGAATTTGGCAAAAGATTCCCCGAGACTTGCACCGGATTCGGATGCTCAGTAAATTCT   636

WAK       ATCCCTAGCGGACTGCAGAATATTACGGTGGATGTGTGGACACTCGGTACCGGCCTCGGAAAAACGGAGCAGTGGGGACTGAGCTACCCTTGCAGTTACG   691
WAK_7-1   ATCCCTATCGGACTGCAGAATATTACGTTGGGTGTGTGGACACTCGGTACCAGCCTCGGAACAACGGCGCAGTGGGGCCTGAGCTACCCTTGCAGTTACG   436
WAK_7-4   ATCCCTAGTGGACTGCAAAATATTACGGTGGATGTGTGGACACTCGGTACCGCCCTCGGAACAACGGAGCAGTGGGGACTGAGCTACCCTTGTAGTTACG   736

WAK       GCTTCGTTGTGGATGAACGCAACTTCACATTCGCCGGGAATCAAAGTTTTGTTGCACTGAGGTCCGGCGCAATAAAGAAGCTTCCGGTGCTTGCTAATTG   791
WAK_7-1   GCTTCGTTGTGGATGAACGCAACTTCACATTCGCCGGGAATCAAAGTTTTGATGCACTGAGGTCCGGCGCAATAAAGAAGCTTCCGGTGCTTGCTAATTG   536
WAK_7-4   GCTTCGTTGTGGATGAACGCAACTTCACATTCGCCGGGAATCGAAGTTTTGATGAACTGAA---TCGCACACGACATGAGCTTCCGGTGCTTGCTAGTTG   833

WAK       GGCGATTGGGAATGACAACTGCGAAGCAGCAAAGAAGAAGAACGAGACTGCTTTTGCATGCAAGACTGTGAATTCCAAGTGTGTCGACCGTGCTGGTGGT   891
WAK_7-1   GGCGATTGGGAATGACAACTGCGAAGCAGCAAAGAAGAAGAACGAGACTGCTTTTGCATGTAAGACTGTGAATTCCAAGTGTGTCGACCGTGCTGGTGGT   636
WAK_7-4   GGCGATTGGGAATGACAACTGCGAAGCAGCAAAGAAGAAGAACGAGACAGCTTTTGCATGCAAGGGTGAGAATACGGAGTGTGTCGACCGCGC---TGGT   930

WAK       GGTTACTTTTGCCAGTGTGAGGATGGTTACGAAGGGAACCCATACCTCCCCGATAATTGCCTAGGTAATTAATTACAAAAATCATCTTAATTTCGCGTTA   991
WAK_7-1   GGTTACTTTTGCCAGTGTGAGGATGGTTACGAAGGGAACCCATACCTCCTCGATAATTGCCTAGGTAA-TACATACACAAATCATCTTAATTTCTTGTTA   735
WAK_7-4   GGCTACTTCTGCCAGTGTTGGGATGGCTACGAAGGGAACCCATACCTCCTCAAAGATTGCGTAGGTAA-----TACACAAATCATCTTAATTTCGTGTTA  1025

WAK       A-TTTGTGATCTATATATAGGAAAGTGATTGTTTCTATATTTGCTTTATATATAGCTTCCCCT-AATGTGATG--TCTTTTTGTTTCAGATATCAATGAG  1087
WAK_7-1   ATTTTGTGATCTATAT-----AACTGATGGTTTCTAGATTTGATTT--ATACAGCTTCCCCTAAATGTAATATGTCTTTTTGTTTCAGATATCAATGAG   827
WAK_7-4   A-TTTGTGATCTATAT-----AAGGTGATGGTTTCTCGATTTGCCTTTCATATAACTT---AAAAATGTAATGTCTTTTTGTTTCAGATGTCGATGAG  1116

WAK       TGC---AAGCATTCAACCCTTTGCAGTGGTCCTGCAACTTGCATAAACTCAATTGGAAGTTACACCTGTAAATGTCACAAGGGCCATAGAAATGATGACC  1184
WAK_7-1   TGCAATAAGAACTCAACCCTTTGCAGTGGTCCTGCAACTTGCATAAACTCAATTGGAGGTTACACTTGTAAATGTCACAAGGGCTACAGAAATGATGACC   927
WAK_7-4   TGCAAGAATAATTCAACCCTTTGCAGTGGTCCTGCAACTTGGGTAAACTCAATTGGAAGTTACAAATGTAAATGTCACAAGGGCTATAGAAATGATGACA  1216

WAK       ACGACAAGAATAAGTGTGTCCTAATCACTGAAACCTCCAGCAAAAATGACAAAGAAATGAAAATTTCCCTGGGTATGCGCACTCATTTTTTTTTTTCTTT  1284
WAK_7-1   ACGACAAGAATAAGTGTGTCCAAATCACTGAAACCTCCAGCAAAAATGACAAAGAAATGAAAATTTCCTTGGGTATGCACTCTCATCTTTTTTTTT----  1024
WAK_7-4   ACGACAAGAATAAGTGTGTCAAAATCAACGAAACCTCCAGCAAAAATGATACAGAAATGAAAATTTCCTTGGGTATGCATCTCTTTTCTTTTTCTTT--  1314

WAK       TCTCTTTTCTCAGAGCACTTTCATCTCTGTTAATCTATTATTGAC------AAGTTTGTCAATTCTATATATCAATTTCTAAAATTGTCGACTTTTTAAG  1378
WAK_7-1   ----------------------------------------------------------------------------------------------------  1024
WAK_7-4   -------TGTCGAGGCACTCTTAT--------ATTCATTATTGACAAATTAAAATTTGTCAATTTTATCTATCAATTCCTAAATTTTGT----------  1388

WAK       TATTATATTTTTTTTTATACGAACTCTTTAACTATCGTGTTGGAATTTATATTTACCTTAAAAATTGATAAATTTTAAC---------------------  1456
WAK_7-1   ------CTTTTTTTT-------------------------------------------------------------------------------------  1033
WAK_7-4   --CTAACTCTTGTTTA-------------AGTATTATGTTGGTGCTCATATTTACCTTTAAAATTGATAGATTTTAACTTAAAATAATAATAATAATAAT  1473

WAK       ----------AGTTTTTTTTTTTTTATAACAAATCTGCTGGAGATGCTCCTAATATAGTCAAGTTGTAGGAGTGGATTTGAGTACG-TTGTGTTGATCCG  1545
WAK_7-1   ------------------------TAATAATAACTCTGCTAGAGATGCTC----AATAGTCAAGTTGTAGGAATGGATTTGAGTACGTTTGTGCTAATCCG  1106
WAK_7-4   AATAATAATAATAATAATAATAATAATAATAACTCTGCTGGAGATGCTCTTAACATAGTCAAGTTGTAGGAGTGGATTTGAGTACG-TTGTGCTGATCCG  1572

WAK       TTCTTTTGAATTTTTGTAGGTGTCTCTTTGACCTTCCTAGTCATACTGATTATAACTTTTTCGATATACTGCGTACTGAAAAGAAGAAAGTTCAAGCATC  1645
WAK_7-1   ATCTTTTGAATTTTTGCAGGTGTCTCTTTGAGCTTCCTAGTCATATTAATTATAACATTTTCGATATACTGCGTGATGAAAAGAAGAAAGTTCAAGCTAC  1206
WAK_7-4   TTCTTATGAATTTTTGCAGGTGTCTCTTTGAGCTTCCTAGTCATACTGATTATAACTTTTTCGATATACTGCCTACTGAAAAGAAGAAAGTTCAAGCAGC  1672

WAK       TCTGTGACAGGTACTACAAAGACAATGGGGGCTTCTTGTTACCACAAGAAATGCAGAAGTACAAAGGGTCTCAGGCACCCAGAATCTTTAAATTAGAAGA  1745
WAK_7-1   TCTGTGACAAGTACTACAAAGACAATGGGGGCTTCTTGTTACCACAAGAAATGCAGAAGCACAAAGGGTCTCAGGCACCCAGAATCTTTAAATTAGAAGA  1306
WAK_7-4   TATGTGACAGGTACTACGAAGTCAATGGGGGCTTCTTGTTACCACAAGAAATGCAAAAGTACAAAGGGTCTCAGGCACCCAGAATCTTTAAATTAGAAGA  1772

WAK       GCTTAACAAGGCAACCAACAAGTTTGATCCCCATGAAATAATTGGAGAAGGAGGCTTTGGATTGGTTTACAAGGGAACATTGCCGGACAACAAGCAGGAG  1845
WAK_7-1   GCTTAACAAGGCAAGCAGCAAGTTTGATGCCACTAATATAATTGGAGAAGGAGGCTTTGCATTTGTTTACAAGGGAACATTGCCAGATAACAAGCAGGTG  1406
WAK_7-4   GCTTAAGAAGGCAACCAACAAGTTTGATCCCCATGAAATAATTGGAGAAGGAGGCTTTGGATTGGTTTACAAGGGAACATTGCCGGACAACAAGCAGGTG  1872
```

```
WAK      GTTGCCATAAAGAAGTCAAAAACTGATGCTCCCACCATGACCCCCGAAAGTAGGATCGCCCACACTAAGCAGTTCATTAATGAGATGATTGTTCTCTCTG  1945
WAK_7-1  GTTGCCATAAAGAAGTCAAAAACTGATGCTCCCACCATGACCCCCGAAAGTAGGATCGCCCACACTAAGCAGTTCATTAATGAGATGATTGTTCTCTCTG  1506
WAK_7-4  GTTGCCATAAAGAAGTCAAAAACTGATGCTCCCACCATGACCCCCGAAAGTAGGATCGCCCACACTAAGCAGTTCATTAATGAGATGATTGTTCTCTCTG  1972

WAK      GAATCAAGCATAGAAATGTCGTGAGGCTCTTAGGTTGTTGTTTGGAGACCAAAACGCCTATATTAGTGTACGAGTTTGTCTGCAATGGCACACTTTATGA  2045
WAK_7-1  GAATCAAGCATAGAAATGTCGTGAGACTCTTAGGTTGTTGTTTGGAAACCAAAACGCCTATATTAGTGTACGAGTTCGTCAGCAACGGCACACTTTATGA  1606
WAK_7-4  GAATCAAGCATAGAAATGTCGTGAGGCTCTTAGGTTGTTGTTTGGAGACCAAAACGCCTATATTAGTGTACGAGTTGGTCAGCAATGGCACTCTTTATGA  2072

WAK      GCACATTCATAAACAGAAAGGCAAAGGACCTCTTTCCTTTAGGCAACGAGTGAAGATAGCAGCAGAAACAGCAGGATCCCTAGCATACTTGCACTACAAC  2145
WAK_7-1  ACACATTCATAAAAAGAAAGGCAAAGGACCTCTTTCCTTTGGTCAACGAATGAAGATAGCAGCAGAAACAGCAGGATCCCTAGCATACTTGCACTACAAC  1706
WAK_7-4  GCACATTCAGAAAAAGAAAGGCAAAGGACCTCTTTCCTTTGGGCAACGAATGAAGATAGCAGCAGAAACAGCAGGATCCCTAGCATACTTGCACTACAGC  2172

WAK      GCTTCTTCCACGCCAATTCTGCATCGAGACGTAAAGGCATCTAACATCCTACTGGATGAGAATTGCACAGCCAAAATATCAGACTTTGGAGCTTCAAAAT  2245
WAK_7-1  GCTTCTTCCACGCCAATTCTGCATCGAGATGTCAAGGCATCTAACATCCTACTGGATGAGAATTGCACAGCCAAAGTATCAGACTTTGGAGCTTCAAAAT  1806
WAK_7-4  GCTTCTTCCACGCCAATTCTGCACCGAGATGTCAAGGCGACGAACATCCTGCTGGATGAGAATTTGACAGCCAAAGTATCAGACTTTGGAGCTTCAAAAT  2272

WAK      TGGTTCCCGAAGATGAAAACACTCAACTGGCTACTTTGGTGCAAGGGACGCTAGGGTACTTGGATCCTCAATATATTCAAACACACACACTGACGGAGAA  2345
WAK_7-1  TGGTTCCTGAAGATGAAAATACTCAAATGGCTACTTTAGTGCAAGGGACACTAGGGTACTTGGACCCTGAATATCTTCAGTCACACGCACTGACGGAGAA  1906
WAK_7-4  TGGTTCCCGAGGATCAAAATACTCAAATGGCTACTTTGGTGCAAGGGACGCTAGGGTACTTGGACCCTGAATATATTCAGACACACAAACTGACGGAGAA  2372

WAK      GAGTGATGTCTACAGTTTCGGGGTTGTCCTTGTGGAACTGATAACGAGTCAAGCGGCAATTAATTCTAACAAGAAGCTTGAGGCAGAGAAAAACCTAGCC  2445
WAK_7-1  GAGTGATGTCTATAGTTTTGGGGTTGTCCTTGTGGAGCTAATAACGAGTCAAGTGGCAAT---TTCTAACAAGAAGCTTGAGGCAGAGAAAAACCTAGCA  2003
WAK_7-4  GAGTGATGTCTATAGTTTCGGGGTTGTCCTTGTGGAGCTATTAACGAGTCAAATGGCAATTAATTCCAACAAGAAGCTTGAGGAAGAGAGAAACCTAGCG  2472

WAK      AATTGTTTTTTAAAGTCGGTCGAAGATAATCACTTGGATCAGATTCTTGATCATGAAATTATCA---AAGATAGCTTAGAAATAGCTGAACAAGTAGCCC  2542
WAK_7-1  AACTGTTTTTTAAAGTCGGTGGCAGATAATAACTTGGATCAGATTCTTGATCATGAAATTATCAAAGAAGAAAGCTTGCGAAATAGCCGAACAAGTAGCCC  2103
WAK_7-4  AACTGTTTTTTAATGGCTGTGGAAGATAATCACTTGGATCAGATTCTTGATCATGAAATTATCAAAGAAGAAAGCTTTGAAATAGCTGAACAAGTCGCCC  2572

WAK      ATCTCGCCAAAAGATGTTTGAGTTTGAAAGGAGATGATAGGCCTGCCATGAGAGAAGTAGAAACGGAGCTAGTGGCAATATTGGCAGTTATGGAAAAGCG  2642
WAK_7-1  ATCTCGCCAAAAGATGTTTAAGTTTGAAAGGAGAAGATAGGCCTACCATGGAAGAAGTAGAAACGGAGCTAAAGGTAATTTTGGCAGTAATGGCAAAGCA  2203
WAK_7-4  AACTCGCCCAAAGATGTTTAAGTCCGAAAGGAGGAAACAGGCCTACCATGATAGAAGTAGAAACGGAGCTAGGGGCAATATTGGCAGTTATGGCAAAGCA  2672

WAK      TCCCGGAGGAAAGCCTGACTCCTCCCCCAAAGAGACCGATTACTTGCTTGCAGCGTCACCTTCAAATGCTTTCGTTGTTGATGTTAGAAGTGATGAAGGT  2742
WAK_7-1  TCCAGGAGGAAAGCCTGACTCCTCCCCCAAAGAGACAGATTACTTGCTTGCAGCGTCACCTTCAAACGCTTACGTTGTTGATGTTAGAAGTGATGAAGGT  2303
WAK_7-4  TCCAGGAGGAAAGCCCGACTCCTCCCCCAAAGAGACCGATTACTTGCTTGCAGCGTCACCTTCAAATGCTTTCGTTGTTGATGTTAGAAGTGATGAAGGT  2772

WAK      GAACTCACAACCAGCATAGATTATGACCAGAGCATGCAGAATCAATCCCAGATGACGAGGCCTTATGATAGTGGGAGATAG-------------------  2842
WAK_7-1  GAAGTCATAACTAGCATAGATTATGACAAGAGCATGCAGAATCAAGCCCAGATGATGAAGCCTTATGATGGTGGGAGATAGTTCATCAATTGGCAAGCTA  2403
WAK_7-4  GAAGTCATAACCAGCATAGATTATGACAAGAGCATGCATAGTCATGCCCAGATGATGAAGCCTTATGATGGCGGGAGATAGTTCCTCAATTGGCAAGCTA  2872

WAK      ----------------------------------------------------------------------------------------------------  2942
WAK_7-1  GCTTCGAATCTCTTTCATAGCTAGTTATGTATGGAGTTGTTCTTTTGTGTGTGCTATGTTCTCATGATGGTGGGGGATAGTTCATCAATTAGCTAGCTAG  2503
WAK_7-4  GCTTCGAATCTCTTTCATAGCTAGTTATGTATGGAGTTGTTCTTTTGTGTGTGTTATGTTCTCATGATGGTGGGAGATAGTTCATCAATTAGCTAGCTAG  2972

WAK      ----------------------------------------------------------------------------------------------------  3042
WAK_7-1  CTATATAGCTTCGAATTTTCTTTCATAGCTAGCTAGCTAGTTAACTATGGAGTGTTGTTCTTAATTTTCTTTACTACGTACTTTTGAG----TATGCTGT  2599
WAK_7-4  CTATATAGCTTCGAATTTTCTTTCATAGCTAGCTAGCTAGTTAACTATGGAGTGTTGTTCTTAATTTTCTTAACTACGTACTTTTGAGTATATATGCTGT  3072

WAK      --------------------------------------------------------                                              2823
WAK_7-1  GTTTAATGAAAGATATGGATGTAT--------------------------------                                              2623
WAK_7-4  GTTTGAT-AAAGATATGGATGTATGCATGTAATGACAGTAAAATCGATCTGTTGTA                                              3127
```

**CDS of WAK**

```
WAK      ATGGCCTTACATGGGAGAATGCTTCTTTTGCAAC---TCTCTTTGGCCGCAG---TGCTATTAGCAACGACAATGCTAGCAGCTGCTCAAGCCTCACCGC    94
WAK_7-1  ----------------------------------------------------------------------------------------------------
WAK_7-4  ATGGCCTTACAGGGGAGAATGCATCTTTTGCAACTGATCTCTTTGGCCGCAGTACTGCTGTTAGCATCCACAATGCTAGCAGCTGCTCAATCCCCGCCGC   100

WAK      CTGTCTGCAATGAGAGCTGCGGTGGTGTCTCAGTTCCATATCCATTTGGTCTCACTGATGGTTGTTACCTACATGTGCCGGGACAAGATTCTCAGCAGCC   194
WAK_7-1  ----------------------------------------------------------------------------------------------------
WAK_7-4  CTGACTGCAAGGAAACCTGTGGTGGCGTCTCAATTCCATATCCATTTGGTCTCACTGACGGTTGTTACCTACATGTGCCGGGACAAGATTCACAGCA---   197

WAK      ATTCAAGATCACTTGCAACACCA---CCACCTCGCAACCATCCCTACAGTTCTCGGACAGCGATAACTTCCCCACAAATATTGCCAACATTTTTTTGGAA   291
WAK_7-1  ----------------------------------------------------------------------------------------------------
WAK_7-4  ATTCAAGCTCGCTTGCGACACCACCGCCACCTCGCAACCGTCCCTAAATTTCTTGGATAGCTATAACTTCCCCACAAACATTTCCAACATTTTCGCGGGT   297

WAK      GAGAGTGCGCTGCAAGTAATGATGGCCCCGTCCTATAACTGCTACACCAACAATACATACAACCCCTCCATGGACAAAGTCATGGACCTCAATCTTCCGC   391
WAK_7-1  ------------------ATGATGGCCCCGTCCTATAACTGCTACACCAACAATACATACAACTCCTCCATGGACAAAGTCATGGACCTCAATCTTCCGC    82
WAK_7-4  GAGAGTGCGCTGCAAGTAATGTTGACCACGTCCCGAAACTGCTACAACGACACTGTATACAACTCCTCCTTGCACAGAGCCAGTTACCTATATCTTCCGC   397

WAK      CTTCTTACACCCTCTCCAACAGAAACAAGGTTTATAACCTTGGGTGCAACAAAGTAACGAAACTCATAGGTTATCCTCTTGAAGTTACAGACCCTGAGCA   491
WAK_7-1  CTTCTTACACCCTCTCCGACAGAAACAAGGTTTACAACCTTGGGTGCAACAAAGTAACGAAGCTAATAGGTTATCCTCTTGAAGTAACAGACCCTGAGCA   182
WAK_7-4  CTTCTTACACCCTCGCCGACAAAAACAAGGTTTATAACATTGGCTGCAACAAAGCAACGAAGCTCGTAGGCTATCCTCTTCATGTTACAGACCCTGAGCA   497

WAK      GCTCTTTCAAGTACGGTTCAGCGGTGTAAGCTTGTGCCAAGATGAATTCGGCAAGAGATTCCCCGACACTTGCACCGGATTCGGGTGCTCCGTAAATTCT   591
WAK_7-1  GCTCTTTGAAGTAGGGTTCAGCGGTGTAAGCTTGTGCCAAGATGAATTTGGCAAACGATTCCCCGACACTTGCACCGGATTTGGGTGCTCCGTAAATTCT   282
WAK_7-4  GCTCTTTGAAGTAGCTCTCAGCGGTGTAAGCTTGTGCCAAGATGAATTTGGCAAAAGATTCCCCGAGACTTGCACCGGATTCGGATGCTCAGTAAATTCT   597

WAK      ATCCCTAGCGGACTGCAGAATATTACGGTGGATGTGTGGACACTCGGTACCGGCCTCGGAAAAAACGGAGCAGTGGGGACTGAGCTACCCTTGCAGTTACG   691
WAK_7-1  ATCCCTATCGGACTGCAGAATATTACGTTGGGTGTGTGGACACTC-------------------------------------------------------   327
```

```
WAK_7-4   ATCCCTAGTGGACTGCAAAATATTACGGTGGATGTGTGGACACTC-------------------------------------------------------   642

WAK       GCTTCGTTGTGGATGAACGCAACTTCACATTCGCCGGGAATCAAAGTTTTGTTGCACTGAGGTCCGGCGCAATAAAGAAGCTTCCGGTGCTTGCTAATTG   791
WAK_7-1   ----------------------------------------------------------------------------------------------------   327
WAK_7-4   ----------------------------------------------------------------------------------------------------   642

WAK       GGCGATTGGGAATGACAACTGCGAAGCAGCAAAGAAGAAGAACGAGACTGCTTTTGCATGCAAGACTGTGAATTCCAAGTGTGTCGACCGTGCTGGTGGT   891
WAK_7-1   ----------------------------------------------------------------------------------------------------   327
WAK_7-4   ----------------------------------------------------------------------------------------------------   642

WAK       GGTTACTTTTGCCAGTGTGAGGATGGTTACGAAGGGAACCCATACCTCCCCGATAATTGCCTAGATATCAATGAGTGCAAG---CATTCAACCCTTTGCA   988
WAK_7-1   ---------------------------------------------------------------GATATCAATGAGTGCAATAAGAACTCAACCCTTTGCA   364
WAK_7-4   ---------------------------------------------------------------GATGTCGATGAGTGCAAGAATAATTCAACCCTTTGCA   679

WAK       GTGGTCCTGCAACTTGCATAAACTCAATTGGAAGTTACACCTGTAAATGTCACAAGGGCCATAGAAATGATGACCACGACAAGAATAAGTGTGTCCTAAT   1088
WAK_7-1   GTGGTCCTGCAACTTGCATAAACTCAATTGGAGGTTACACTTGTAAATGTCACAAGGGCTACAGAAATGATGACCACGACAAGAATAAGTGTGTCCAAAT   464
WAK_7-4   GTGGTCCTGCAACTTGCGTAAACTCAATTGGAAGTTACAAATGTAAATGTCACAAGGGCTATAGAAATGATGACAACGACAAGAATAAGTGTGTCAAAAT   779

WAK       CACTGAAACCTCCAGCAAAAATGACAAAGAAATGAAAATTTCCCTGGGTGTCTCTTTGACCTTCCTAGTCATACTGATTATAACTTTTTCGATATACTGC   1188
WAK_7-1   CACTGAAACCTCCAGCAAAAATGACAAAGAAATGAAAATTTCCTTG------------------------------------------------------   510
WAK_7-4   CAACGAAACCTCCAGCAAAAATGATACAGAAATGAAAATTTCCTTG------------------------------------------------------   825

WAK       GTACTGAAAAGAAGAAAGTTCAAGCATCTCTGTGACAGGTACTACAAAGACAATGGGGGCTTCTTGTTACCACAAGAAATGCAGAAGTACAAAGGGTCTC   1288
WAK_7-1   ----------------------------------------------------------------------------------------------------   510
WAK_7-4   ----------------------------------------------------------------------------------------------------   825

WAK       AGGCACCCAGAATCTTTAAATTAGAAGAGCTTAACAAGGCAACCAACAAGTTTGATCCCCATGAAATAATTGGAGAAGGAGGCTTTGGATTGGTTTACAA   1388
WAK_7-1   ----------------------------------------------------------------------------------------------------   510
WAK_7-4   ----------------------------------------------------------------------------------------------------   825

WAK       GGGAACATTGCCGGACAACAAGCAGGAGGTTGCCATAAAGAAGTCAAAAACTGATGCTCCCACCATGACCCCCGAAAGTAGGATCGCCCACACTAAGCAG   1488
WAK_7-1   ----------------------------------------------------------------------------------------------------   510
WAK_7-4   ----------------------------------------------------------------------------------------------------   825

WAK       TTCATTAATGAGATGATTGTTCTCTCTGGAATCAAGCATAGAAATGTCGTGAGGCTCTTAGGTTGTTGTTTGGAGACCAAAACGCCTATATTAGTGTACG   1588
WAK_7-1   ----------------------------------------------------------------------------------------------------   510
WAK_7-4   ----------------------------------------------------------------------------------------------------   825

WAK       AGTTTGTCTGCAATGGCACACTTTATGAGCACATTCATAAACAGAAAGGCAAAGGACCTCTTTCCTTTAGGCAACGAGTGAAGATAGCAGCAGAAACAGC   1688
WAK_7-1   ----------------------------------------------------------------------------------------------------   510
WAK_7-4   ----------------------------------------------------------------------------------------------------   825

WAK       AGGATCCCTAGCATACTTGCACTACAACGCTTCTTCCACGCCAATTCTGCATCGAGACGTAAAGGCATCTAACATCCTACTGGATGAGAATTGCACAGCC   1788
WAK_7-1   ----------------------------------------------------------------------------------------------------   510
WAK_7-4   ----------------------------------------------------------------------------------------------------   825

WAK       AAAATATCAGACTTTGGAGCTTCAAAATTGGTTCCCGAAGATGAAAACACTCAACTGGCTACTTTGGTGCAAGGGACGCTAGGGTACTTGGATCCTCAAT   1888
WAK_7-1   ----------------------------------------------------------------------------------------------------   510
WAK_7-4   ------------------------------------------------------------------------GGGACGCTAGGGTACTTGGACCCTGAAT   853

WAK       ATATTCAAACACACACACTGACGGAGAAGAGTGATGTCTACAGTTTCGGGGTTGTCCTTGTGGAACTGATAACGAGTCAAGCGGCAATTAATTCTAACAA   1988
WAK_7-1   ----------------------------------------------------------------------------------------------------   510
WAK_7-4   ATATTCAGACACACAAACTGACGGAGAAGAGTGATGTCTATAGTTTCGGGGTTGTCCTTGTGGAGCTATTAACGAGTCAAATGGCAATTAATTCCAACAA   953

WAK       GAAGCTTGAGGCAGAGAAAAACCTAGGCAATTGTTTTTTAAAGTCGGTCGAAGATAATCACTTGGATCAGATTCTTGATCATGAAATTATCA---AAGAT   2085
WAK_7-1   ----------------------------------------------------------------------------------------------------   510
WAK_7-4   GAAGCTTGAGGAAGAGAGAAACCTAGGGAACTGTTTTTTAATGGCTGTGGAAGATAATCACTTGGATCAGATTCTTGATCATGAAATTATCAAAGAAGAA   1053

WAK       AGCTTAGAAATAGCTGAACAAGTAGCCCATCTCGCCAAAAGATGTTTGAGTTTGAAAGGAGATGATAGGCCTGCCATGAGAGAAGTAGAAACGGAGCTAG   2185
WAK_7-1   ----------------------------------------------------------------------------------------------------   510
WAK_7-4   AGCTTTGAAATAGCTGAACAAGTCGCCCAACTCGCCCAAAGATGTTTAAGTCCGAAAGGAGGAAACAGGCCTACCATGATAGAAGTAGAAACGGAGCTAG   1153

WAK       TGGCAATATTGGCAGTTATGGAAAAGCGTCCCGGAGGAAAGCCTGACTCCTCCCCCAAAGAGACCGATTACTTGCTTGCAGCGTCACCTTCAAATGCTTT   2285
WAK_7-1   --------------------------------GGAGGAAAGCCTGACTCCTCCCCCAAAGAGACAGATTACTTGCTTGCAGCGTCACCTTCAAACGCTTA   578
WAK_7-4   GGGCAATATTGGCAGTTATGGCAAAGCATCCAGGAGGAAAGCCCGACTCCTCCCCCAAAGAGACCGATTACTTGCTTGCAGCGTCACCTTCAAATGCTTT   1253

WAK       CGTTGTTGATGTTAGAAGTGATGAAGGTGAACTCACAACCAGCATAGATTATGACCAGAGCATGCAGAATCAATCCCAGATGACGAGGCCTTATGATAGT   2385
WAK_7-1   CGTTGTTGATGTTAGAAGTGATGAAGGTGAAGTCATAACTAGCATAGATTATGACAAGAGCATGCAGAATCAAGCCCAGATGATGAAGCCTTATGATGGT   678
WAK_7-4   CGTTGTTGATGTTAGAAGTGATGAAGGTGAAGTCATAACCAGCATAGATTATGACAAGAGCATGCATAGTCATGCCCAGATGATGAAGCCTTATGATGGC   1353

WAK       GGGAGATAG                                                                                    2394
WAK_7-1   GGGAGATAG                                                                                     687
WAK_7-4   GGGAGATAG                                                                                    1362
```

**Peptide sequence of WAK**

```
WAK       MALHGRMLLLQL-SLAAV-LLATTMLAAAQASPPVCNESCGGVSVPYPFGLTDGCYLHVPGQDSQQPFKITCNTT-TSQPSLQFSDSDNFPTNIANIFLE   97
WAK_7-1   ----------------------------------------------------------------------------------------------------
WAK_7-4   MALQGRMHLLQLISLAAVLLLASTMLAAAQSPPPDCKETCGGVSIPYPFGLTDGCYLHVPGQDSQQ-FKLACDTTATSQPSLNFLDSYNFPTNISNIFAG   99

WAK       ESALQVMMAPSYNCYTNNTYNPSMDKVMDLNLPPSYTLSNRNKVYNLGCNKVTKLIGYPLEVTDPEQLFQVRFSGVSLCQDEFGKRFPDTCTGFGCSVNS   197
WAK_7-1   ------MMAPSYNCYTNNTYNSSMDKVMDLNLPPSYTLSDRNKVYNLGCNKVTKLIGYPLEVTDPEQLFEVGFSGVSLCQDEFGKRFPDTCTGFGCSVNS   94
WAK_7-4   ESALQVMLTTSRNCYNDTVYNSSLHRASYLYLPPSYTLADKNKVYNIGCNKATKLVGYPLHVTDPEQLFEVALSGVSLCQDEFGKRFPETCTGFGCSVNS   199
```

```
WAK        IPSGLQNITVDVWTLGTGLGKTEQWGLSYPCSYGFVVDERNFTFAGNQSFVALRSGAIKKLPVLANWAIGNDNCEAAKKKNETAFACKTVNSKCVDRAGG  297
WAK_7-1    IPIGLQNITLGVWT--------------------------------------------------------------------------------------  108
WAK_7-4    IPSGLQNITVDVWT--------------------------------------------------------------------------------------  213

WAK        GYFCQCEDGYEGNPYLPDNCLDINEC-KHSTLCSGPATCINSIGSYTCKCHKGHRNDDHDKNKCVLITETSSKNDKEMKISLGVSLTFLVILIITFSIYC  396
WAK_7-1    --------------------LDINECNKNSTLCSGPATCINSIGGYTCKCHKGYRNDDHDKNKCVQITETSSKNDKEMKISL------------------  170
WAK_7-4    --------------------LDVDECKNNSTLCSGPATCVNSIGSYKCKCHKGYRNDDNDKNKCVKINETSSKNDTEMKISL------------------  275

WAK        VLKRRKFKHLCDRYYKDNGGFLLPQEMQKYKGSQAPRIFKLEELNKATNKFDPHEIIGEGGFGLVYKGTLPDNKQEVAIKKSKTDAPTMTPESRIAHTKQ  496
WAK_7-1    ----------------------------------------------------------------------------------------------------  170
WAK_7-4    ----------------------------------------------------------------------------------------------------  275

WAK        FINEMIVLSGIKHRNVVRLLGCCLETKTPILVYEFVCNGTLYEHIHKQKGKGPLSFRQRVKIAAETAGSLAYLHYNASSTPILHRDVKASNILLDENCTA  596
WAK_7-1    ----------------------------------------------------------------------------------------------------  170
WAK_7-4    ----------------------------------------------------------------------------------------------------  275

WAK        KISDFGASKLVPEDENTQLATLVQGTLGYLDPQYIQTHTLTEKSDVYSFGVVLVELITSQAAINSNKKLEAEKNLANCFLKSVEDNHLDQILDHEIIK-D  695
WAK_7-1    ----------------------------------------------------------------------------------------------------  170
WAK_7-4    ------------------------GTLGYLDPEYIQTHKLTEKSDVYSFGVVLVELLTSQMAINSNKKLEEERNLANCFLMAVEDNHLDQILDHEIIKEE  351

WAK        SLEIAEQVAHLAKRCLSLKGDDRPAMREVETELVAILAVMEKRPGGKPDSSPKETDYLLAASPSNAFVVDVRSDEGELTTSIDYDQSMQNQSQMTRPYDS  795
WAK_7-1    --------------------------------------------GGKPDSSPKETDYLLAASPSNAYVVDVRSDEGEVITSIDYDKSMQNQAQMMKPYDG  226
WAK_7-4    SFEIAEQVAQLAQRCLSPKGGNRPTMIEVETELGAILAVMAKHPGGKPDSSPKETDYLLAASPSNAFVVDVRSDEGEVITSIDYDKSMHSHAQMMKPYDG  451

WAK        GR*                                                                                      798
WAK_7-1    GR*                                                                                      229
WAK_7-4    GR*                                                                                      454
```

**Supplementary Fig. S10. Genomic, CDS, and peptide sequence alignment of *CNGC* and homoeologous genes from FL 16.33-8-phase1 assembly.**

**Genomic sequence of *CNGC***

```
CNGC1    ----------------------------------------------------------------------------------------------------
CNGC2    ----------------------------------------------------------------------------------------------------
CNGC_7-2 AAGACAAAAACCAATTTCACGTGTATCATGAAGTTTGGTTTTCTCCAACTCACAGTTTCTCTATGGATGACCATTACTCATCCACCAGTTATAGGCAAGA   100
CNGC_7-4 --GACAAAGACCAATTTCACGTGTATCATGAAGTTTGGTTTTCTCCAACTCACAGTTTTTCTATGGATGACCATTTCTCATCCACCAGTTATAGGCAAGA    98

CNGC1    ----------------------------------------------------------------------------------------------------
CNGC2    ----------------------------------------------------------------------------------------------------
CNGC_7-2 GAGGAAAGAGTTGTTAGGATTGAGATATATCGTCAGCGGTAACAAACAT----------------ATGATAGTGGAAGCAAAAGAAAACGACCAAGACC   184
CNGC_7-4 GAGGAAAGAGTTGTTAGGGTTGAGATATATCGTAAGCGGTAACAAACATAGAAAAACACATGTGAATGACAGTGGAAGCAAAAGAAAACGACCAAGACC   198

CNGC1    ----------------------------------------------------------------------------------------------------
CNGC2    ----------------------------------------------------------------------------------------------------
CNGC_7-2 AAGAACAAGACCCATTGTCTAATAGTGCGAGGGAGGACCTATCTTCTTTCTTTGATTTCATACCACTCGGCCACTATAATAATTCATACAACTTTACCGA   284
CNGC_7-4 AAGAACAAGACCCATTGTCTAACAGTGCGAGGGAGGACCTATCTTCTTTCTTTAATTTCATACCACTCGGACACTATAATAATTCATACAACTTTTCCGA   298

CNGC1    ----------------------------------------------------------------------------------------------------
CNGC2    ----------------------------------------------------------------------------------------------------
CNGC_7-2 ATGAATCATTCATCACTCACCCA-----------------------------------------------------------------------------   307
CNGC_7-4 ATGAATCATTGAACATTCACCAAAAAACAAAATAAAAGGAATCATTGACACCAAAAGGAAAAAGACATGTACCCAGGTGGGTTTGATCTAGTGGTAAATC   398

CNGC1    ----------------------------------------------------------------------------------------------------
CNGC2    ----------------------------------------------------------------------------------------------------
CNGC_7-2 ----------------------------------------------------------------------------------------------------   307
CNGC_7-4 ATGATAGGTCTGCTTATTGGGTGAATGGTTCAGAGATACTCCTGGTAATAGGAGGTCTCCGGCAAGTGAAAAACACCCCATGAGGAGAGGTCTTGACCTT   498

CNGC1    ----------------------------------------------------------------------------------------------------
CNGC2    ----------------------------------------------------------------------------------------------------
CNGC_7-2 ----------------------------------------------------------------------------------------------------   307
CNGC_7-4 GACATGCTCTAGGCCCCGGAGTGACCACCATTTTCTGTAACCAGAAAAAGGAAAAAGACATGTGAATGATAGTGGAAGCAAAAAGAAAACGACCAAGACC   598

CNGC1    ----------------------------------------------------------------------------------------------------
CNGC2    ----------------------------------------------------------------------------------------------------
CNGC_7-2 ----------------------------------------------------------------------------------------------------   307
CNGC_7-4 AAGACCAAGAACAAGACCCATTGTCTAACAGTGCGAGGGAGGACCTATCTTCTTTCTTTAATTTCATACCACTCGGCCACTATAATAATTCATACAACTT   698

CNGC1    ----------------------------------------------------------------------------------------------------
CNGC2    ----------------------------------------------------------------------------------------------------
CNGC_7-2 ----------------------------------------------------------------------------------------------------   307
CNGC_7-4 TTCCGAATGAATCATTGAACATTCACCAAAAAACAAAATAAAAGGAATCATTGACACCAAAAGGAAAAAGACATGTACCCATGTGGGTTTGATCTAGTGG   798

CNGC1    ----------------------------------------------------------------------------------------------------
CNGC2    ----------------------------------------------------------------------------------------------------
CNGC_7-2 ----------------------------------------------------------------------------------------------------   307
CNGC_7-4 TAAATCATGATAGGTCTGCTTATTGGGCGAATGGTTCAGAGATATTCCTGGTAATAAGAGGTCTCTGGCAAGTGAAAAACACCCCATGAGGTGGGGTCTT   898

CNGC1    ----------------------------------------------------------------------------------------------------
CNGC2    ----------------------------------------------------------------------------------------------------
CNGC_7-2 ----------------------------------------------------------------------------------------------------   307
CNGC_7-4 GACCTTGACATGCTCTAGGCCCCGGAGTGACCACCATTTTCTGTAACCAGAAAAAGGAAAAAGACATGTGAATGATAGTGGAAGCAAAAAGAAAACGACC   998

CNGC1    ----------------------------------------------------------------------------------------------------
CNGC2    ----------------------------------------------------------------------------------------------------
CNGC_7-2 -----------------------------------------------------------------------------------AAAAAACAAAATAAAAG   324
CNGC_7-4 AAGACCAAGAACAAGACCCATTAGGGGAGGACATATCTTCTTTATTTAATTTCATACCACTCGGTCACTATAATAATTCATACAAAAAACAAAATAAAAG  1098

CNGC1    ----------------------------------------------------------------------------------------------------
CNGC2    ----------------------------------------------------------------------------------------------------
CNGC_7-2 GAATCATTGATACCAAAAGGAAAAAACTTTACCAAATGAAGTGAATCCTGAGCTTATATCAAACAAACTTTCATGTCAATATAATAGAATGTGGTGCTAT   424
CNGC_7-4 GAATCATTGATACCAAAAGTAAAAAACTTTACCAAATGAAGTGAATCATGAGCTTATATC-AACCAACTTGCATGTCAATATAATAGAATGTGGTGCTAT  1197

CNGC1    ----------------------------------------------------------------------------------------------------
CNGC2    ----------------------------------------------------------------------------------------------------
CNGC_7-2 TGATTCACACTTATAGTTGATCGAGTATAGTGCATGTTTCTCAATCTTTGTTTCGAAGGTGATTCAGTTGCTGCAAAATTTAGTTGGTCGAGTATAATGT   524
CNGC_7-4 TGATCCACACTTATAGTTGATCGAGTATAGTGCATGTTTCTCAATCTTTGTTTCGAAGGCGATTCATTTGCTGCAAAATTTAGTAGATCGAGTATAGTAT  1297

CNGC1    ----------------------------------------------------------------------------------------------------
CNGC2    ----------------------------------------------------------------------------------------------------
CNGC_7-2 ATGATTTCCTTAGCCACTAGGTGACCATGTGAAC----ACTACTATTTGGTTTATAACATCGTATACCCTCAAAAGACTCCAACATATGGGAATCCCCAA   620
CNGC_7-4 ATGATTTTCTTGGTCATTAGGTGATCATATAAATACTAACTACTATTTGGTTTATAACATCGTATATGCATAAAATACCCCAACATATGGGAATCCCCAA  1397

CNGC1    ----------------------------------------------------------------------------------------------------
CNGC2    ----------------------------------------------------------------------------------------------------
CNGC_7-2 CGTA---TGCCAGAAACTAATATCAAAATCAAAATACGTTATGTACACATGTACATGTTGACTGTTGAGGTTATGAAAGTTTTGGATTCAGTTTTCCGGC   717
CNGC_7-4 CGTATGGTGCCAGAAACTAATATCAAAATCAAAATACGTACTGTACACATGTACTT--------------------------------------------  1453

CNGC1    ----------------------------------------------------------------------------------------------------
CNGC2    ----------------------------------------------------------------------------------------------------
CNGC_7-2 TGCGGTTTTTCTTCTTCTTCTTCTTCTTCTTCGTCGAGTACAGAGCAAGTCGGATCCATCTCTATCTCACCGTCTTTCGCTAACACCCACTAACTTCACT   817
CNGC_7-4 ----TTTCTTCTTCTTCTTCTTCTTCTTCGTCGTCAAGTACAGAGCAAGTCGGATCCATCTCTATCTCACCGTCTTTCGCTAACACCCACTAACTTCACT  1549
```

```
CNGC1     ----------------------------------------------------------------------------------------------------
CNGC2     ----------------------------------------------------------------------------------------------------
CNGC_7-2  TGTGCCTAACATCGTTAATTTCAACGACGTGTGGGCAACACGCGCCAATTCAACTGCAGTCTATGGGCACCGCAGTCCATACTAGACGCTCTCAAGAAAC   917
CNGC_7-4  TGTGCCTAACGTTCC-----------------------CACGCGCCAATTCAACTGCAGTCTATGGGCACCGCAGTCCATACTAGATGCTCTCATGAAAC  1626

CNGC1     ----------------------------------------------------------------------------------------------------
CNGC2     ----------------------------------------------------------------------------------------------------
CNGC_7-2  TTTATTATAACTTCACGGATAGTCAAGTTCCTTCCTAGTTTCACTAAACATATCAGGTAAATAATTCTATATAACTTTCATTACCTATGTCCTTACGACG  1017
CNGC_7-4  TTTATTATAATTTCACGAATAGTCAAGTTCCTTC--------------CATATCAGGTAAATAAGTCTATATAACTTTCATTACTTATATCCTTATGATG  1712

CNGC1     ----------------------------------------------------------------------------------------------------
CNGC2     ----------------------------------------------------------------------------------------------------
CNGC_7-2  TAAACATTACTCCTATATAAAAGGGTCTAATGAGAATGAATGATATACTTCTTCCTTCTCCAAAAAAAAAAAAAAAAGAAAGTCTATATAACTTAATCCA  1117
CNGC_7-4  TAAACATTACTCATATATAAATAGATCAAATGATAATGAATGATATACTTCTTTCTCCT---------AAAAAAAAAAAAAGTCTATATAACTTAATCTA  1803

CNGC1     ----------------------------------------------------------------------------------------------------
CNGC2     ----------------------------------------------------------------------------------------------------
CNGC_7-2  GACATAATTGGGTTCCATAGGCTAGCTATTTCAGATTAATCTGGTTTCAAGTTAATGATCAATTCCAGC----TTTCCATCATGCAATCTTATAATGTAT  1213
CNGC_7-4  GACATAATTGGGTTCCATAGGCTATCTATTTCAAATTAATCTGGTTTCAAGTTAATGATCAATTCCAGCTGAATTTCCATCTGTCAATCTTATAATGTAT  1903

CNGC1     ----------------------------------------------------------------------------------------------------
CNGC2     ----------------------------------------------------------------------------------------------------
CNGC_7-2  GCTACTTCGATTCTAGAGCGCCAATCTTTGCTCTATAGCTAGCTACC---TAGTACTACTAATTGTCTAGTTGATCCTGCTTTCCTCAAGTAAGAAAAAA  1310
CNGC_7-4  GCTACTTCAATTCTAGAGCGCCAATCTTTGCTCTATAGCTAGCTACCCAGTAGTACTACTAATTGTCTAGTTGATCCTGCGTTCCACAAGTAAGAAAAAG  2003

CNGC1     ----------------------------------------------------------------------------------------------------
CNGC2     ----------------------------------------------------------------------------------------------------
CNGC_7-2  -----AACATACTCTTTCACTGCATATATCTTCCTTAATTGGTAATTCTGTCTCTATTCTTGTTTGTAAACCGTAAACTAACAATTTTCTAGCTTCCGAC  1405
CNGC_7-4  AGTATAACATACTCTTTCTCTGCATATATCTTCCTTAACTGGTAATTCTGTCTCTATGCTTGATTGTAAACCGTAAACTAACAATTTTCTAGCTTCCGAC  2103

CNGC1     ----------------------------------------------------------------------------------------------------
CNGC2     ----------------------------------------------------------------------------------------------------
CNGC_7-2  CACCGTTGTTCCTCTTTCTAGCTAGATCTAAACCACATGCACCAGTCATTGCTGTGACTAATTTTCATA--TTTATATTTTTCAAATTTCAGTACTGCGG  1503
CNGC_7-4  CACCGTTGTTCCTCTTTCTAGCTAGATCTAAACCACATGCACCAGTCATTGCTGTAACTAATTTTCATATTTTTTTTTTTTCAAATTTCAGTACTGCGG  2203

CNGC1     ----------------------------------------------------------------------------------------------------
CNGC2     ----------------------------------------------------------------------------------------------------
CNGC_7-2  ATTGGAAAATTGCTCGAGTACGAAGACTACCATCGCGATCGGCTGTGTTAGTAATTCTGAGACCATCTCATCCAGCATAACAGAAAGGCATAAAATAATT  1603
CNGC_7-4  ATTGGAAAATTGCTCGAGTACGAAGACTACCATCGCGATTGGCTGTGTTAGTAATTCTGAGACCATCTCATCCAGCATAACAGAAAGGCATAAAATAATT  2303

CNGC1     ATGGCTAAACATCAGGGACCAAACAATGAAATCCAAATGAGGTTTGTTTTATTAACCTAGCAATAAGTTAGTTTTCTGCAATGGT--ACTGTTTCTTTTC    98
CNGC2     ATGGCTAAACATCAGGGACCAACCAATGAAATCCAAATGAGGTTTGTTTTATTAACCTAGCAATAAGTTAGTTTTCTGCAATAGT--ACTGTTTCTTTTC    98
CNGC_7-2  ATGGCTAACCATCAGGGACCAAACAATGAAATCCAAATGAGGTTTGTTTTATTAACCTAGCAATAAGTTGGTTTTCTGCAATAGT--ACTGTTTCTTTTC  1701
CNGC_7-4  ATGGCTAAC---CAGGGACCAAACAATGAATTCCAATTGAGGTTTGTTTTATTGACCTAGTAATAAGTTAGTTTTCTGCAATAGTACACCGTTTCTTTTC  2400

CNGC1     CTTTAATTTGGAAAGGATCGACTTCTCCGTTCGAGAGGGTATCGGCTCTAGATCTGATAAATTTGACGGCTTCGATTCATTTAATTTACATTTTTTTTGT   198
CNGC2     CTTTAATTTGGAAAGGATCGACTTCTCCGTTCGAGAGGGTATCGGCTCTAGATCTGATAAATTTGACGGCTTCGATTCATTTAATTTACATTTTTTTTGT   198
CNGC_7-2  CTTTAATTTGGAAAGGGTCGACTTCTCCGTTCGAGAGGGTATCGACTCTGA--TAAATTTATTTGACGGCTTCGATTCATTTAATTTACATTGTTTTTGT  1799
CNGC_7-4  CTTTAATTTGGAAAGGATCGACTTCTCCGTTCGAGAGGGTATCGACTCTGA--TAAATTTATTTGACGGCTTCGATTCATTTAATTTACATTGTTTTTGT  2498

CNGC1     TTTTCTTCTATCAGCATTACACCAGATGTGTCTTATGATTTCCATAGTCCTGAGGC---TCTAAAACCGACACGCGAACATGGACAACCTCAAGCAAGCA   295
CNGC2     TTTTCTTCTATCAGCATTACACCAGATGTGTCTTATGATTTCCATAGTCCTGAGGC---TCTAAAACCGACACCCGAACATGGACAACCTCAAGCAAGCA   295
CNGC_7-2  TTTTCTTCTATCAGCATTACACCAGATGTGTCTTATGATTTCGGTAGTCCTAAGGCTCATCGAAAACCGACACCCGAACATGGACAACCTCAAGCAAGCA  1899
CNGC_7-4  TTTTCTTCTATCAGCATTACACCAGATGTGTCTTATGATTTCCATAGTCCTGCGGC---TCTAAAACCGACACCCAAACATGGACAACCTCAAGCAAGTA  2595

CNGC1     GCAAAAGATGGAATTGGCCAGCAGTTGCCGGAGTTTCTTCAGCTTTGAACCCATGGTGGAATAAGATACTGATAACTTCATGTGTGATTGCAGTTTTTTT   395
CNGC2     GCAAAAGACGGAATTGGCCAACAGTTGCCGGAGTTTCTTCAGCTTTGAACCCATGGTGGAATAAGATACTGGTAACTTCATGTGTGATTGCAGTTTTATT   395
CNGC_7-2  GCAAAAGATGGAATTGGCCAACGGTTGCCGGAGTTTCTTCAGCTTTGAACCCATGGTGGAATAAGATACTGGTAACTTCATGTGTGATTGCAGTTTTATT  1999
CNGC_7-4  GCCAAAGATGGAATTGGCCAATAGTTGCCGGAATTTCTTCAGCTTTGAACCCATGGTGGAATAAGATACTGGTAACTTCATGTGTGATTGCAGTTTTATT  2695

CNGC1     TGATCCCTTGTTCTTTTACATTCCATACACCAGCGAGGAAAAAAAGTGTATGGGAAACGACGAAAGAGTGCAGACTCCAGCTCTGATTTTCCGATCAGTC   495
CNGC2     TGATCCCTTGTTCTTTTACATTCCATACACCAGCGAGGAAAAAAAGTGTATGGGAAACGACGAAAGAGTGCAGACTCCAGCTCTGATTTTCCGATCAGTC   495
CNGC_7-2  TGATCCCTTGTTCTTTTACATTCCATACACCAGCGAGGAAAAAAAGTGTATGGGAAACGACGAAAGAGTGCAGACTGCAGCTCTGATTTTCCGATCAGTC  2099
CNGC_7-4  TGATCCCTTGTTCTTTTACATTCCATACACCAGCGAGGAAAAAAAGTGTATGGGAAACGACGAAAGAGTGCAGACTGCAGCTCTGATTTTCCGATCCGTC  2795

CNGC1     ACAGACATCATTTTCCTGGTGCATATCATATACCAACTATATGTGGCTATCAAATTTGCACACTCAAGGGTTTCCGAGGATGAAAGTTTGGTTTTTAGAT   595
CNGC2     ACAGACATCATTTTCCTGGTGCATATCATATACCAACTATATGTGGCTATCAAATTTGCACTCTCAAGGGTTTCCGAGGATGAAAGTTTGGTTTTTAGAT   595
CNGC_7-2  ACAGACATCACTTTCCTGGTGCATATCATATACCAACTATATGTGGCTATCAAATTTGCACTCTCAAGGGT------GGATGAAACTTTGGTTTTTAGAT  2193
CNGC_7-4  ACAGACATCACTTTCCTGGTGCATATCATATACCAACTATATGTGGCTATCAAATTTGCACTCTCAAGGGTTTCAGAGGATGAAAGTTTGGTTTTTAGAT  2895

CNGC1     ATGAATCGGTATCCCGAGAGCAAATGATTCAGTTTGCGAAGGCATTTTCTCACAAGTTGTCATGGCGATCTTTCCTAACTGACGTTTTCGCTATTTTGCC   695
CNGC2     ATGAATCGGTATCCCGAGAGCAAATGATTCAGTTTGCGAAGGCATTTTCTCACAAGTTGTCATGGCGATCTTTCCTAACTGACGTTTTCGCTATTTTGCC   695
CNGC_7-2  ATGAATCGGTATCCCGAGTGCAAATGATTCGGTTTGCAAAGGCATTTTCTCTCAAGTTGTCATGGCGATCTTTCCTAACTGACGTTTTCGCTATTTTGCC  2293
CNGC_7-4  ATGAATCGGTATCCCGAGAGCAAATGATTCAGTTTACAAAGGCATTTTCTCACAAGTTGTCATGGCGATCTTTCCTAACTGACGTTTTCGCTATTTTGCC  2995

CNGC1     CATACCACAAGTAAGACATATCGTAACTCTCTTTCTAGTTATTAGCTAGAGATTGG---TTAATTAACTGGCTGTAATTTATGGTCTTTTT-AAAGTTTT   791
CNGC2     CATACCACAAGTAAGACATATCGTAACTCTCTTTCTAGTTATTAGCTAGAGATTGG---TTAATTAACTGGCTGTAATTTATGGTCTTTTT-AAAGTTTT   791
CNGC_7-2  CATACCACAAGTAAGACGTATCGTAACTCTCTTTCTAGTTATTAGCTAGAGATTGGCTAATAATTAACTGGCTGTAATTTACGGTCTTTTTAAAAGTTTT  2393
CNGC_7-4  CATACCACAAGTAAGACATATCGTAACTCTCTTTCTAGTTATTAGCTAGAGATTGG---ATAATTAACTGGCTGTAATTTATGGTATTTTT-AAAGTTTT  3091

CNGC1     AACCAACGATTTTTTGTCCAGCTTCTAATAGTAGGATTGTTTTTCAACATCGGAGGCAATAACTACTTGTTAAAAAGAAAGATCGTGGGCGGTTTTCTTC   891
CNGC2     AACCAACGATTTTTTGTCCAGCTGCTAATAGTAAGATTGTTTTTCAACATTGGAGGCAATAACTACTTGTTAAAAAGAAAGATTGTGAGCGGTTTTCTTC   891
CNGC_7-2  AACCCACGATTTTTTGTCCAGCTGCTAATAGTAGGATTGTTTTTCAACATCGGAGGCAATGACTACTTGTTAAGAAGAAAGATTGTGAGTGGTTTTCTTC  2493
```

```
CNGC_7-4  AACCCTCGATTTTTTGTCCAGCTGCTAATAGTAGGATTGTTTTTCAAAATCGGAGGCAATGACTACTTGTTAAAAAGAAAGATTCTGAGCGGTTTTCTTC  3191

CNGC1     TAATCCAATATGTGCCAAGAATCTATCGAATTTATCTTTCATCTGTCGCGGTCACAGAATCTGCATTGTGGGTTAGAGGTGTATTTAACTTCTTCCTCTA   991
CNGC2     TAATCCAATATGTGCCAAGAATCTATCGAATTTATCTTTCATCTGTCGCGGTCACAGAATCTGCAATGTGGGTTAGAGGTGTATTTAACATTTTTCTCTA   991
CNGC_7-2  TAATCCAATTTGTGCCAAGAATCTATCGAATCTATCTTTCATCTGTCGCGATCACAGAATCTACATTGTGGGTTAGAGGTGTATTTAATCTTTTCCTCTA  2593
CNGC_7-4  TAATCCAATTTGTGCCAAGAATCTATCGAATTTATCTTTCATCTGTCGCTATCACAGAATCTGCGTTGTGGGTTAGAGGTGTATTTAACCTTTTTCTCTA  3291

CNGC1     CATTCTTGCGAGTCATGTAAGTTTATCCATC----TCACCTCTTCTCATCAATTC---ATTATGT-----------TTTTTTTTTCATTGAATTAGGTGA  1073
CNGC2     CATTCTTGCGAGTCATGTAAGTTTATCCATC----TCACCACTTCTCATCAATTC---ATTATG-------------TTTTTTTTCATTGAATTAGGTGA  1071
CNGC_7-2  CATCCTTGCGAGTCATGTAAGTTTATCCATCTCGATCACCTCTTCTCATCAATTTTTGATTATGTTTTTTTTTTTTTCTCTTTTCATTGAATTAGGTGA  2693
CNGC_7-4  CATCCTTGCGAGTCATGTAAGTTTATCCATC----TCACCTCTTCTCATCAATTC---ATTATATA------TGTTTTTTTTTTTCATTGAATTAGGTGA  3378

CNGC1     GTTGCTTGAGTTAAGTTTGTGTTGATCAAAAATGTTTGTTGTTAATTATAGGCACTTGGAGCCTTTTGGTACTTTTGTTCTATTCAGCGAGAGACATCAT  1173
CNGC2     GTTGCTTGAGTTAAGTTTGTGTTGATCAAAAATGTTTGTTGTTAATTATAGGCACTTGGAGCCTTTTGGTACTTTTGTTCTATTCAGCGAGAGACATTAT  1171
CNGC_7-2  GTTGCTTGAGTTAAGTTTGTGTTGATAAAAAATGTTTGTTGTTAATTATAGGCACTTGGAGCCTTTTGGTACTTTTTTTCTATTCAGCGACAAACATCAT  2793
CNGC_7-4  GTTGCTTGAGTTAAGTTTGTGTTGATAAAAAATG--TGTTGTTAATTATAGGCACTTGGAGCCTTTTGGTATTTTTTTTCTATTCAGCGAGAGACGTCAT  3476

CNGC1     GTTGGTATCGAACTTGCGTCGCTACAAATATATCAGAATCAAAATGCACCTTTTACTGCCATAACAGTA---CTGCTGCACTTATA------ACCCCACA  1264
CNGC2     GTTGGTATCGAACTTGCCTCGCTACAAATATATCAGAATCAAAATGCACCTTTTACTGCCATAACAGTG---CTGCTGCACTTATG------ACCCCACA  1262
CNGC_7-2  GTTGGTATCGAACTTGCCTCGATACAAATCGATCAGAATTAGAATGCACCTTTTACTGCGATGGCAGTA---CTGCTGCACTTATAACCGATACCCCACA  2890
CNGC_7-4  GTTGGTATCGAACTTGCCTCGATACAAACCCATCAGAATCAGAATGCACTTTTTATTGCCATGACAGTACTGCTGCTGCACTTATA------ACCCCAGA  3570

CNGC1     ATTCAAGACAACTCTAGACGAGAAATGCATTCTAAAAGTTCCATATAACATGACCGATCCACCATTTGATTTTGGAATATTTTTTGATGCCCTCAAAAAT  1364
CNGC2     ATTCAAGACAACTCTAGACGAGAAATGCATTCTAAAAGTTCCATATAACATGACCGATCCACCATATGATTTTGGAATATTTTTTGATGCCCTCAAAAAT  1362
CNGC_7-2  ATTCAATACAACTCTAGACGAGCAATGCGTTCTAAAAGTTCCATATAACATGACCGATCCACCATTTGATTTTGGAATATTTTTTGATGCCCTCAAAAAT  2990
CNGC_7-4  ATTCAATAGAACTCTAGACGAGAAATGCATTCTAAAAGTTCCATATAATATGACTGATCCACCATTTGATTTTGGAATATTTTTTGATGCCCTCAAGAAT  3670

CNGC1     GATATCCAAGGGAAAATAAATGTTCCACAAAAGATTGGTTACTGTTTTTGGTGGGGCTTGCGAAACTTGAGGTAAATCTTCATTTATAACTTAATTCAGT  1464
CNGC2     GATATCCAAGGGAAAATAAATGTTCCACAAAAGATTGGTTACTGTTTTTGGTGGGGCTTGCGAAACTTGAGGTAAATCTTCATTTATAACTTAATTCAGT  1462
CNGC_7-2  GATATCCAAGGGAAAATAAATGTTCCACAAAAGATTGGTTACTGTTTTTGGTGGGGCTTGCGAAACTT-AGGT-AATCTTCATTTATAACTTAATTCAGT  3088
CNGC_7-4  GATATCCAAGGGAAAATAAATGTTCCACAAAAGATTGGTTACTGTTTTTGGTGGGGCTTGCGAAACTTGAGGT-AATCTTCATTTATACCTTAATTCAGT  3769

CNGC1     CGGGTCATTCATCAACCAGCATTTTACAATTACTAATCATTTAACAATTTCTTCTGTGTCCGGCAGTAATTTTGGTACAAGTTTGGAAACAAGTACCTAT  1564
CNGC2     CGGGTCATTCATCAACCAGCATTTTACAATTACTAATCATTTAACAATTTCTTCTGTGTCCGGCAGTAATTTTGGTACAAATTTGGAAACAAGTACCTAT  1562
CNGC_7-2  CGGGTCATTCATCAACCAGCATTTTACAATTACTAATCATTTAACAATTTCTTCTGTGGCCGGCAGTAATTTTGGTACAAGTTTAACAACGAGTACCTAT  3188
CNGC_7-4  CGGGTCATTCATCAACCAGCATTTTACAATTACTAATCATTTAACAATTTCTTCTGTGGCCGGCAGTAATTTTGGTACAAGTTTAACAACAAGTACCTAT  3869

CNGC1     CTGTGGGAAAACAGCTTTGCCATTTTAATTTCTATCATCGGCTTGCTGTTGTTTTTATACCTCATTGGAAATGTACAGGTTAGTAACCTAATCGATCACC  1664
CNGC2     CTGTGGGAAAACAGCTTTGCCATTTTAATTTCTATCATCGGCTTGCTGTTGTTTTTATACCTCATTGGAAATGTACAGGTTGGTAACCTAATCGATCACC  1662
CNGC_7-2  CTGTGGGAAAACAGCTTTGCCATTTTAATTTCTATCATCGGCTTGCTATTGTTTTTATACCTCATTGGAAATGTACAGGTTGGTAACCTAATCGATCACC  3288
CNGC_7-4  CTGTGGGAAAACAGCTTTGCCATTTTAATTTCTATCATCGGCTTGCTATTGTTTTTATATCTCATTGGAAATGTACAGGTTGGTAACCTAATCGATCACC  3969

CNGC1     CATATATGTTATGTTCTTTTTAGGATTTGTT----------------CATTTTAATTTTCCGTAAAT-------TATTAACTTTGTTTTTC---------  1732
CNGC2     CATATATGTTATGTTCTTTTTAGGATTTGTT----------------CATTTTAATTTTCCACAAAT-------TCTTAACTTTGTTTTTC---------  1730
CNGC_7-2  CATATATGTTATGTTACTTATGTA----------------------------------------------------------------------------  3312
CNGC_7-4  CATATATCATATGTTACTTATATGATATGTTACTTATGTATATACAATATTATAGTCCGTTTCAGATATGGACGTCTGCACCGTGCGCTACCGTACGACG  4069

CNGC1     ---ATGAAGAAAAGGTTTAATTTTAGATTTCTTTCAAAAGAAAAAAGTTCAATTTTAGATTATAAATATGTGATGA-----------TGCCATGTTATTT  1818
CNGC2     ---ATGAAGAAAAGATTTAATTTTAGATTTCTTTCAAAAGAAAAAAGTTTAATTTTAGATTATAAATATGTGATGA-----------TGCCATGTTATTT  1816
CNGC_7-2  ----------------------------------------------------------------------------------------------------  3312
CNGC_7-4  CCGGTGAGGGAGCGCCTCCACTCCAG--------CGGACCCCAGCAGCTTCCCTGACCATTTTCTATCCTCGGTGAGTCGGCCGAATTCCAGTTTTCCGG  4161

CNGC1     TCACATCACTTGGTGTTGGGTGTTTACATCATTGAAACAATAAATCTAAA------------------------------CCGTTGAACATTTTAAAATT  1888
CNGC2     TCACATCAGTTGGTGTTGGGTGTTTACATCATTCAAACAATAAATCTAAA------------------------------CCGTTGAACATTTTAAAATT  1886
CNGC_7-2  ----------------------------------------------------------------------------------------------------  3312
CNGC_7-4  TCATATTGACTTTGTTTGAAGTTTTGGATGATCTGGCCGAAAAACTGGAATTCGGCAGACTCGCCGGGGATGGAAAGTGGTCGGGGAAGTTGCTGGAGTC  4261

CNGC1     TAATGGCTCAGAA---------------------ACGGTGTCAAAATTTATGTA-----AAATTTTATGTCTTTTGAACCCGACCCTATATATATATATA  1962
CNGC2     TAATGGCTCAGAA---------------------ACGGTGTCAAAATTTATGTA-----AAATTTTATGTCCTTTGAACCGGACCCTATATATATATAT-  1959
CNGC_7-2  --------------------------------------------------------------------------TATACAGTATTATATATATATATATA  3338
CNGC_7-4  CGCTAGGGTGGAGGCGCGCCCTCGCTGGTGCCATACGGTGGTGAACGGTATGGACGTCCGCACCTGATAACTGTTGTACAGTATTATATATATATATATA  4361

CNGC1     TATATATATATATACACACGCTATTGTGCATAGCAATGGTTAATTTAATTTGATCGAGGAGGTGATTAAATTGATGGATGGAGTT---------------  2047
CNGC2     ---------------ACACGATATTGTGCATAGCAATGGTTAATTTAATTTGATCGAGGAGGTGATTAAATTGATGGATGGAGTTATTAATTACAGGGGC  2044
CNGC_7-2  TATATATATATAT--A-TTGCTATTGTACATAGCAATGGTTAACTTAATTTGATCGAGGAGGTGATTAAATTGATGGATGGAGTT---------------  3420
CNGC_7-4  TATATATATATAT--ATTTGCTATTGTGCATAGCAATGGTTAATTTAATTTGATCGAGGAGGTGATTAAATTGATGGATGGAGTT---------------  4444

CNGC1     ----------------------------------------------------------------------------------------------------  2047
CNGC2     GGAGCCACACCAAGACCAAGGGGTGCCATGGCACCCCCACACATTTTGATGATGGTCAATTACGCCACTGTTGTAGGATTAAATGGTTACAATTGTATAA  2144
CNGC_7-2  ----------------------------------------------------------------------------------------------------  3420
CNGC_7-4  ----------------------------------------------------------------------------------------------------  4444

CNGC1     ----------------------------------------------------------------------------------------------------  2047
CNGC2     TCTGGCCCCCCTTTTTATGTAATAATATGTCACGTAATAATTACTAATTTGATATTTTGGACTATATAATCTCCTTTTTTGTTGCATCTTGCCATATTCA  2244
CNGC_7-2  ----------------------------------------------------------------------------------------------------  3420
CNGC_7-4  ----------------------------------------------------------------------------------------------------  4444

CNGC1     ----------------------------------------------------------------------------------------------------  2047
CNGC2     TCACCATATCGATTAGTTATTAACTAATCATTTTACATGCATGTAAACTTTAAATATATAATCATTTACATATACATAAATTAAACTATATATACATCAA  2344
CNGC_7-2  ----------------------------------------------------------------------------------------------------  3420
CNGC_7-4  ----------------------------------------------------------------------------------------------------  4444

CNGC1     ----------------------------------------------------------------------------------------------------  2047
CNGC2     CAAAAACAATAAATTCAATTTTAATTAACATATAATTTTGACAGACAATTTAAAATTTTATATATATTTTTTCGTATTTTATATCTCGGCCCCCCTAGAC  2444
```

```
CNGC_7-2   ----------------------------------------------------------------------------------------------------   3420
CNGC_7-4   ----------------------------------------------------------------------------------------------------   4444

CNGC1      -----------------------ATTAATTATATGCAACGTGCAGATTTATATGCAGAGGGCAACTACCAGAGCGGAGGAGGTAAGAGAGAAGCTTCGGA   2124
CNGC2      TTATAGTTCTGGCTCCGCCCCTGATTAATTATATGCAACATGCAGATTTATATGCAGAGGGCAGTTACCAGAGCGGAGGAGGTAAGAGAGAAGCTTCGGA   2544
CNGC_7-2   -----------------------GTTAATTATATGCAACATGCAGATTTATATGGAGAGGGCAACTACCAGAGCGGATGAGGTAAGAGAGAAGCTTCGGA   3497
CNGC_7-4   -----------------------AATAATTATATGCAACATGCAGATTTATATGGAGAGGGCAACTACCAGAGCGGAGGAGGTAAGAGAGAAGCTTCGGA   4521

CNGC1      TCAAAAAGAACGATATAGATAATTGGGTCCAAAATAATGGTATCGAGAAGGACATGAAGGAAGAGATCATGAAAAACATCGTTGAAAAGTTGGAAAACGA   2224
CNGC2      TCAAAAAGAACGATATAGATAATTGGATCCAAAATAATGGTATCGAGAATGACATGAAGGAAGAGATCATGAAAAACATCGTTGAAAAGTTGGAAAACGA   2644
CNGC_7-2   TCAAAAAGAACGATATAGATGATTGGATCGAAAATAATGGTATCGGGAAGGACATGAAGGAAGAGATCATGAAAAACATCAAGGAGAAGTTGGAAAAGGA   3597
CNGC_7-4   TAAAACAGAACGATATAGATAATTGGATCGAAAATAATGGTCTCGAGAAGGACATGAAGGAAGAGATCATGAAAAACATCAATGAGAAGTTGGAAAAGGA   4621

CNGC1      CATAAGTGCAGATCTGGATGATATATACTCTATTCTTCCCCCGTATACCAGGAAAGTTGTGAAGCGATTTGTCGGCATGAGGACGCTAAGAAATGTATGT   2324
CNGC2      CATAAGTGCAGATCTGGATGATATATACTCTATTCTTCCCCCGTATACCAGGAAAGTTGTGAAGCGATTTGTCGGCATGAGGACGCTAAGAAATGTATGT   2744
CNGC_7-2   CATAAGTGCAGATCTGGATGATATATACTCTATTCTTCCCCCGAATACCAGGAAAGTTGTGAAGCGATTTGTCGGCATGAGGACGCTAAGAAATGTATGT   3697
CNGC_7-4   CATAAATGCAGATCTGGATGATATATACTCTATTCTTCTCCCGTATACCAGGAAAGTTGTGAAGCGATTTGTCGGCATGAGGACGCTAAGAAATGTATGT   4721

CNGC1      CACAGTACCCTTGTTTGTGACTTGTGTCATCAGTACGTTGTGTGTGTTGTTGAAGCTAGCTTACTAATCAGTGGTCTGCTTATACATACATATGTAGGTA   2424
CNGC2      CACAGTACCCTTGTTTGTGACTTGTGTCATCAGTACGTTGTGTGTGTTGTTGAAGCTAGCTTACTAATCAGTGGTCTGCTTATACATACATATGTAGGTA   2844
CNGC_7-2   CACAGTACCCTTGTTTGTGACTTGTGTCATCAGTACGTTGTGTGTGTTGTTGAAGCTAGCTTACTAATCAGTGGTCTGCTCATACATACATATGTAGGTA   3797
CNGC_7-4   CACAGTACCCTTGTTTGTGACTTGTGTCATCAGTACGTTGTGTGTGTTGTTGAAGCTAGCTTACTAATCAGTGGTCTGCTTATACATACATATGTAGGTA   4821

CNGC1      CCAATGCTTAGCCAGGTGGATGGAAGAGTTTTGAAGATGATGTGCGACTATCTGAAGCCGGTCAAGTACCAGGAGAATACAATGGTTTTTGAAACGGGAG   2524
CNGC2      CCAATGCTTAGCCAGGTGGATGGAAGAGTTTTGAAGATGATGTGCGACTATCTGAAGCCGGTCAAGTACCAGGAGAATACAATGGTTTTTGAAACGGGAG   2944
CNGC_7-2   CCAATGCTTAGGGAGGTGGATGGAAGAGTTTTGAAGATGATGTGCGACTATCTGAAGCCGGTCAAGTTCCAGGAGAATACAATGGTTTTAAAAATGGGAA   3897
CNGC_7-4   CCAATGCTTAGCCAGGTGGATGGAAGAGTTTTGAAGATGATGTGCGACTATCTGAAACCGGTCAAGTACCAGGAGAATACAATGGTTTTTAAAATGGGAG   4921

CNGC1      TACCATTTGATAGAATGGTCTTCATTACAGATGGGTCAATGTGGACTTACACGATTGCTACTGATCATCAGACTAGTCATTCTGGAAAAGCAACGTCAGG   2624
CNGC2      TACCATTTGATAGAATGGTCTTCATTACAGATGGGTCAATGTGGACTTACACGATTGCTACTGATCATCAGACTAGTCATTCTGGAAAAGCAACGTCAGG   3044
CNGC_7-2   AACCATTTGATAGAATGGTCTTCATTACAGATGGGTCAATGTGGACTTAC---------ACTGATCATCAGACTAGTGATTCTGGAAAAGCAACGTCAGG   3988
CNGC_7-4   TACCATTTGATAGAATGGTCTTCATTACAGATGGGTCAATGTGGACTTACACGACTGCTACTGATCATCAGACTAGTCATTCTGGAAAAGCAACGTCAGG   5021

CNGC1      AGGACTAGTTACTTCTTCGTCCACGGAAGCTCACTTCCTCAAGAAGGGTGATGCTTATGGACACATGCTTCTGCAATTAGCATCATCATCTTCTACGGCG   2724
CNGC2      AGGACTAGTTACTTCTTCGTCCACGGAAGCTCACTTCCTCAAGAAGGGTGATGCTTATGGACACATGCTTCTGCAATTAGCATCATCATCTTCTACGGCG   3144
CNGC_7-2   AGGACTAGTTACTTCTTCGTCCACGGAAGCTCACTTCCTCGAGAAGGGTGATGCTTATGGACACATGCTTCTGCAATTAGCATCATCATCTTTTACGACG   4088
CNGC_7-4   AGGACTAGTTACTTCTTCGTCCACGGAAGCTCACCTCCTCGAGAAGGGTGATGCTTATGGACACATGCTTCTGCAATTAGCATCATCATCTTTTACGGCG   5121

CNGC1      GTTCCTATCTCAGTTGCAAATGTCAGGTGCCACACAAAGGTAGAGGCGTTTGTCCTCATGGCCAATGACTTGAGAAACATAGTAACTGCG---------T   2815
CNGC2      GTTCCTATCTCAGTTGCAAATGTCAGGTGCCACACAAAGGTAGAGGCGTTTGTCCTCATGGCCAATGACTTGAGAAACATAGTAACTGCG---------T   3235
CNGC_7-2   CTTCCAATCTCAGATGCAAATGTCAAGTGCCACACAAAGGTAGAGGCGTTTGTCCTCATGGCCAGTGACTTGAGAAACATAGTAACTGAGTGCGGTCGGT   4188
CNGC_7-4   CTTCCTATCTCAGTTGCTAATGTCAGGTGCCACACAAAGGTAGAGGCGTTTGTCCTCATGGCCAATGACTTGAGAAACATTGTAACTACG---------T   5212

CNGC1      GCGGAGATTTGTGGCCGGCTTTTGACTACAAGGCTTCTCAAGATCAGGCGGCTGATCAGATGGCACCGCCGGGGCATTTTCAGCAACAACAAGGGCCAAA   2915
CNGC2      GCGGAGATTTGTGGCCGGCTTTTGACTACAAGGCTTCTCAAGATCAGGCGGCTGATCAGATGGCACCGCCGGGGCATTTTCAGCAACAACAAGGGCCAAA   3335
CNGC_7-2   GCGGAGATTTGTGGCCGGATTTCAAGTATACTGCTTCTCAAGATGAGGAGGCTGATCAGAAGGCACCGCCGGGGCACTTTCAGCAACAACAAGGGCCAAA   4288
CNGC_7-4   GCGGACGTTTGTGGCCGGATTTCGAGTATAATGCTTCTCAAGATGAGGCGGCTGATCAGAAGGCACCGCCGGGGCACTTTCAACAACAACAAGGGCCAAA   5312

CNGC1      GAAGCGTACTGTAGCATCATCAAATATCCATCCTACCGGCTACAGTGGGTGA---------------------------------------------   3015
CNGC2      GAAGCGTACTGTAGCATCATCAAATATCCATCCTACCGGCTACAGTGGGTGA---------------------------------------------   3435
CNGC_7-2   GAAGCG---TGGAACAACATCAAATATCCATCCTACCGGCTACAGTGGGTGATGCATGCGAAGCTGAAGCCTGCTGCAAATCGCCCCTAGCTAGCCTGCA   4385
CNGC_7-4   GAAGAGTACTGGAGCAACATCAAATATCCATCCTACCGGCTACAGTGGGTGATGCATGCGAAGCTGAAGCCTGCTGCAAATCGCCCCTAGCTAGCCTGCA   5412

CNGC1      ----------------------------------------------------------------------------------------------------   3115
CNGC2      ----------------------------------------------------------------------------------------------------   3535
CNGC_7-2   GTGATCTAGGGCGCGCAGTACGTTGTGCATTATTTGCTTTTACATATATAGCAGCTACAGATAAAGCTAGCTAGCTAGCTATCCATCAACTCCATCCTTC   4485
CNGC_7-4   GTGATCTAGGGCGCGCAGTACGTTGTGCATTATTTGCTTTTACATATATAGCAGCTAC------AGCTAGCTAGCTAGCTATCGATCAACTCCATCCTT-   5505

CNGC1      ----------------------------------------------------------------------------------------------------   3215
CNGC2      ----------------------------------------------------------------------------------------------------   3635
CNGC_7-2   AAGCTAGCTGCTACTTACTCTTATTGTCATATATATCATCGTTTCTTCATTTGTATCGCTACGTACTTGAGCTTTGGTACTTGTTATCGATCAGTGATTG   4585
CNGC_7-4   AAGCTAGCTGCTACTTACTCTTATTGTTATATATATCGTCGTTTATTCATTTGTGTCGCTACGTACTTGAGCTTTGGTACTTGTTATCGATCAGTGATTG   5605

CNGC1      ----------------------------------------------------------------------------------------------------   3315
CNGC2      ----------------------------------------------------------------------------------------------------   3735
CNGC_7-2   CCTGTTTTAATATATCAATTATTTGAGA------------------------------------------------------------------------   4685
CNGC_7-4   CCTGTTTTAATATATCAATTATTTGAGAATAAAAATAAGACTTGAGTTTTGATAACGGAGTTGTATGGACAAATTTCTTTCACTCCCACCAGTTCACTCA   5705

CNGC1      ----------------------------------------------------------------------------------------------------   3415
CNGC2      ----------------------------------------------------------------------------------------------------   3835
CNGC_7-2   ----------------------------------------------------------------------------------------------------   4785
CNGC_7-4   AAGATCAACAAGACACTCAAGGAAGGCAATTCCTATGGAGTGGAACAACTTCTACGTTGGGTAGCAGTACCTGACGAACATTTCTCCAAACTTCCTAGCT   5805

CNGC1      ----------------------------------------------------------------------------------------------------   3515
CNGC2      ----------------------------------------------------------------------------------------------------   3935
CNGC_7-2   ----------------------------------------------------------------------------------------------------   4885
CNGC_7-4   TGAACGAGAATGTGAGATCTCATTCAAAAGTGGAAGGCTTTACTCTCATGGCCAGCGACTTGAGAAAGATAGCCAACAACTGCGCAGGCCTTTGGAAGTT   5905

CNGC1      ----------------------------------------------------------------------------------------------------   3615
CNGC2      ----------------------------------------------------------------------------------------------------   4035
CNGC_7-2   ----------------------------------------------------------------------------------------------------   4985
CNGC_7-4   CACCAATGCCCTGATTGCCCTCAATCCTGAAGAAAGAGAGGTGGCGCGGCACGTACTAGTACGTACTACACTACTACGTACCATAGCAGCAGCAGTCAGT   6005

CNGC1      ----------------------------------------------------------------------------------------------------   3715
```

```
CNGC2       ----------------------------------------------------------------------------------------------------  4135
CNGC_7-2    ----------------------------------------------------------------------------------------------------  5085
CNGC_7-4    CGTTTCCGAAATGCAGCAAAGAAGTCGTCCGGAGCAAACCCGTCGGCAGTTGAATTTTGCCTCAGAACTCGCCAATGAACAACGTTTGGCTATAGCTTGT  6105

CNGC1       ----------------------------------------------------------------------------------                    2967
CNGC2       ----------------------------------------------------------------------------------                    3387
CNGC_7-2    ----------------------------------------------------------------------------------                    4613
CNGC_7-4    GTTGCCTTAGCTCGCAACTTGAATTGGTATATATCCATTTTTATTTTACAAGGCAACTAGCTAATAATTAAGACAAAGTCGA                    6187
```

**CDS of WAK**

```
CNGC1       ATGGCTAAACATCAGGGACCAAACAATGAAATCCAAATGAGCATTACACCAGATGTGTCTTATGATTTCCATAGTCCTGAGGCTCTAAAACCGACACGCG   100
CNGC2       ATGGCTAAACATCAGGGACCAACCAATGAAATCCAAATGAGCATTACACCAGATGTGTCTTATGATTTCCATAGTCCTGAGGCTCTAAAACCGACACCCG   100
CNGC_7-2    ----------------------------------------------------------------------------------------------------
CNGC_7-4    ----------------------------------------------------------------------------------------------------

CNGC1       AACATGGACAACCTCAAGCAAGCAGCAAAAGATGGAATTGGCCAGCAGTTGCCGGAGTTTCTTCAGCTTTGAACCCATGGTGGAATAAGATACTGATAAC   200
CNGC2       AACATGGACAACCTCAAGCAAGCAGCAAAAGACGGAATTGGCCAACAGTTGCCGGAGTTTCTTCAGCTTTGAACCCATGGTGGAATAAGATACTGGTAAC   200
CNGC_7-2    ----------------------------------------------------------------------------------------------------
CNGC_7-4    ----------------------------------------------------------------------------------------------------

CNGC1       TTCATGTGTGATTGCAGTTTTTTTTGATCCCTTGTTCTTTTACATTCCATACACCAGCGAGGAAAAAAAGTGTATGGGAAACGACGAAAGAGTGCAGACT   300
CNGC2       TTCATGTGTGATTGCAGTTTTATTTGATCCCTTGTTCTTTTACATTCCATACACCAGCGAGGAAAAAAAGTGTATGGGAAACGACGAAAGAGTGCAGACT   300
CNGC_7-2    ----------------------------------------------------------------------------------------------------
CNGC_7-4    ----------------------------------------------------------------------------------------------------

CNGC1       CCAGCTCTGATTTTCCGATCAGTCACAGACATCATTTTCCTGGTGCATATCATATACCAACTATATGTGGCTATCAAATTTGCACACTCAAGGGTTTCCG   400
CNGC2       CCAGCTCTGATTTTCCGATCAGTCACAGACATCATTTTCCTGGTGCATATCATATACCAACTATATGTGGCTATCAAATTTGCACTCTCAAGGGTTTCCG   400
CNGC_7-2    ----------------------------------------------------------------------------------------------------
CNGC_7-4    ----------------------------------------------------------------------------------------------------

CNGC1       AGGATGAAAGTTTGGTTTTTAGATATGAATCGGTATCCCGAGAGCAAATGATTCAGTTTGCGAAGGCATTTTCTCACAAGTTGTCATGGCGATCTTTCCT   500
CNGC2       AGGATGAAAGTTTGGTTTTTAGATATGAATCGGTATCCCGAGAGCAAATGATTCAGTTTGCGAAGGCATTTTCTCACAAGTTGTCATGGCGATCTTTCCT   500
CNGC_7-2    ----------------------------------------------------------------------------------------------------
CNGC_7-4    ----------------------------------------------------------------------------------------------------

CNGC1       AACTGACGTTTTCGCTATTTTGCCCATACCACAACTTCTAATAGTAGGATTGTTTTTCAACATCGGAGGCAATAACTACTTGTTAAAAAGAAAGATCGTG   600
CNGC2       AACTGACGTTTTCGCTATTTTGCCCATACCACAACTGCTAATAGTAAGATTGTTTTTCAACATTGGAGGCAATAACTACTTGTTAAAAAGAAAGATTGTG   600
CNGC_7-2    ----------------------------------------------------------------------------------------------------
CNGC_7-4    ----------------------------------------------------------------------------------------------------

CNGC1       GGCGGTTTTCTTCTAATCCAATATGTGCCAAGAATCTATCGAATTTATCTTTCATCTGTCGCGGTCACAGAATCTGCATTGTGGGTTAGAGGTGTATTTA   700
CNGC2       AGCGGTTTTCTTCTAATCCAATATGTGCCAAGAATCTATCGAATTTATCTTTCATCTGTCGCGGTCACAGAATCTGCAATGTGGGTTAGAGGTGTATTTA   700
CNGC_7-2    ----------------------------------------------------------------------------------------------------
CNGC_7-4    ----------------------------------------------------------------------------------------------------

CNGC1       ACTTCTTCCTCTACATTCTTGCGAGTCATGCACTTGGAGCCTTTTGGTACTTTTGTTCTATTCAGCGAGAGACATCATGTTGGTATCGAACTTGCGTCGC   800
CNGC2       ACATTTTTCTCTACATTCTTGCGAGTCATGCACTTGGAGCCTTTTGGTACTTTTGTTCTATTCAGCGAGAGACATTATGTTGGTATCGAACTTGCCTCGC   800
CNGC_7-2    ----------------------------------------------------------------------------------------------------
CNGC_7-4    ----------------------------------------------------------------------------------------------------

CNGC1       TACAAATATATCAGAATCAAAATGCACCTTTTACTGCCATAACAGTACTGCTGCACTTATAACCCCACAATTCAAGACAACTCTAGACGAGAAATGCATT   900
CNGC2       TACAAATATATCAGAATCAAAATGCACCTTTTACTGCCATAACAGTGCTGCTGCACTTATGACCCCACAATTCAAGACAACTCTAGACGAGAAATGCATT   900
CNGC_7-2    ----------------------------------------------------------------------------------------------------
CNGC_7-4    ----------------------------------------------------------------------------------------------------

CNGC1       CTAAAAGTTCCATATAACATGACCGATCCACCATTTGATTTTGGAATATTTTTTGATGCCCTCAAAAAATGATATCCAAGGGAAAATAAATGTTCCACAAA  1000
CNGC2       CTAAAAGTTCCATATAACATGACCGATCCACCATATGATTTTGGAATATTTTTTGATGCCCTCAAAAAATGATATCCAAGGGAAAATAAATGTTCCACAAA  1000
CNGC_7-2    ----------------------------------------------------------------------------------------------------
CNGC_7-4    ----------------------------------------------------------------------------------------------------

CNGC1       AGATTGGTTACTGTTTTTGGTGGGGCTTGCGAAACTTGAGTAATTTTGGTACAAGTTTGGAAACAAGTACCTATCTGTGGGAAAACAGCTTTGCCATTTT  1100
CNGC2       AGATTGGTTACTGTTTTTGGTGGGGCTTGCGAAACTTGAGTAATTTTGGTACAAATTTGGAAACAAGTACCTATCTGTGGGAAAACAGCTTTGCCATTTT  1100
CNGC_7-2    ----------------------------------------------------------------------------------------------------
CNGC_7-4    ----------------------------------------------------------------------------------------------------

CNGC1       AATTTCTATCATCGGCTTGCTGTTGTTTTTATACCTCATTGGAAATGTACAGATTTATATGCAGAGGGCAACTACCAGAGCGGAGGAGGTAAGAGAGAAG  1200
CNGC2       AATTTCTATCATCGGCTTGCTGTTGTTTTTATACCTCATTGGAAATGTACAGATTTATATGCAGAGGGCAGTTACCAGAGCGGAGGAGGTAAGAGAGAAG  1200
CNGC_7-2    ----------------------------------------------------------ATGGAGAGGGCAACTACCAGAGCGGATGAGGTAAGAGAGAAG    42
CNGC_7-4    ----------------------------------------------------------ATGGAGAGGGCAACTACCAGAGCGGAGGAGGTAAGAGAGAAG    42

CNGC1       CTTCGGATCAAAAAGAACGATATAGATAATTGGGTCCAAAATAATGGTATCGAGAAGGACATGAAGGAAGAGATCATGAAAAACATCGTTGAAAAGTTGG  1300
CNGC2       CTTCGGATCAAAAAGAACGATATAGATAATTGGATCCAAAATAATGGTATCGAGAATTGACATGAAGGAAGAGATCATGAAAAACATCGTTGAAAAGTTGG  1300
CNGC_7-2    CTTCGGATCAAAAAGAACGATATAGATGATTGGATCGAAAATAATGGTATCGGGAAGGACATGAAGGAAGAGATCATGAAAAACATCAAGGAGAAGTTGG   142
CNGC_7-4    CTTCGGATAAAACAGAACGATATAGATAATTGGATCGAAAATAATGGTCTCGAGAAGGACATGAAGGAAGAGATCATGAAAAACATCAATGAGAAGTTGG   142

CNGC1       AAAACGACATAAGTGCAGATCTGGATGATATATACTCTATTCTTCCCCCGTATACCAGGAAAGTTGTGAAGCGATTTGTCGGCATGAGGACGCTAAGAAA  1400
CNGC2       AAAACGACATAAGTGCAGATCTGGATGATATATACTCTATTCTTCCCCCGTATACCAGGAAAGTTGTGAAGCGATTTGTCGGCATGAGGACGCTAAGAAA  1400
CNGC_7-2    AAAAGGACATAAGTGCAGATCTGGATGATATATACTCTATTCTTCCCCCGAATACCAGGAAAGTTGTGAAGCGATTTGTCGGCATGAGGACGCTAAGAAA   242
CNGC_7-4    AAAAGGACATAAATGCAGATCTGGATGATATATACTCTATTCTTCTCCCGTATACCAGGAAAGTTGTGAAGCGATTTGTCGGCATGAGGACGCTAAGAAA   242

CNGC1       TGTACCAATGCTTAGCCAGGTGGATGGAAGAGTTTTGAAGATGATGTGCGACTATCTGAAGCCGGTCAAGTACCAGGAGAATACAATGGTTTTTGAAACG  1500
CNGC2       TGTACCAATGCTTAGCCAGGTGGATGGAAGAGTTTTGAAGATGATGTGCGACTATCTGAAGCCGGTCAAGTACCAGGAGAATACAATGGTTTTTGAAACG  1500
```

```
CNGC_7-2  TGTACCAATGCTTAGCGAGGTGGATGGAAGAGTTTTGAAGATGATGTGCGACTATCTGAAGCCGGTCAAGTTCCAGGAGAATACAATGGTTTTAAAAATG   342
CNGC_7-4  TGTACCAATGCTTAGCCAGGTGGATGGAAGAGTTTTGAAGATGATGTGCGACTATCTGAAACCGGTCAAGTACCAGGAGAATACAATGGTTTTTAAAATG   342

CNGC1     GGAGTACCATTTGATAGAATGGTCTTCATTACAGATGGGTCAATGTGGACTTACACGATTGCTACTGATCATCAGACTAGTCATTCTGGAAAAGCAACGT  1600
CNGC2     GGAGTACCATTTGATAGAATGGTCTTCATTACAGATGGGTCAATGTGGACTTACACGATTGCTACTGATCATCAGACTAGTCATTCTGGAAAAGCAACGT  1600
CNGC_7-2  GGAAAACCATTTGATAGAATGGTCTTCATTACAGATGGGTCAATGTGGACTTAC---------ACTGATCATCAGACTAGTGATTCTGGAAAAGCAACGT   433
CNGC_7-4  GGAGTACCATTTGATAGAATGGTCTTCATTACAGATGGGTCAATGTGGACTTACACGACTGCTACTGATCATCAGACTAGTCATTCTGGAAAAGCAACGT   442

CNGC1     CAGGAGGACTAGTTACTTCTTCGTCCACGGAAGCTCACTTCCTCAAGAAGGGTGATGCTTATGGACACATGCTTCTGCAATTAGCATCATCATCTTCTAC  1700
CNGC2     CAGGAGGACTAGTTACTTCTTCGTCCACGGAAGCTCACTTCCTCAAGAAGGGTGATGCTTATGGACACATGCTTCTGCAATTAGCATCATCATCTTCTAC  1700
CNGC_7-2  CAGGAGGACTAGTTACTTCTTCGTCCACGGAAGCTCACTTCCTCGAGAAGGGTGATGCTTATGGACACATGCTTCTGCAATTAGCATCATCATCTTTTAC   533
CNGC_7-4  CAGGAGGACTAGTTACTTCTTCGTCCACGGAAGCTCACCTCCTCGAGAAGGGTGATGCTTATGGACACATGCTTCTGCAATTAGCATCATCATCTTTTAC   542

CNGC1     GGCGGTTCCTATCTCAGTTGCAAATGTCAGGTGCCACACAAAGGTAGAGGCGTTTGTCCTCATGGCCAATGACTTGAGAAACATAGTAACTGCGTGCGG-  1799
CNGC2     GGCGGTTCCTATCTCAGTTGCAAATGTCAGGTGCCACACAAAGGTAGAGGCGTTTGTCCTCATGGCCAATGACTTGAGAAACATAGTAACTGCGTGCGG-  1799
CNGC_7-2  GACGCTTCCAATCTCAGATGCAAATGTCAAGTGCCACACAAAGGTAGAGGCGTTTGTCCTCATGGCCAGTGACTTGAGAAACATAGTAACTGAGTGCGGT   633
CNGC_7-4  GGCGCTTCCTATCTCAGTTGCTAATGTCAGGTGCCACACAAAGGTAGAGGCGTTTGTCCTCATGGCCAATGACTTGAGAAACATTGTAACTACGTGCGG-   641

CNGC1     --------AGATTTGTGGCCGGCTTTTGACTACAAGGCTTCTCAAGATCAGGCGGCTGATCAGATGGCACCGCCGGGGCATTTTCAGCAACAACAAGGGC  1891
CNGC2     --------AGATTTGTGGCCGGCTTTTGACTACAAGGCTTCTCAAGATCAGGCGGCTGATCAGATGGCACCGCCGGGGCATTTTCAGCAACAACAAGGGC  1891
CNGC_7-2  CGGTGCGGAGATTTGTGGCCGGATTTCAAGTATACTGCTTCTCAAGATGAGGAGGCTGATCAGAAGGCACCGCCGGGGCACTTTCAGCAACAACAAGGGC   733
CNGC_7-4  --------ACGTTTGTGGCCGGATTTCGAGTATAATGCTTCTCAAGATGAGGCGGCTGATCAGAAGGCACCGCCGGGGCACTTTCAACAACAACAAGGGC   733

CNGC1     CAAAGAAGCGTACTGTAGCATCATCAAATATCCATCCTACCGGCTACAGT--------------------------------------------------  1941
CNGC2     CAAAGAAGCGTACTGTAGCATCATCAAATATCCATCCTACCGGCTACAGT--------------------------------------------------  1941
CNGC_7-2  CAAAGAAGCG---TGGAACAACATCAAATATCCATCCTACCGGCTACAGT--------------------------------------------------   780
CNGC_7-4  CAAAGAAGAGTACTGGAGCAACATCAAATATCCATCCTACCGGCTACATTCACTCAAAGATCAACAAGACACTCAAGGAAGGCAATTCCTATGGAGTGGA   833

CNGC1     ----------------------------------------------------------------------------------------------------  1941
CNGC2     ----------------------------------------------------------------------------------------------------  1941
CNGC_7-2  ----------------------------------------------------------------------------------------------------   780
CNGC_7-4  ACAACTTCTACGTTGGGTAGCAGTACCTGACGAACATTTCTCCAAACTTCCTAGCTTGAACGAGAATGTGAGATCTCATTCAAAAGTGGAAGGCTTTACT   933

CNGC1     ----------------------------------------------------------------------------------------------------  1941
CNGC2     ----------------------------------------------------------------------------------------------------  1941
CNGC_7-2  ----------------------------------------------------------------------------------------------------   780
CNGC_7-4  CTCATGGCCAGCGACTTGAGAAAGATAGCCAACAACTGCGCAGGCCTTTGGAAGTTCACCAATGCCCTGATTGCCCTCAATCCTGAAGAAAGAGAGGTGG  1033

CNGC1     ----------------------------------------------------------------------------------------------------  1941
CNGC2     ----------------------------------------------------------------------------------------------------  1941
CNGC_7-2  ----------------------------------------------------------------------------------------------------   780
CNGC_7-4  CGCGGCACGTACTAGTACGTACTACACTACTACGTACCATAGCAGCAGCAGTCAGTCGTTTCCGAAATGCAGCAAAGAAGTCGTCCGGAGCAAACCCGTC  1133

CNGC1     ----------------------------GGGTGA                                                              1947
CNGC2     ----------------------------GGGTGA                                                              1947
CNGC_7-2  ----------------------------GGGTGA                                                               786
CNGC_7-4  GGCAGTTGAATTTTGCCTCAGAACTCGCCAATGA                                                              1167
```

**Peptide sequence of WAK**

```
CNGC1     MAKHQGPNNEIQMSITPDVSYDFHSPEALKPTREHGQPQASSKRWNWPAVAGVSSALNPWWNKILITSCVIAVFFDPLFFYIPYTSEEKKCMGNDERVQT  100
CNGC2     MAKHQGPTNEIQMSITPDVSYDFHSPEALKPTPEHGQPQASSKRRNWPTVAGVSSALNPWWNKILVTSCVIAVLFDPLFFYIPYTSEEKKCMGNDERVQT  100
CNGC_7-2  ----------------------------------------------------------------------------------------------------
CNGC_7-4  ----------------------------------------------------------------------------------------------------

CNGC1     PALIFRSVTDIIFLVHIIYQLYVAIKFAHSRVSEDESLVFRYESVSREQMIQFAKAFSHKLSWRSFLTDVFAILPIPQLLIVGLFFNIGGNNYLLKRKIV  200
CNGC2     PALIFRSVTDIIFLVHIIYQLYVAIKFALSRVSEDESLVFRYESVSREQMIQFAKAFSHKLSWRSFLTDVFAILPIPQLLIVRLFFNIGGNNYLLKRKIV  200
CNGC_7-2  ----------------------------------------------------------------------------------------------------
CNGC_7-4  ----------------------------------------------------------------------------------------------------

CNGC1     GGFLLIQYVPRIYRIYLSSVAVTESALWVRGVFNFFLYILASHALGAFWYFCSIQRETSCWYRTCVATNISESKCTFYCHNSTAALITPQFKTTLDEKCI  300
CNGC2     SGFLLIQYVPRIYRIYLSSVAVTESAMWVRGVFNIFLYILASHALGAFWYFCSIQRETLCWYRTCLATNISESKCTFYCHNSAAALMTPQFKTTLDEKCI  300
CNGC_7-2  ----------------------------------------------------------------------------------------------------
CNGC_7-4  ----------------------------------------------------------------------------------------------------

CNGC1     LKVPYNMTDPPFDFGIFFDALKNDIQGKINVPQKIGYCFWWGLRNLSNFGTSLETSTYLWENSFAILISIIGLLLFLYLIGNVQIYMQRATTRAEEVREK  400
CNGC2     LKVPYNMTDPPYDFGIFFDALKNDIQGKINVPQKIGYCFWWGLRNLSNFGTNLETSTYLWENSFAILISIIGLLLFLYLIGNVQIYMQRAVTRAEEVREK  400
CNGC_7-2  --------------------------------------------------------------------------------------MERATTRADEVREK   14
CNGC_7-4  --------------------------------------------------------------------------------------MERATTRAEEVREK   14

CNGC1     LRIKKNDIDNWVQNNGIEKDMKEEIMKNIVEKLENDISADLDDIYSILPPYTRKVVKRFVGMRTLRNVPMLSQVDGRVLKMMCDYLKPVKYQENTMVFET  500
CNGC2     LRIKKNDIDNWIQNNGIENDMKEEIMKNIVEKLENDISADLDDIYSILPPYTRKVVKRFVGMRTLRNVPMLSQVDGRVLKMMCDYLKPVKYQENTMVFET  500
CNGC_7-2  LRIKKNDIDDWIENNGIGKDMKEEIMKNIKEKLEKDISADLDDIYSILPPNTRKVVKRFVGMRTLRNVPMLSEVDGRVLKMMCDYLKPVKFQENTMVLKM  114
CNGC_7-4  LRIKQNDIDNWIENNGLEKDMKEEIMKNINEKLEKDINADLDDIYSILLPYTRKVVKRFVGMRTLRNVPMLSQVDGRVLKMMCDYLKPVKYQENTMVFKM  114

CNGC1     GVPFDRMVFITDGSMWTYTIATDHQTSHSGKATSGGLVTSSSTEAHFLKKGDAYGHMLLQLASSSSTAVPISVANVRCHTKVEAFVLMANDLRNIVTACG  600
CNGC2     GVPFDRMVFITDGSMWTYTIATDHQTSHSGKATSGGLVTSSSTEAHFLKKGDAYGHMLLQLASSSSTAVPISVANVRCHTKVEAFVLMANDLRNIVTACG  600
CNGC_7-2  GKPFDRMVFITDGSMWTY---TDHQTSDSGKATSGGLVTSSSTEAHFLEKGDAYGHMLLQLASSSFTTLPISDANVKCHTKVEAFVLMASDLRNIVTECG  211
CNGC_7-4  GVPFDRMVFITDGSMWTYTTATDHQTSHSGKATSGGLVTSSSTEAHLLEKGDAYGHMLLQLASSSFTALPISVANVRCHTKVEAFVLMANDLRNIVTTCG  214

CNGC1     D---LWPAFDYKASQDQAADQMAPPGHFQQQQGPKKRTVASSNIHPTGY---------------------------------------------------  646
CNGC2     D---LWPAFDYKASQDQAADQMAPPGHFQQQQGPKKRTVASSNIHPTGY---------------------------------------------------  646
```

```
CNGC_7-2  RCGDLWPDFKYTASQDEEADQKAPPGHFQQQQGPKKR-GTTSNIHPTGY---------------------------------------------------  259
CNGC_7-4  R---LWPDFEYNASQDEAADQKAPPGHFQQQQGPKKSTGATSNIHPTGYIHSKINKTLKEGNSYGVEQLLRWVAVPDEHFSKLPSLNENVRSHSKVEGFT  311

CNGC1     -------------------------------------------------------------SG*-------------                      649
CNGC2     -------------------------------------------------------------SG*-------------                      649
CNGC_7-2  -------------------------------------------------------------SG*-------------                      262
CNGC_7-4  LMASDLRKIANNCAGLWKFTNALIALNPEEREVARHVLVRTTLLRTIAAAVSRFRNAAKKSSGANPSAVEFCLRTQ*                      389
```

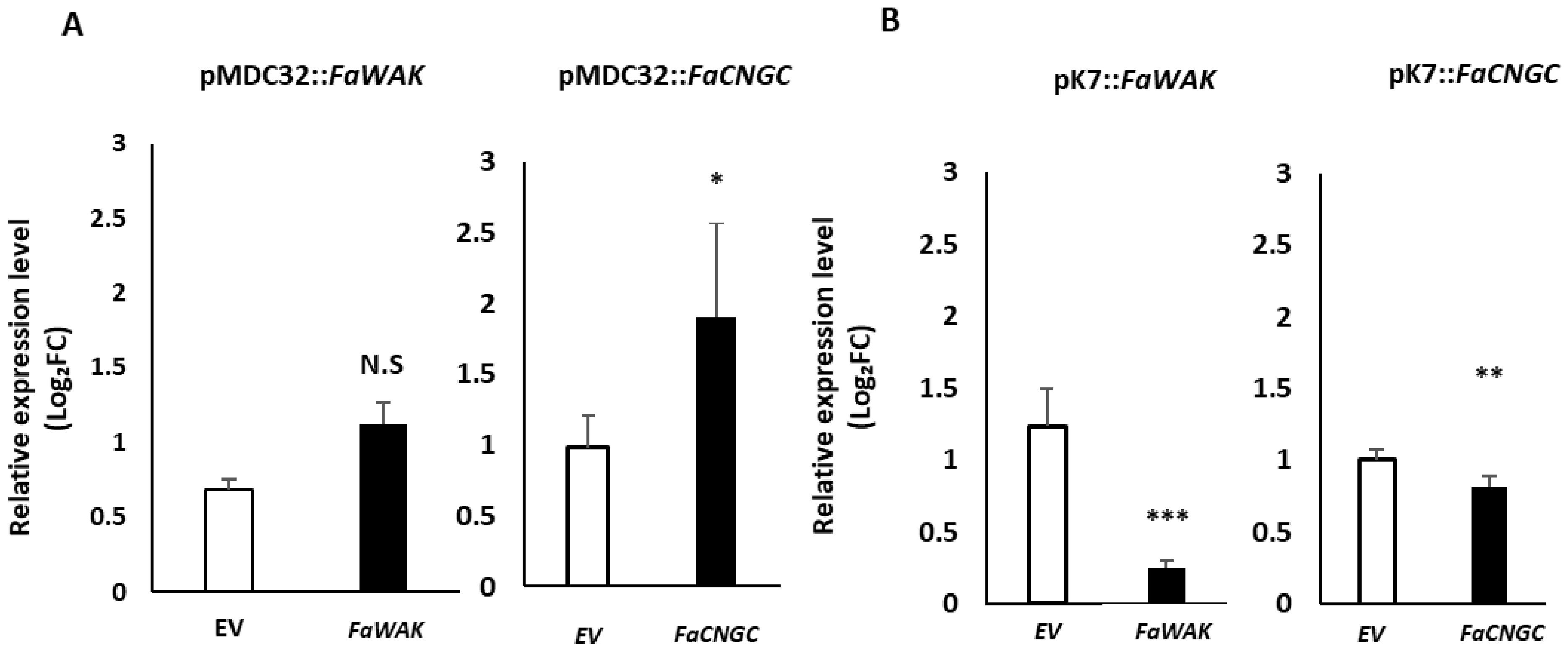


**Supplementary Fig. S11 Gene expression analysis of *FaWAK* and *FaCNGC* by qRT-PCR in strawberry resistance to *P. cactorum* pathogens**. (A) The transient expression assay was conducted employing the pMDC32 overexpression vector to drive the expression of *FaWAK* and *FaCNGC* genes. (B) The transient expression assay was conducted employing the pK7GWIWG(II) RNAi vector to drive the knockdown expression of *FaWAK* and *FaCNGC* genes. The x-axis represents the vector composition used. EV denotes 'Empty Vector, 'Error bars represent standard deviation. Asterisks indicate significant outcomes with P values of Student's t-test (*P < 0.05, **P < 0.01, ***P < 0.001, Student's t-test), respectively.

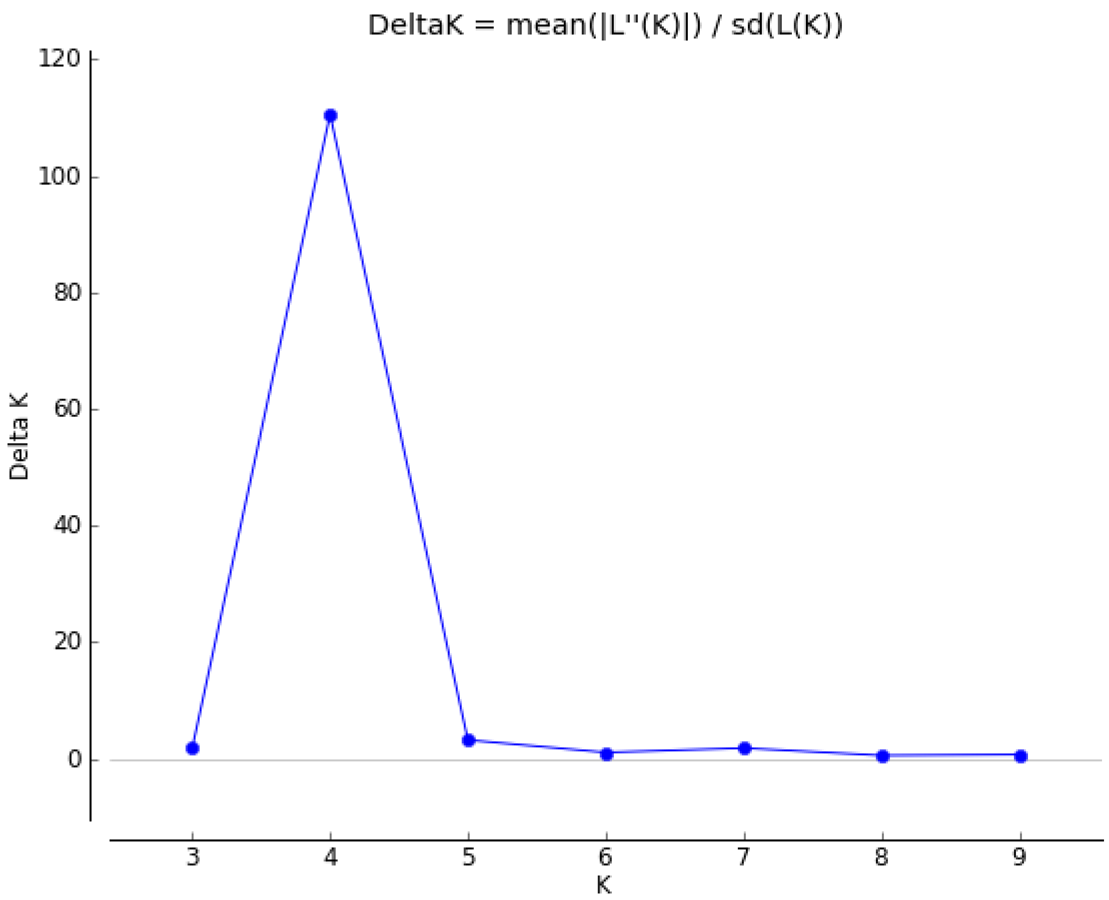


**Supplementary Fig. S12. Delta K values for STRUCTURE analysis of octoploid strawberries.**

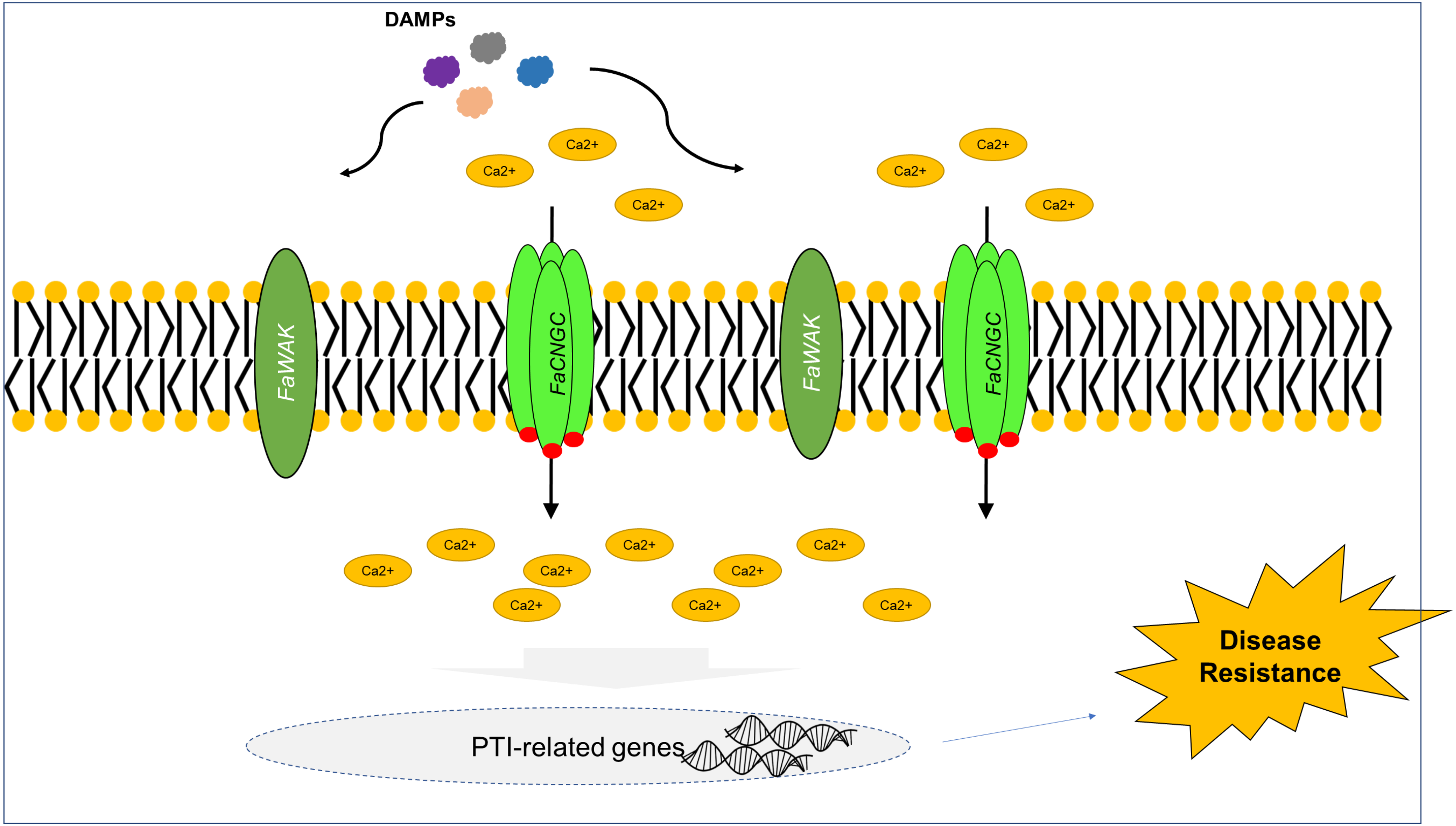


**Supplementary Fig. S13. A hypothetical model for the enhanced resistance against *Phytophthora cactorum* in the *FaWAK, FaCNGC1* knockdown strawberry root.** Differentially expressed genes identified in *FaWAK* and *FaCNGC1* knockdown crown in response to *P. cactorum* were represented. DAMPs such as Oligogalacturonides, and pectin fractions are sensed by *FaWAKs*, leading to the activation of *FaRLCK*. Subsequently, the activated *FaRLCK* phosphorylates *FaCNGC* induces extracellular $Ca^{2+}$ influx and then triggers PTI-related gene expressions, which result in disease resistance in strawberry.
